# Serum profiles of metabolomic and GDF-15 differentiate idiopathic pulmonary fibrosis from other interstitial lung diseases: a preliminary study

**DOI:** 10.1101/2022.03.22.484740

**Authors:** Meng Li, Hongyan Ren, Lifan Liu, Wenting Li, Ziqi Wang, Jingjing Zhang, Xiaoli Wang, Yiping Hu, Kaixuan Zang, Yunxia An, Zhiwei Xu, Zhiping Guo, Ulrich Costabel, Huaping Dai, Xiaoju Zhang, Zheng Wang

## Abstract

**Objective:** 1.Explore the differential metabolites and metabolic pathways between different interstitial lung disease (ILD) and healthy controls, so as to find out new biomarkers and metabolic pathways of ILD and provide new biological targets for IPF diagnosis and treatment.2.Detect the levels of serum related fibrosis and metabolic indexes of ILD, explore the relationship between these indexes and the types and general parameters of ILD diseases, and focus on GDF-15 on the basis of metabonomics, explore new metabolic pathways related to ILD, and further expound the pathogenesis of idiopathic pulmonary fibrosis.

**Method:** 1.According to the inclusion criteria and exclusion criteria, the ILD patients admitted to our hospital from January 1, 2021 to January 1, 2022 were signed with informed consent, and all of them had clear multidisciplinary diagnosis. At the same time, the physical examination in the health management center of our hospital was selected as the healthy control group. General clinical data of ILD patients and healthy controls were collected, including gender, age, onset, clinical manifestations, clinical biochemistry, lung function and other related data. Collect patients’ peripheral venous blood, keep serum samples, and analyze the serum of ILD patients and healthy controls by liquid chromatography-mass spectrometry (LC-MS) to obtain differential metabolites, metabolic maps and related data. According to the diagnosis, the patients were divided into Idiopathic pulmonary fibrosis (IPF), connective tissue disease-associated interstitial lung disease (CTD-ILD), other interstitial lung disease (ILD) patients and healthy control group. R statistical software was used to observe whether there are differential metabolites in IPF group and healthy control group, CTD-ILD group and healthy control group, other ILD groups and healthy control group, IPF group and CTD-ILD group, IPF group and other ILD groups, CTD-ILD group and other ILD groups by using positive and negative ion flow charts and score charts of orthogonal partial least squares-discriminant analysis. And the potential biomarkers were found through the S-plot diagram of OPLS-DA and the variable weight value VIP>1. ROC curve analysis of differential metabolites was carried out by SPSS21.0 software to analyze the value of differential diagnosis and find out new biomarkers with differential diagnosis significance. Based on KEGG database, the metabolic pathway enrichment analysis of differential metabolites was carried out to find out the main metabolic pathways. 2.Serum levels of type I collagen, KL-6, IL-1β, TNF-α, IGF-1 and GDF-15 were detected by Enzyme linked immunosorbent assay (ELISA) in all patients and healthy controls. Using SPSS21.0 statistical software and R statistical software, using T-test, variance analysis or nonparametric test method, according to different diagnoses (IPF/CTD-ILD/ other ILD/ healthy control group, ILD/ healthy control group, IPF/ other), the data were analyzed and compared in subgroups. Pearson correlation analysis was used to analyze the correlation between serum biomarkers and other factors such as lung function, metabolomics and so on. ROC curve was used to analyze the diagnostic efficiency of each index, and the best critical value, sensitivity and specificity were found out.

**Results:** 1.Analysis of general clinical data and clinical indicators: 26 cases of IPF, 21cases of CTD-ILD, 23 cases of others ILD and 20 cases of healthy control group were included, respectively. There were significant differences in age, sex and smoking history among the four groups (P < 0.05), and the differences were statistically significant. There were statistically significant differences among IPF group, CTD-ILD group and other ILD groups in cough and expectoration symptoms, eosinophil count percentage, creatine kinase, creatine kinase isoenzyme, C-reactive protein and carbon monoxide diffusion capacity (DLco) (P < 0.05). 2. Metabonomics analysis: LC-MC total ion flow chart of serum samples and orthogonal partial least squares discriminant analysis (OPLS-DA) of serum samples showed the difference of ion peak intensity, and there were different metabolites among each group. The OPLS-DA model has good quality evaluation and reliable data analysis. There are 193, 115, 101, 188, 148 and 169 differential metabolites in IPF group and healthy control group, IPF group and CTD-ILD group, CTD-ILD group and other ILD groups, others ILD groups and healthy control group respectively. There is a correlation between the differential metabolites of each ILD subgroup and the healthy control group. The most relevant metabolic pathways in IPF mainly include choline metabolic pathway, metabolic pathway and linoleic acid metabolic pathway in cancer. The metabolic pathways most related to CTD-ILD and other ILD include choline metabolic pathway and retrograde neural signal metabolic pathway in cancer. Metabolic pathway and linoleic acid metabolic pathway may distinguish IPF from other ILD. Metabolites involved in metabolic pathway are L-Glutamate, PE(18:3(6Z,9Z,12Z)/P-18:0) and PC (18: 2 (9z, 12z)/20: 2 (11z, 14z)), respectively. There are three differential metabolites of linoleic acid pathway, namely 12,13-EpOME, 13S-HODE and PC(18:2(9Z,12Z)/20:2(11Z,14Z)). There are 35 kinds of differential metabolites in IPF group and healthy control group, and the top three places of AUC are 3- sulfodeoxycholic acid, 20,22-dihydrodigitalis glycoside, (±14) (15)-EET-SI, with AUC values of 0.985, sensitivity of 100% and specificity of 92.3%. L- acetylcarnitine, 5-hydroxydodecanoate and L-Glutamate are involved in significant enrichment pathways, among which the ROC curve area of L-Glutamate is the largest, with AUC value of 0.921, cutoff value of 40145.490, sensitivity of 80%, specificity of 96%, and participation in metabolic pathways. 3. Serum metabolism-related detection indexes: The six serum detection indexes of IPF group and CTD-ILD group are higher than those of healthy control group, and the difference is statistically significant (P < 0.05), but the difference between IPF group and CTD-ILD group is not statistically significant (P>0.05). The expression of GDF-15 in IPF group and non-IPF ILD group was higher than that in healthy control group, with significant difference, but there was no significant difference between IPF group and non-IPF ILD group. Type I collagen and IL-1β were negatively correlated with DLco. KL-6 is negatively correlated with the measured values of VC and DLco; IGF-1 was negatively correlated with FVC, VC and DLco (all P<0.05). GDF-15 was positively correlated with type I collagen, KL-6, IL-1β, TNF-α and IGF-1, and the difference was statistically significant (P < 0.05). ROC curve shows that the area under ROC curve (AUC) of six serum indexes in IPF group is greater than 0.9; In CTD-ILD group, the AUC of IGF-1 was greater than 0.9, the cutoff value was 35.081ng/ml, the sensitivity was 90%, the specificity was 76%, and the area under ROC curve was greater than 0.8, and the difference was statistically significant. In other ILD groups, the ROC curve area of these six indicators ranged from 0.6 to 0.8, among which IGF-1, GDF-15 and TNF-α had diagnostic value. In IPF, L-Glutamate was positively correlated with type I collagen, KL-6, IGF-1, IL-1β, TNF-α and GDF-15, and the difference was statistically significant (P < 0.05), with the highest correlation with IGF-1.

**Conclusion:** In this study, the serum samples of IPF patients, CTD-ILD patients, other interstitial lung diseases and healthy controls were analyzed by LM-MS and ELISA, and the following conclusions were drawn. 1.There are differences in serum metabolites in different groups. IPF has three main metabolic pathways, namely choline metabolic pathway, metabolic pathway and linoleic acid metabolic pathway in cancer, among which metabolic pathway and linoleic acid metabolic pathway are different from other groups. There are 35 differential metabolites in IPF and healthy control group with the area under ROC curve larger than 0.80, among which the metabolites with the highest diagnostic value are 3- sulfodeoxycholic acid, 20,22-dihydrodigitalis glycoside and (±) 14 (15)-EET-SI. The AUC value of L- glutamic acid is the highest among the metabolites that participate in the enrichment pathway significantly. 2. The expressions of type I collagen, KL-6, IL-1β, TNF-α, IGF-1 and GDF-15 are different in different ILDs, which have potential diagnostic and differential diagnostic value, and are correlated with lung function indexes. In IPF, GDF-15 is positively correlated with L-Glutamate, which may be involved in metabolic pathway to further promote pulmonary fibrosis.

## Introduction

Interstitial lung disease is a non-neoplastic disease caused by different degrees of inflammatory cell infiltration and fibrosis, which mainly shows diffuse lung lesions, involving alveolar walls and surrounding tissues, mainly in fibrous tissues, lymphatic vessels and blood vessels of interstitial lung, and often involving lung parenchyma and alveolar epithelial cells. There are nearly 300 kinds of ILD diseases, many of which have unclear pathogenesis, and their clinical manifestations, treatment and prognosis are different. The pulmonary function of ILD is mainly characterized by restrictive and diffuse dysfunction, eventually leading to hypoxemia and respiratory failure. IPF is a type with poor prognosis in ILD, which is rare in clinic. Its incidence rate is about 2 ∼ 29/100,000, and it is on the rise in recent years. IPF mainly occurs in middle-aged and elderly people. Pulmonary tissue and HRCT show common interstitial pneumonia (UIP), with a median survival period of 3-5 years, which is one of the important diseases affecting human health [1]. The correct diagnosis of ILD is of great significance to the treatment and prognosis. At present, the evaluation of ILD’s condition and therapeutic effect depends on comprehensive indicators, mainly including clinical symptoms, physiological functions and HRCT. Therefore, ideal biomarkers that can be used for diagnosis, differential diagnosis, evaluation of illness and curative effect are still of great significance and one of the research hotspots at present [2]. Metabolomics or metabonomics is a branch of biology after genomics, transcriptomics and proteomics. Metabonomics is a technology to study the metabolic pathway of biological system through the change of metabolites or the change of metabolites with time after organisms are stimulated [3]. The detected biological samples can include urine, serum, plasma, bronchoalveolar lavage fluid, etc. The research objects are mainly endogenous small molecules with molecular weight less than 1K, including amino acids, peptides, carbohydrates, lipids and so on. Compared with the traditional metabolic research, metabolomics combines the knowledge of physics, biology and other disciplines, uses modern advanced instrument combined analysis technology to detect the changes of the whole metabolic product spectrum under certain conditions, which has the advantages of high speed and less consumption of samples and solvents, and studies the whole biological function by special multivariate statistical analysis methods [4]. Metabonomics is based on the overall metabolism of organisms, mainly using gas chromatography-mass spectrometry (GC-MS), nuclear magnetic resonance and other technologies to qualitatively and quantitatively analyze metabolites in patients’ body fluids and screen and diagnose diseases, so as to reflect the responses of organisms to pathological stimuli [5], explore new biomarkers, and also reflect metabolic pathways, and further improve the diagnosis rate and treatment scheme of diseases. Metabonomics is now widely used in many research fields, such as drug development, biomarker detection, plant and microorganism related fields [6]. Many biomarkers reflect the process of alveolar epithelial injury and fibrosis, and may be used as biomarkers of ILD. Type I collagen is mainly distributed in skin, tendon and other tissues. In the study of pulmonary fibrosis, type I collagen accounts for 80%, which is an important index to reflect fibrosis. Salivary sugar chain antigen −6(Krebs von den Lungen-6, KL-6) is classified as one of the epithelial mucin 1(MCU1) of Cluster9, which is a high molecular weight glycoprotein expressed on type II lung cells and bronchial epithelial cells. It was first discovered and put forward by Japanese scholars in 1985 [7]. It shows that the study is of great significance in the diagnosis and prognosis of interstitial lung disease as ILD, and it is related to fibrosis. Different types of interstitial lung diseases have different degrees of increase, and can also reflect lung injury and injury degree [8, 9]. −1β(Interleukin-1β (IL-1β), tumor necrosis factor-α (TNF-α) and insulin-like growth factors-1 (IGF-1) are important fibrogenic factors [9, 10]. Growth differentiation factor-15 (GDF15) is a branch member of transforming growth factor −β(TGF-β) superfamily, which is widely expressed under normal physiological conditions. The expression of GDF-15 is increased in many pathological conditions, and it is used as a marker of cell stress. In the animal model of type II alveolar epithelial cell-specific telomere dysfunction, GDF-15 is the most significantly up-regulated protein in aging type II alveolar epithelial cells, so lung epithelial cells are identified as the main source of GDF-15 production in human lungs [11]; Growth factor 15 (GDF-15) is a stress-responsive cytokine, which mediates anorexia and cachexia in many chronic diseases and cancers, and has a great relationship with metabolism [12]. Recent studies show that the increase of GDF-15 level in blood may precede the development of pulmonary fibrosis and may mediate the relationship between aging and interstitial lung abnormalities, but further research is needed [13]. Studies have suggested that GDF-15 may promote pulmonary fibrosis by activating macrophages and fibroblasts [14]. There is no clear research on whether there is a relationship between the types of interstitial lung diseases, and there is no research to explore the metabolic pathway of GDF-15 based on metabonomics. In this study, the serum metabolomics of ILD patients and healthy controls were analyzed by liquid chromatography-mass spectrometry (LC-MS) to screen out the differential metabolites, explore new metabolic pathways and further elaborate the pathogenesis of IPF. At the same time, the biomarkers I collagen, KL-6, IL-1β, TNF-α, IGF-1 and GDF-15 were detected to explore the diagnostic efficiency of these biomarkers and interstitial lung diseases, and whether they can be used as discriminators. At the same time, we focused on GDF-15 based on metabonomics, explored new metabolic pathways, further explored the metabolic regulation mechanism of ILD, and explored possible biomarkers.

## The first chapter is the study of serum metabonomics of ILD patients based on liquid chromatography-mass spectrometry

### 1 research materials and methods

#### 1.1 experimental process

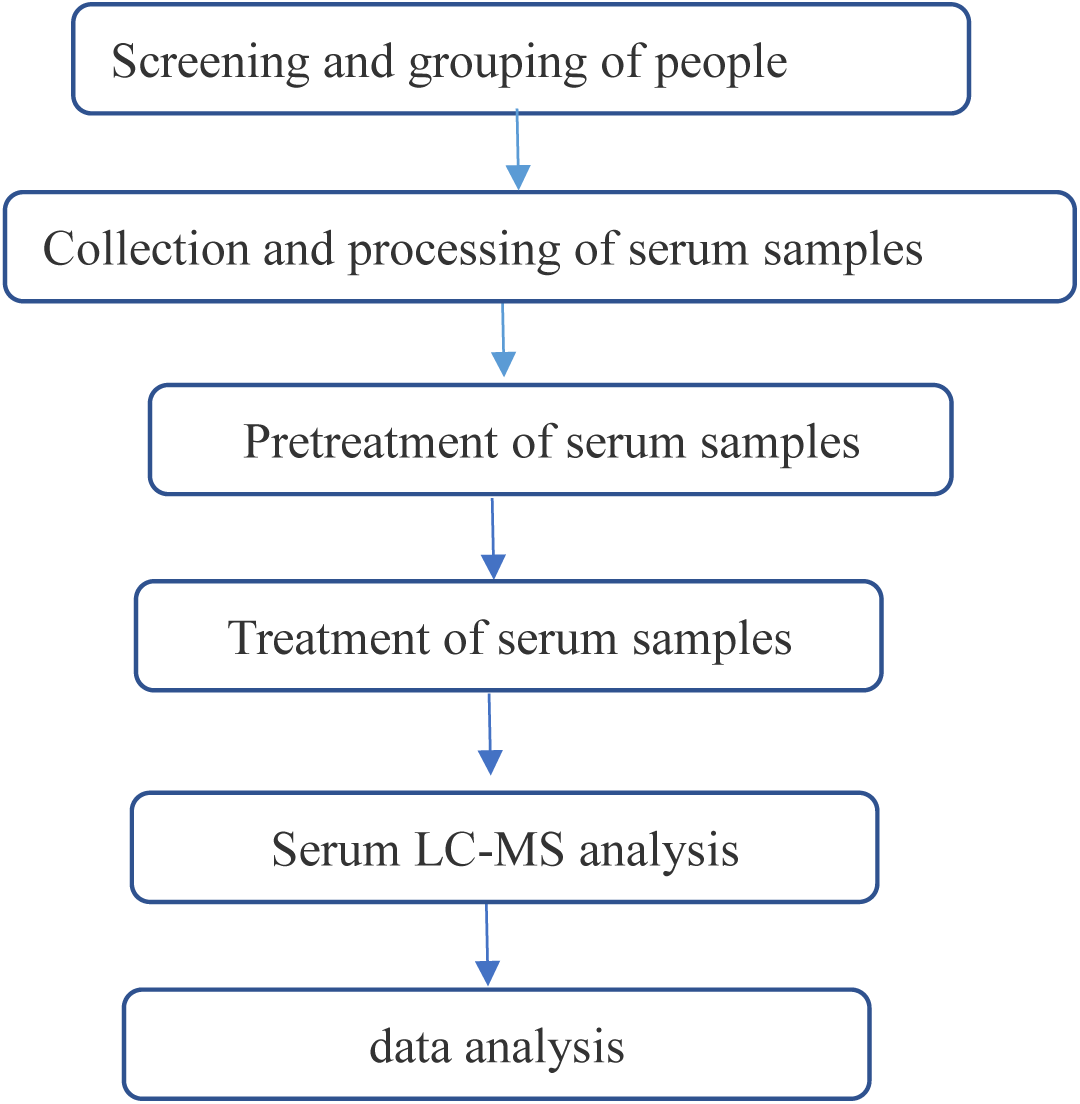
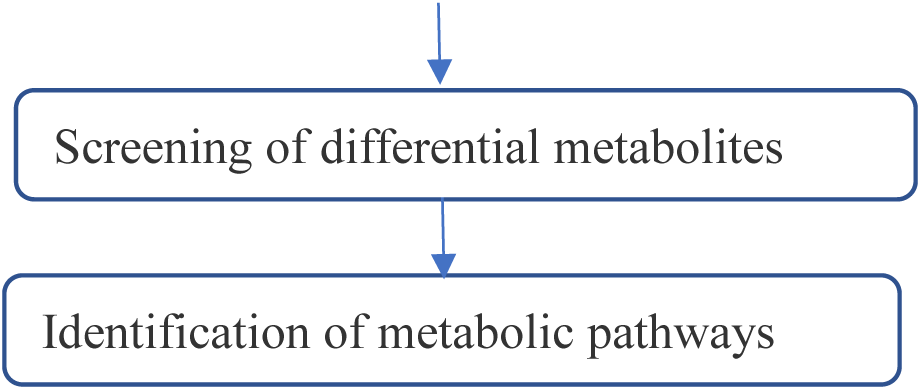

#### 1.1 experimental equipment

Dio U3000 UHPLC type high performance liquid chromatograph (Thermo Fisher Scientific Shier Technology Company), QE plus high resolution mass spectrometer (Thermo Fisher Scientific Shier Technology Company), F-060SD ultrasonic cleaner (Shenzhen Fuyang Technology Group Co., Ltd.), TYXH-I vortex oscillator (TYXH-I), TGL-16MS desktop high-speed freezing centrifuge (Shanghai Luxiangyi Centrifuge Instrument Co., Ltd.), LNG-T98 freeze concentration centrifugal dryer (Taicang Huamei Biochemical Instrument Factory), acquity UPLC HSS T3 (100mm× 2.1mm, 1.8um) chromatographic column (Waters).

#### 1.2 experimental reagents

Methanol, formic acid, water and acetonitrile are all purchased from Thermo Company, and L-2- chlorophenylalanine is purchased from Shanghai Hengchuang Biotechnology Co., Ltd. All chemicals and solvents are analytically pure or chromatographic grade.

#### 1.4 samples

##### 1.4.1 Source of samples

This study was approved by the Ethics Committee of Henan Provincial People’s Hospital, with the ethics batch number: 2018 Lun Shen No.73. Serum samples of ILD patients were collected from patients with interstitial lung diseases who were admitted to Department of Respiratory and Critical Care Medicine and Department of Rheumatology and Immunology of Henan Provincial People’s Hospital from January 1, 2021 to January 1, 2022. Patients in the healthy control group came from the physical examination center of Henan Provincial People’s Hospital without basic diseases. All ILD patients and healthy subjects signed informed consent forms. The inclusion and exclusion criteria of ILD patients are as follows: Inclusion criteria:

1. Male or female, aged 18-85 years, with high resolution CT data (high resolution reconstruction, thickness ≤ 1.25mm);
2. ILD patients were diagnosed as ILD by multi-disciplinary discussion (MDD), and the disease type was clear. All IPF patients meet the diagnostic criteria of idiopathic pulmonary fibrosis established by American Thoracic Society/European Respiratory Society/Japanese Respiratory Society/Latin American Thoracic Association in 2018 [15]. Allergic pneumonia conforms to the international evidence-based guidelines for adult HP diagnosis developed by American Thoracic Society/European Respiratory Society/Japanese Respiratory Society/Latin American Thoracic Society [16]. The diagnosis of pneumoconiosis, CTD-ILD and sarcoidosis respectively meet the consensus standards of Chinese experts on the diagnosis and treatment of pneumoconiosis, CTD-ILD and sarcoidosis [17, 18].
3. Sign the informed consent form and agree to complete this study.

Exclusion criteria:

1. Other diagnoses other than IPF, such as acute exacerbation of chronic obstructive pulmonary disease, pulmonary heart disease, pulmonary embolism, bronchial asthma and bronchiectasis;
2. Obvious disturbance of consciousness, obvious neuromuscular diseases such as myositis, muscle infection, myasthenia gravis, hemiplegia caused by cerebral infarction;
3. Patients with malignant tumors and hematological diseases can still be enrolled in the group if they achieve sustained complete response;
4. Complicated with heart, liver, kidney and other organ failure, except pulmonary heart disease;
5. The researcher can’t complete the researcher’s judgment;
6. Those with irregular information.

##### 1.4.2 Sample pretreatment

All eligible subjects collected 5ml of venous blood on an empty stomach in the morning, and all the blood samples were stored in a vacuum blood collection tube containing coagulant, placed at room temperature for 2 hours, centrifuged at 3000r/min for 10 minutes, and then 500µL of supernatant was sucked by a 1000µL pipette, placed in an EP tube, marked with a marker pen, and uniformly placed in a refrigerator at −80℃ for later use. Avoid repeated freezing and thawing to affect detection.

##### 1.4.3 General information and biochemical indicators

The pre-set data questionnaire was used to collect the clinical data of patients. Including (1) gender, age, course of disease, height, weight, smoking history and clinical symptoms; (2) Routine test results: blood routine, liver and kidney function, C-reactive protein, coagulation +D dimer, etc. (3) Lung function and other inspection indexes; (4) Complications, complications and treatment plan.

##### 1.4.4 Pre-processing of sample detection

Take out the sample stored at −80℃, thaw at room temperature, transfer 100 μL of the sample, and add the internal standard (L-2- chlorophenylalanine, 0.06 mg/ml; Methanol configuration) 20 μL, vortex oscillation for 10 s ; ; Add 300 μL protein precipitant methanol-acetonitrile (V:V=2:1) and vortex for 1 min;; Ultrasonic extraction in ice water bath for 10 min, standing at −20℃ for 30min; Centrifuge for 10 min(13000 rpm, 4℃), take 200 μL supernatant and put it into LC-MS injection vial to evaporate; Re-dissolving with 300μL methanol-water (V:V=1:4) (vortex for 30 s, ultrasonic for 3 min); Standing at −20℃ for 2 hours; Centrifuge for 10 min(13000 rpm, 4℃), suck 150 μL of supernatant with syringe, filter with 0.22 μm organic phase pinhole filter, transfer to LC injection vial, and store at −80℃ until LC- MS analysis.

#### 1.5 test conditions

(1) chromatographic conditions and mass spectrometry conditions: the analytical instrument of this experiment is a liquid chromatography-mass spectrometry system consisting of Dionex U3000 UHPLC ultra-high performance liquid in series with QE plus high resolution mass spectrometer. Chromatographic conditions: chromatographic column: acquity uplc HSS T3 (100mm× 2.1mm, 1.8um); Column temperature: 45℃; The mobile phase includes A- water (containing 0.1% formic acid) and B- acetonitrile (containing 0.1% formic acid); Flow rate: 0.35 mL/min;; Injection volume: 2 μL l. Ion source: ESI, sample mass spectrometry collect positive and negative ion scanning modes respectively. The mobile phase elution gradient and mass spectrometry conditions are as follows.

#### 1.6 data processing

##### 1.6.1 Preparation and collation of data sets

The raw data of serum detected by liquid chromatography-mass spectrometry and mass spectrometry were subjected to baseline filtering, peak identification, integration, retention time correction, peak alignment and normalization by metabonomics processing software Progenesis QI v2.3 (Nonlinear Dynamics, Newcastle, UK). Main parameters: preamble tolerance: 5ppm/10ppm (self-built library), product tolerance: 10 ppm/20ppm (self-built library), product ion threshold: 5%. The identification of compounds is based on accurate mass number, secondary fragments and isotope distribution. The human metabolome database (HMDB), Lipidmaps(v2.3), METLIN database and self-built database are used for qualitative analysis. For the extracted data, delete the ion peaks with missing values (0 value) > 50% in the group, replace the 0 value with half of the minimum value, and screen the qualitatively obtained compounds according to the Score of the qualitative results of the compounds. The screening standard is 36 points (out of 60 points), and those below 36 points are regarded as inaccurate qualitative results and deleted. Finally, the positive and negative ion data are combined into a data matrix table, which contains all the information extracted from the original data and can be used for analysis, and the subsequent analysis is based on this.

##### 1.6.2 Data analysis

1. Multivariate statistical analysis: firstly, unsupervised principal component analysis (PCA) will be used to observe the overall distribution among samples and the stability of the whole analysis process, and then supervised partial least squares analysis (PLS-DA) and orthogonal partial least squares analysis (OPLS-DA) will be used to distinguish the overall differences of metabolic profiles among groups and find out the differential metabolites among groups.
2. In the orthogonal partial least squares analysis (OPLS-DA), the potential biomarkers were found by screening VIP>1, and the data of each two groups were tested by SPSS21.0 statistical software. If they did not conform to the normal state, the metabolites with P<0.05 were screened out by nonparametric test. Only when the VIP value > 1 and P<0.05 were satisfied, they could be screened out for the next analysis.
3. Using software to make heat map, Hierarchical Clustering the expression levels of all significantly different metabolites, and observe the changes of the expression levels of different metabolites between the two groups.
4. On the basis of remarkable enrichment in metabolic pathway, the differential metabolites of IPF group and healthy control group, CTD-ILD group and healthy control group, other groups and healthy control group, IPF group and CTD-ILD group, IPF group and other groups, CTD-ILD group and other groups were analyzed with ROC curve by SPSS21.0 software to find out the differential metabolites with strong discrimination ability. For the correlation between two significant metabolites, Pearson correlation coefficient is used to measure the degree of linear correlation between the two metabolites, with red indicating positive correlation and blue indicating negative correlation.
5. The general clinical data of the patients in the group were collected by Excel 2019 and statistically analyzed by SPSS21.0, and the continuous variables conforming to normal distribution were expressed by mean standard deviation 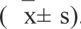. Parameters that do not conform to normal distribution are expressed by median and quartile spacing M(P25, P75). According to the normal distribution, the comparison between two groups adopts T test, and the comparison among multiple groups adopts one-way ANOVA. If it does not conform to the normal distribution or the variance is uniform, Mann-Whitney U test is used between two groups, and Kruskal⁃Wallis rank sum test of independent samples is used for comparison among multiple groups. Chi-square test was used to compare the classification and rate. P0.05, the difference was statistically significant. 1.6.3 Identification of the structure of differential metabolites The molecular weight of QE high-resolution mass spectrometry and the molecular formula of key differential metabolites were inferred from abundance, and the possible structure was identified by HMDB online database.

##### 1.6.4 Enrichment analysis of metabolic pathways of key differential metabolites

The pathway enrichment analysis of differential metabolites is helpful to understand the mechanism of metabolic pathway change in differential samples. Metabolic pathway enrichment analysis of differential metabolites based on KEGG database.

## 2 Results

### 2.1 Analysis of general information and clinical biochemical indexes of research objects Participants

There were 26 IPF patients, including 21 males and 5 females, with an average age of 70.62 7.99 years; There were 21 patients with CTD-ILD, including 8 males and 13 females, with an average age of 61.14 13.53 years. There were 23 other patients, including 8 males and 15 females, with an average age of 52.17 9.03 years. 20 healthy controls, including 7 males and 13 females, with an average age of 42.60 14.56 years. The clinical data of all patients in the group are complete, ILD patients are diagnosed as ILD through clinical-imaging-pathology multi-disciplinary discussion, and the disease types are clear. The general data of patients include gender, age, smoking history and clinical symptoms, which include fever, cough, expectoration, chest tightness, dyspnea and fatigue. Clinical biochemical indicators include: white blood cell count, neutrophil count, lymphocyte count, eosinophil count, red blood cell count, albumin, creatine kinase, creatine kinase isoenzyme, lactate dehydrogenase, D- dimer and C-reactive protein, as shown in Table 3 and Table 4. There were significant differences in age, sex and smoking history among the four groups (P < 0.05). For the clinical symptoms of IPF group, CTD-ILD group and other groups, there were significant differences in cough and expectoration (P < 0.05), and the differences were statistically significant. Fever, chest tightness, dyspnea and fatigue have no significant difference, which is statistically insignificant. For clinical biochemical indicators, the percentage of eosinophil count, creatine kinase, creatine kinase isoenzyme and C-reactive protein, there were significant differences among IPF group, CTD-ILD group and other groups, which were statistically significant; for the percentage of eosinophil count, creatine kinase and C-reactive protein, there were significant differences between IPF group and other groups, P < 0.05, which was statistically significant; The isoenzyme of creatine kinase in IPF group was lower than that in CTD-ILD group (P < 0.05), the difference was statistically significant, while that in CTD-ILD group was lower than that in other groups (P < 0.05), the difference was statistically significant, and other clinical biochemical indexes, P > 0.05, had no significant difference.

**Table 1.**
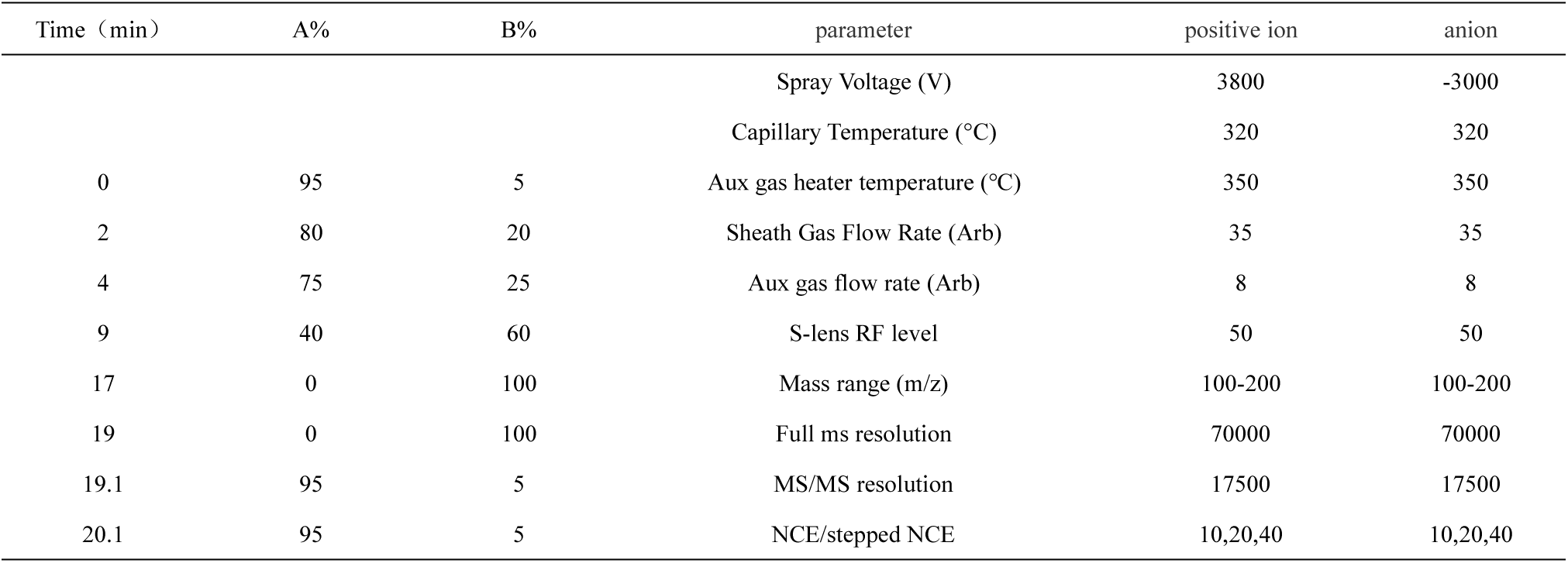
Elution gradient and mass spectrometry conditions

**Table 2.** Distribution of disease types of patients in the group

| Disease type | Case number |
| --- | --- |
| Idiopathic pulmonary fibrosis | 26 |
| Rheumatoid-associated interstitial lung | 6 |
| Autoimmune interstitial pneumonia | 5 |
| Inflammation-associated interstitial pneumonia | 7 |
| Sjogren's syndrome-related interstitial pneumonia | 3 |
| Alveolar proteinosis | 2 |
| sarcoidosis | 5 |
| cryptogenic organizing pneumonia | 5 |
| Allergic pneumonia | 7 |
| pneumoconiosis | 1 |
| sinus histiocytosis with massive lymphadenopathy | 1 |
| Nonspecific interstitial pneumonia | 2 |

**Table 3.**
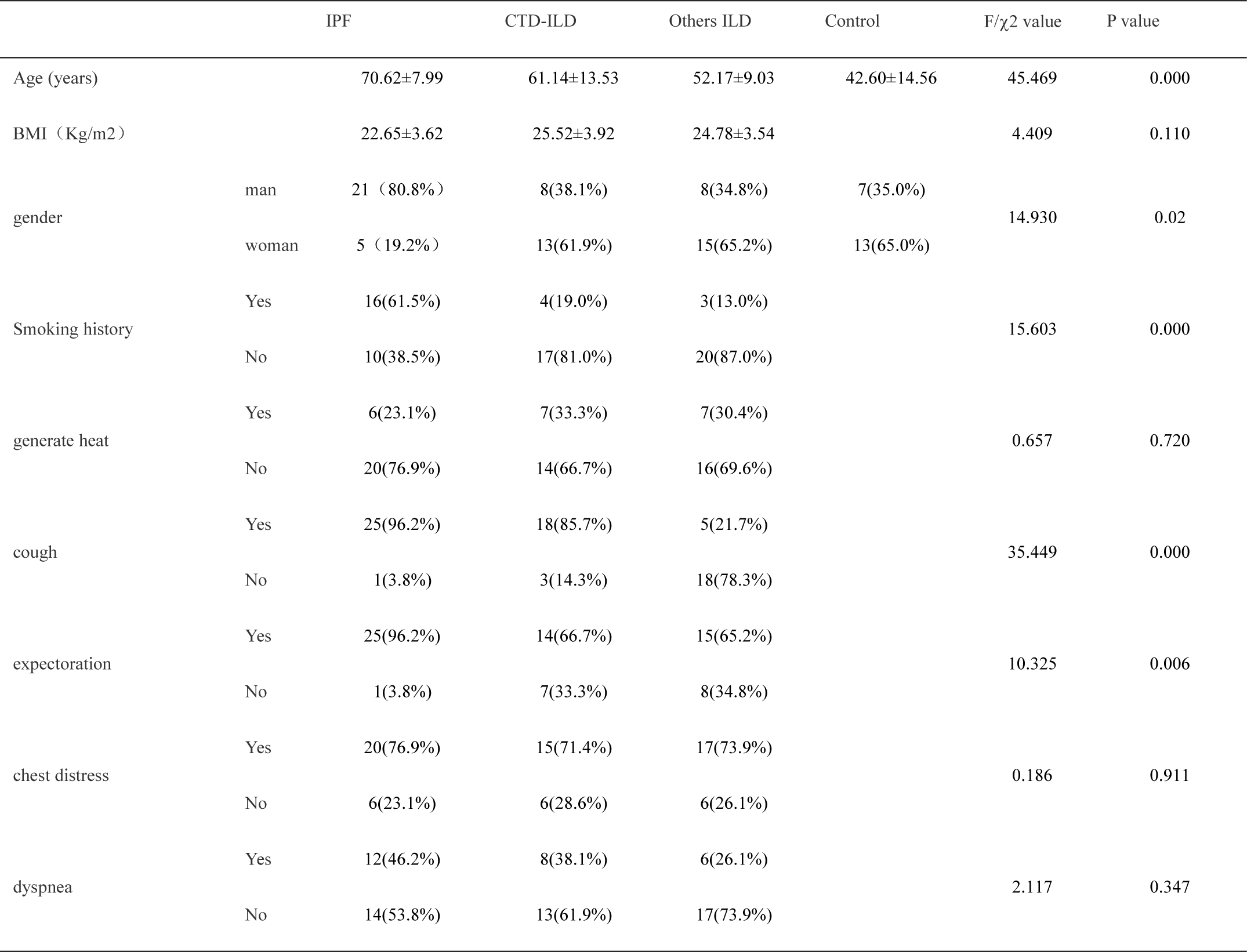
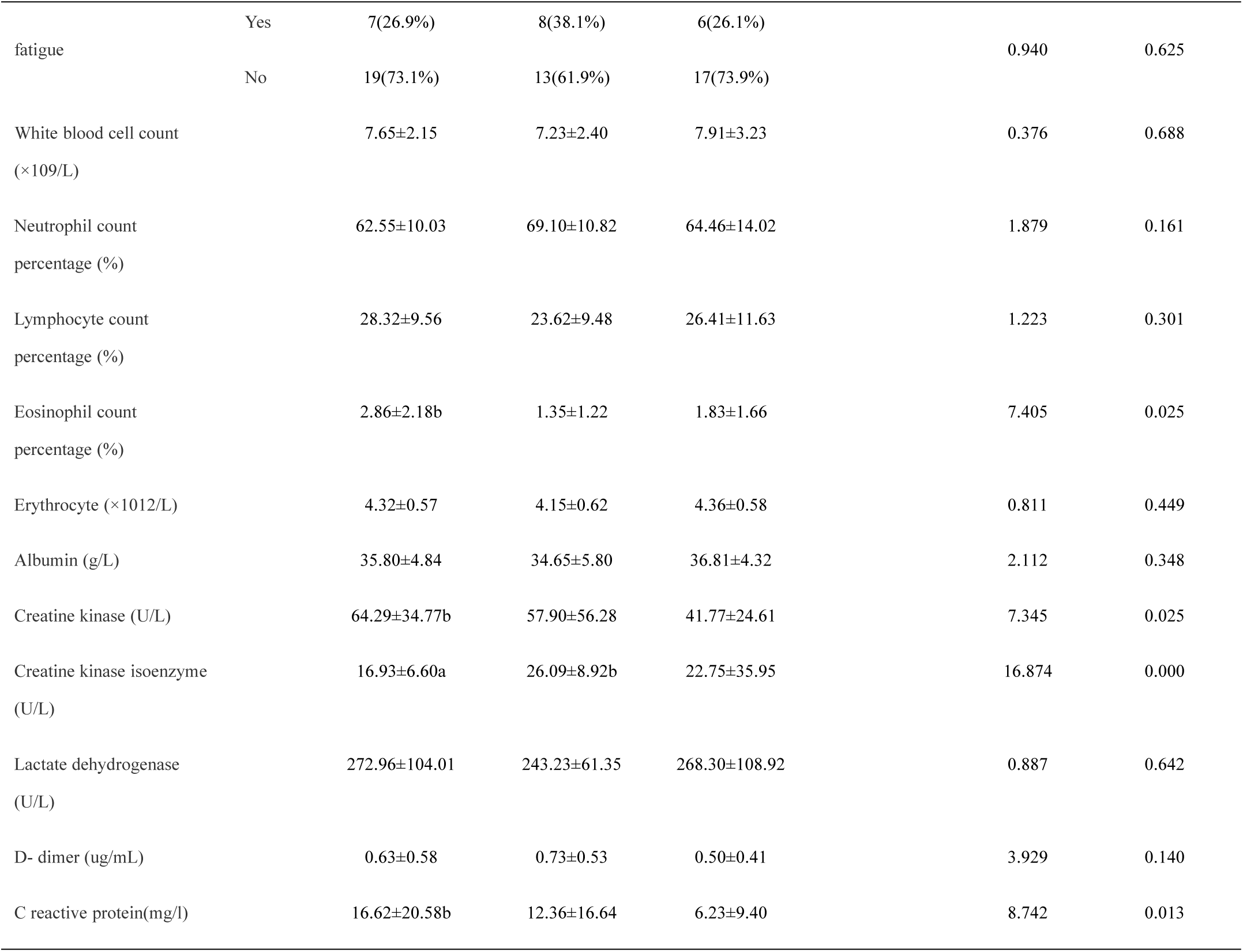
General clinical data and biochemical indexes of each group Note: IPF: Idiopathic pulmonary fibrosis; CTD-ILD: connective tissue-associated interstitial pneumonia; ILD: interstitial pneumonia; BMI: body mass index; A:CTD-ILD group b: other ILD

**Table 4.**
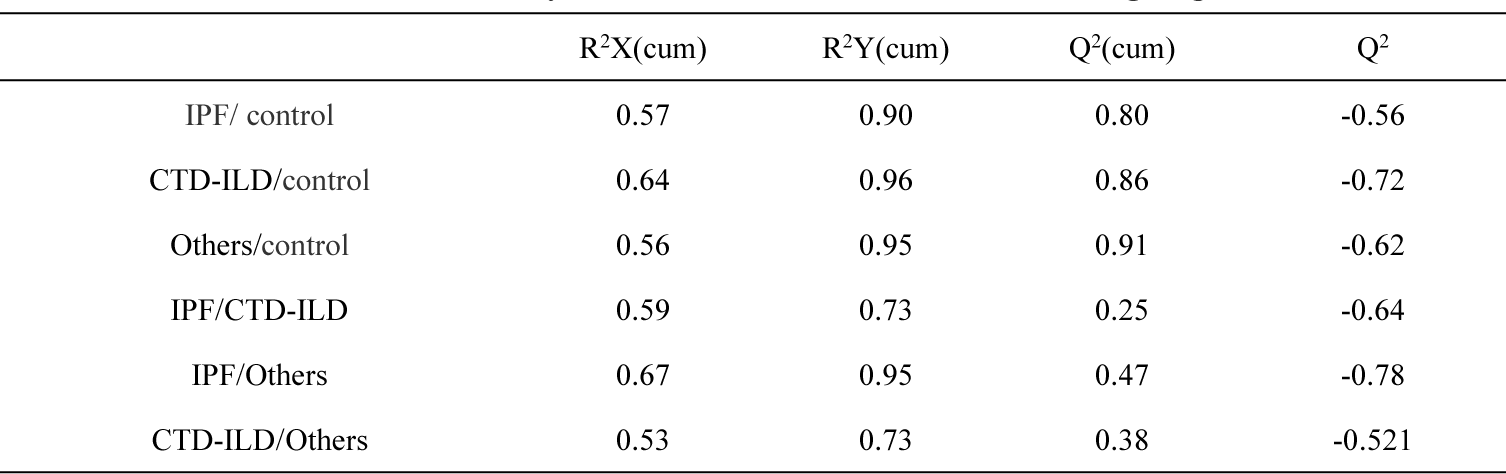
Analysis of OPLS-DA model between two groups

### 2.2 LC-MS total ion flow diagram of serum samples

As shown in Figures 1 and 2, A, B, C and D respectively represent the typical total ion flow diagrams of metabolites of healthy control group, IPF group, CTD-ILD group and other groups in positive ion mode, and E, F, H and I respectively represent the typical total ion flow diagrams of metabolites of healthy control group, IPF group, CTD-ILD group and other groups in negative ion mode. From the ion flow diagrams, it can be seen that whether in positive ion mode or not There is a difference in the intensity of ion peak among groups within a certain time, so we can draw the conclusion that there are differences in serum metabolite maps among groups, and there are different metabolites among groups, but whether they are statistically significant or not requires statistical analysis.

**Fig. 1.**
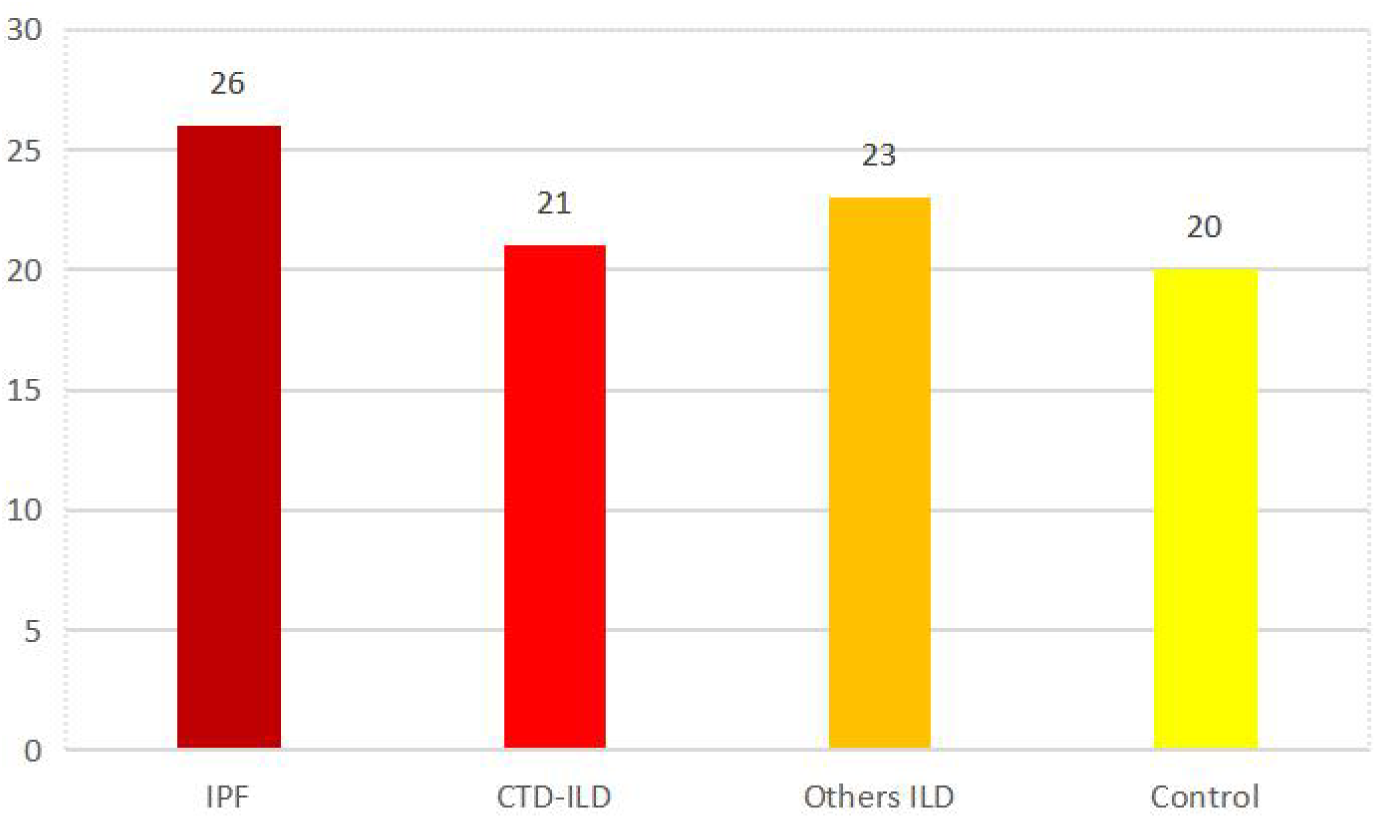
Grouping of enrolled patients

**Fig. 2.**
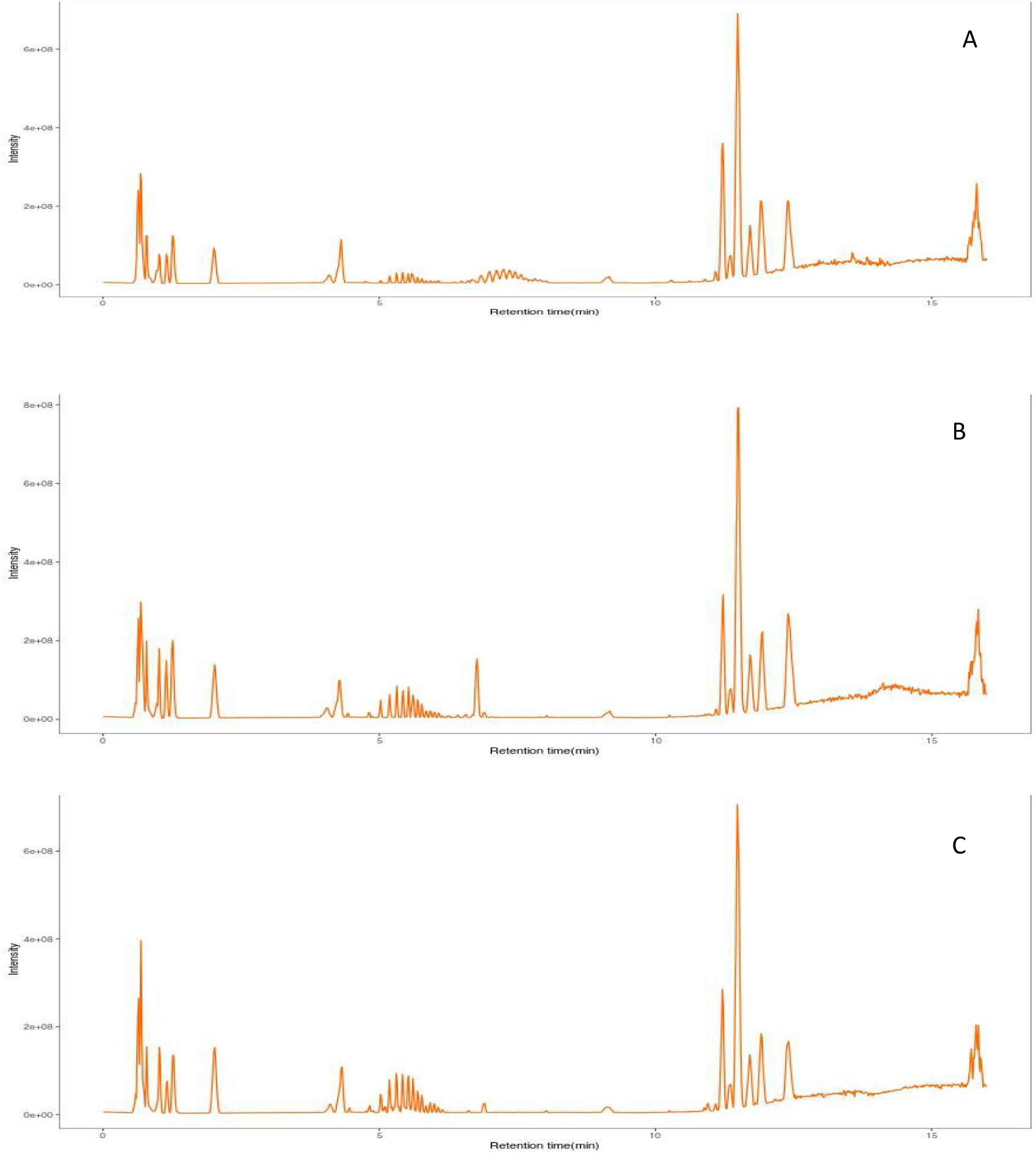
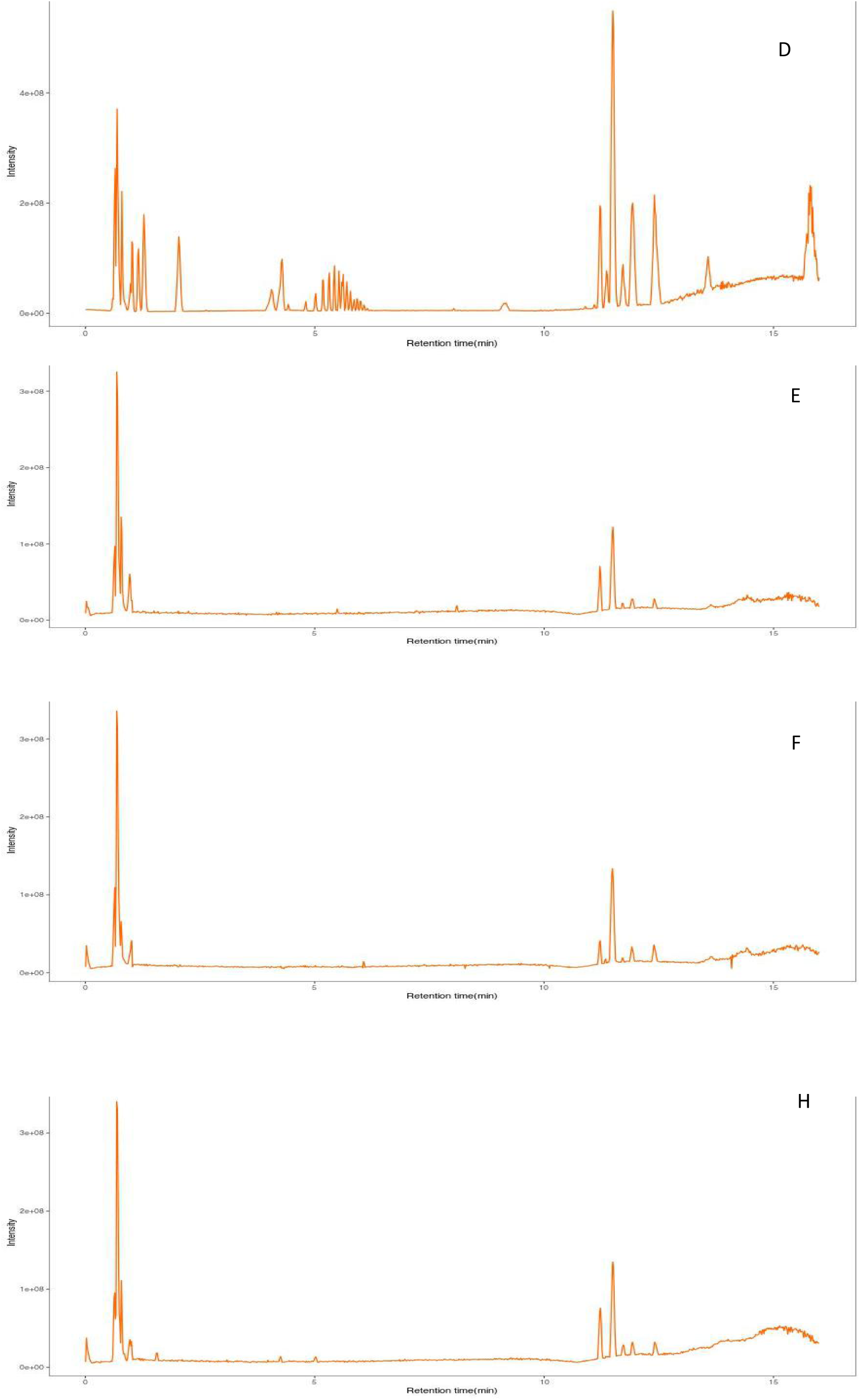
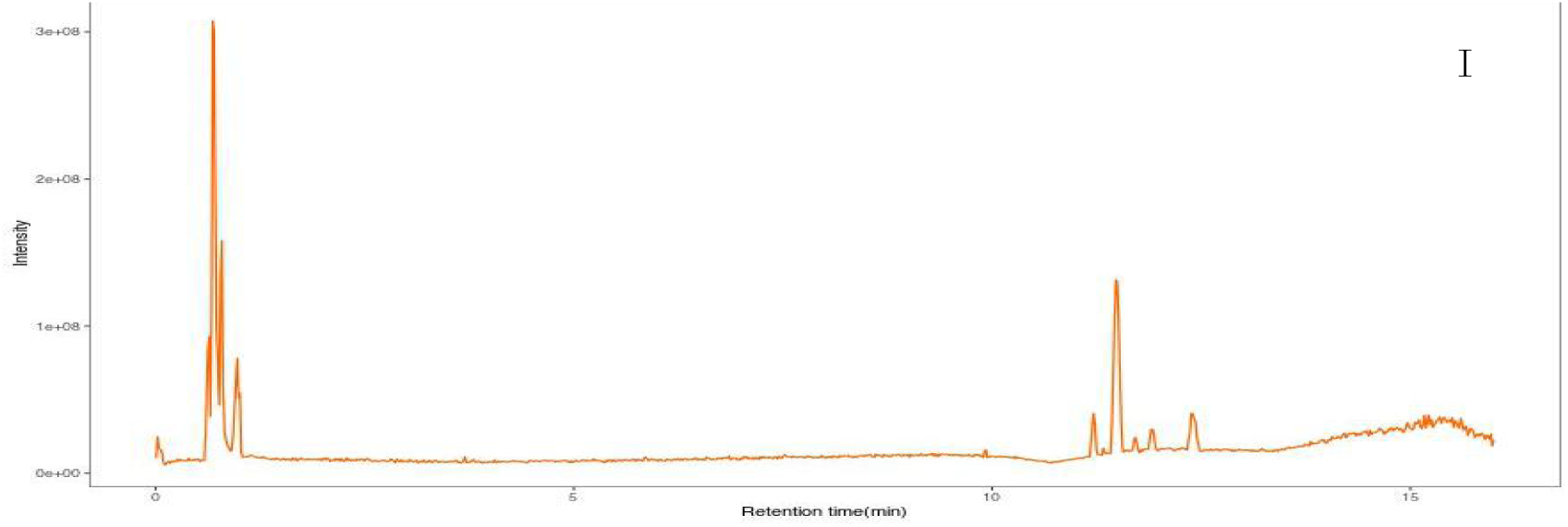
LC-MS (positive and negative ions) total ion flow diagram of each group Note: Positive ion mode A: healthy control group; B: IPF group; C: CTD-ILD group; Other groups d; Negative ion mode E: healthy control group; F: IPF group; H: ctd-ILD group; I: other groups

### 2.3 orthogonal partial least squares discriminant analysis of serum samples (OPLS-DA)

In the OPLS-DA score chart, the yellow diamond represents IPF group, the red triangle represents CTD-ILD group, the purple inverted triangle represents other groups, and the blue square represents the healthy control group. The position of the sample is determined by the difference of metabolites, that is, the more similar the metabolites are, the closer the position of the sample is, and the greater the difference, the farther the spatial distribution distance is. As shown in Figure 3, in the positive ion mode, compared with the healthy control group in IPF group, CTD-ILD group and other groups respectively, it was found that they were distributed on both sides of Y axis and separated obviously, so there were different metabolites among them; Compared with the IPF group and CTD-ILD group, it was found that the metabolites of IPF group and CTD-ILD group were distributed on both sides of Y axis, which indicated that there were different metabolites between IPF group and CTD-ILD group. In the OPLS-DA scores of IPF group and other groups, CTD-ILD group and other groups, it was found that the metabolites of each group were partially distributed on both sides of Y axis, which indicated that there was little difference between IPF group and other groups, CTD-ILD group and other groups.

**Fig. 3.**
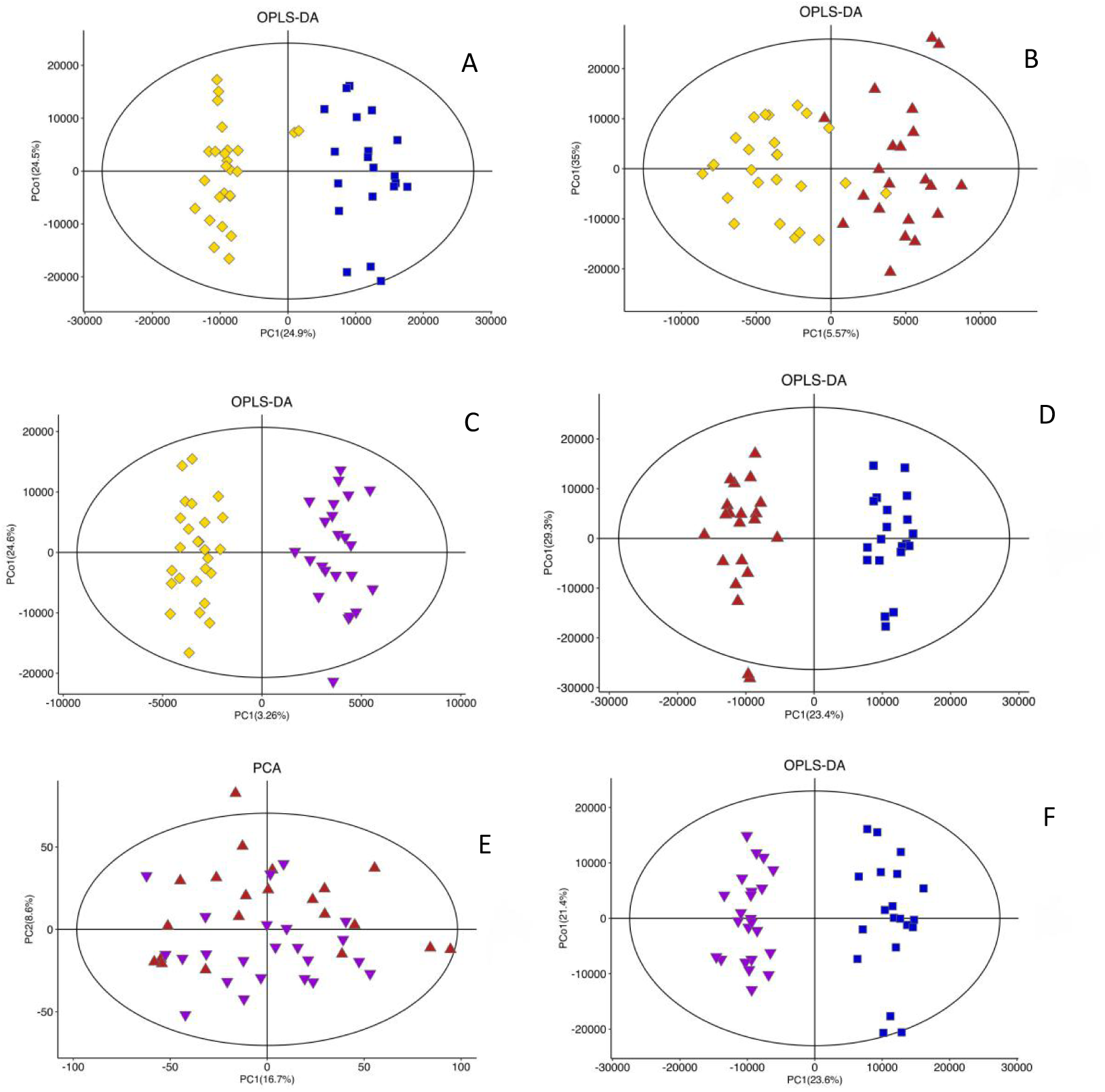
OPLS-DA score between groups (positive ion mode) Note: A: OPLS-DA scores of A: IPF group and healthy control group; B: OPLS-DA scores of B: IPF group and CTD-ILD group; C: OPLS-DA scores of C: IPF group and Others group; D: OPLS-DA scores of D: CTD-ILD group and healthy control group; E: OPLS-DA scores of E: CTD-ILD group and Others group; F: OPLS-DA scores of others group and healthy control group; IPF group (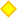); CTD-ILD group (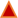); Others group (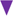); Control group (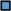)

#### 2.3.2 quality evaluation of opls-da model

To evaluate the quality of OPLS-DA model, R2(R2X(cum), R2Y(cum)) and Q2(cum) are the main parameters. All R2 values are greater than 0.5. In the models of IPF group, CTD-ILD group, IPF group and other ILD groups, Q2 values are less than 0.5, and other models all Q2 values are greater than 0.5, which shows that the analysis of the model data is reliable. It can be seen from Table 4 that in the positive and negative ion mode, the model quality of IPF group and healthy control group, CTD-ILD group and healthy control group, other groups and healthy control group is better, and the data analysis is more reliable, which can better explain the differences between the two groups.

#### 2.3.3 S-polt diagram of OPLS-DA of serum samples

On the basis of the analysis results of OPLS-DA scores between the two groups, it is determined that there are metabolic differences between the two groups. Through the analysis of S-polt graphs (Figure 4), we can further find out the different metabolites. The abscissa is the characteristic value of the influence of metabolites on the comparison group, and the ordinate is the correlation between sample scores and metabolites. Each green dot represents a compound. The farther away from the origin, the greater the contribution of this compound to the grouping. The metabolites closer to the upper right corner and the lower left corner indicate the more significant differences.

**Fig. 4.**
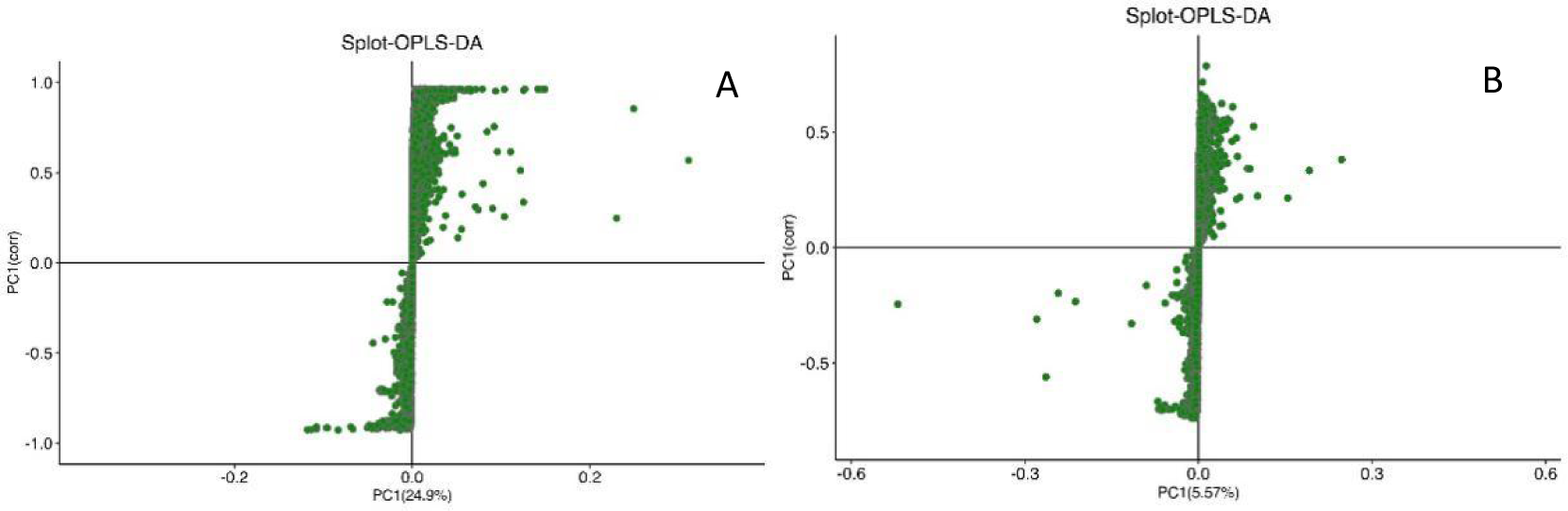
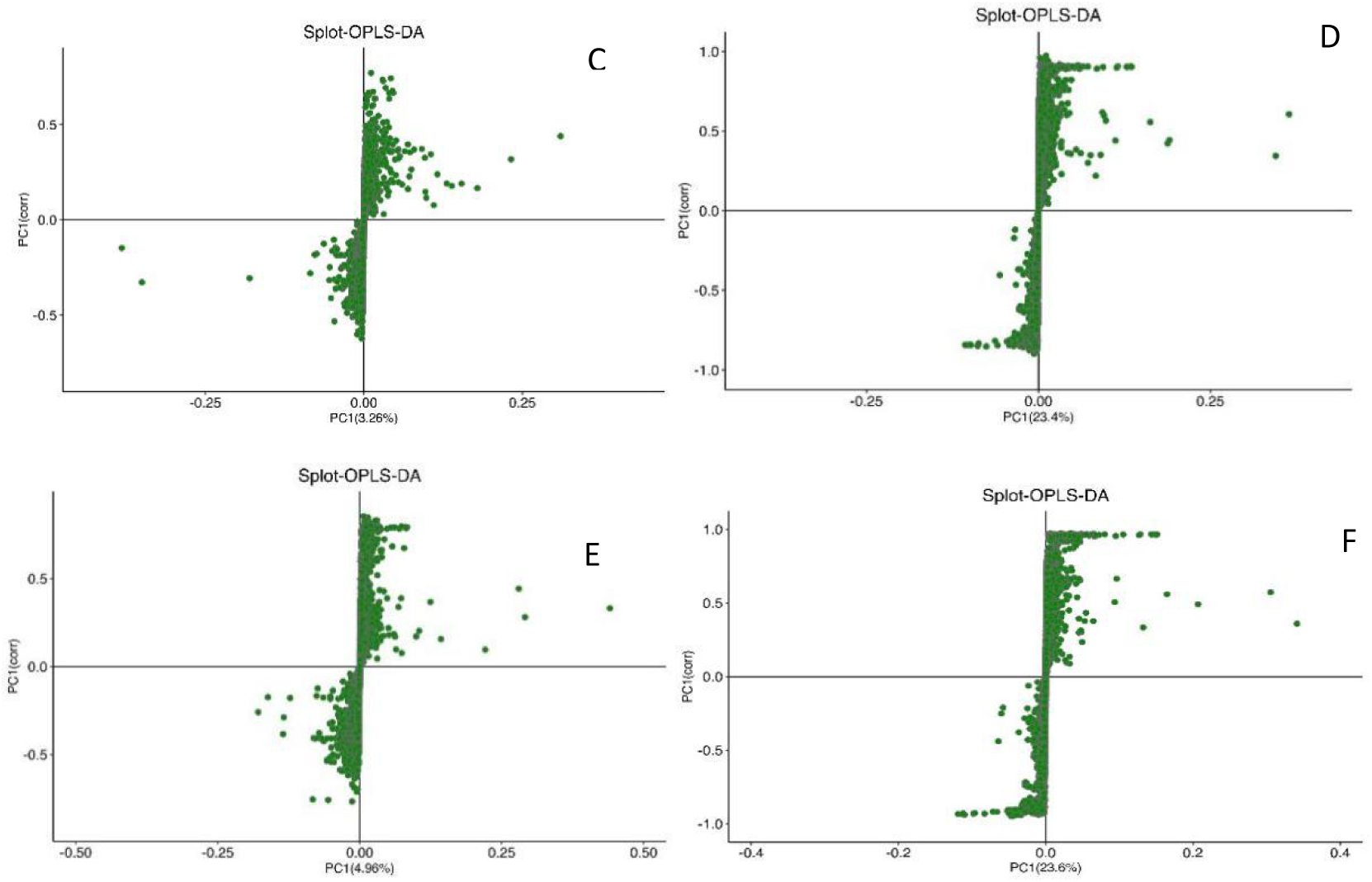
S-polt diagram of OPLS-DA between two groups (positive ion mode) Note: A: S-polt diagram of OPLS-DA in A: IPF group and healthy control group; B: S-polt diagram of OPLS-DA of B:IPF group and CTD-ILD group; S-polt diagram of OPLS-DA of CIPF group and other groups; D: S-polt diagram of OPLS-DA in D: CTD-ILD group and healthy control group; S-polt diagram of ECTD-ILD group and other groups OPLS-DA; F: S-polt diagram of OPLS-DA in other groups and healthy control group

### 2.4 Screening and identification of differential metabolites in serum samples

Univariate statistical analysis was used to screen the differential metabolites between groups. The screening criteria were VIP value of first principal component of OPLS-DA model > 1, P of T test < 0.05, and metabolites meeting the criteria were used as potential biomarkers. There are 193 kinds of differential metabolites between IPF group and healthy control group, including 177 kinds in positive ion mode and 16 kinds in negative ion mode. There are 115 differential metabolites in IPF group and CTD-ILD group, 85 in positive ion mode and 30 in negative ion mode. There are 101 differential metabolites in IPF group and other groups, 68 in positive ion mode and 33 in negative ion mode. There were 188 differential metabolites in CTD-ILD group and healthy control group, 169 in positive ion mode and 19 in negative ion mode. There are 148 differential metabolites in CTD-ILD group and other groups, 117 in positive ion mode and 29 in negative ion mode. There were 169 kinds of differential metabolites between other groups and healthy control group, 156 kinds in positive ion mode and 13 kinds in negative ion mode. As shown in Table 5 and Figure 5. In order to more intuitively show the relationship between samples and the expression differences of metabolites among different samples, we Hierarchical Clustering the expression amounts of all significant difference metabolites, and screen out the top 50 kinds of the most significant difference metabolites, with the name of the samples on the abscissa and the difference metabolites on the ordinate.

**Fig. 5.**
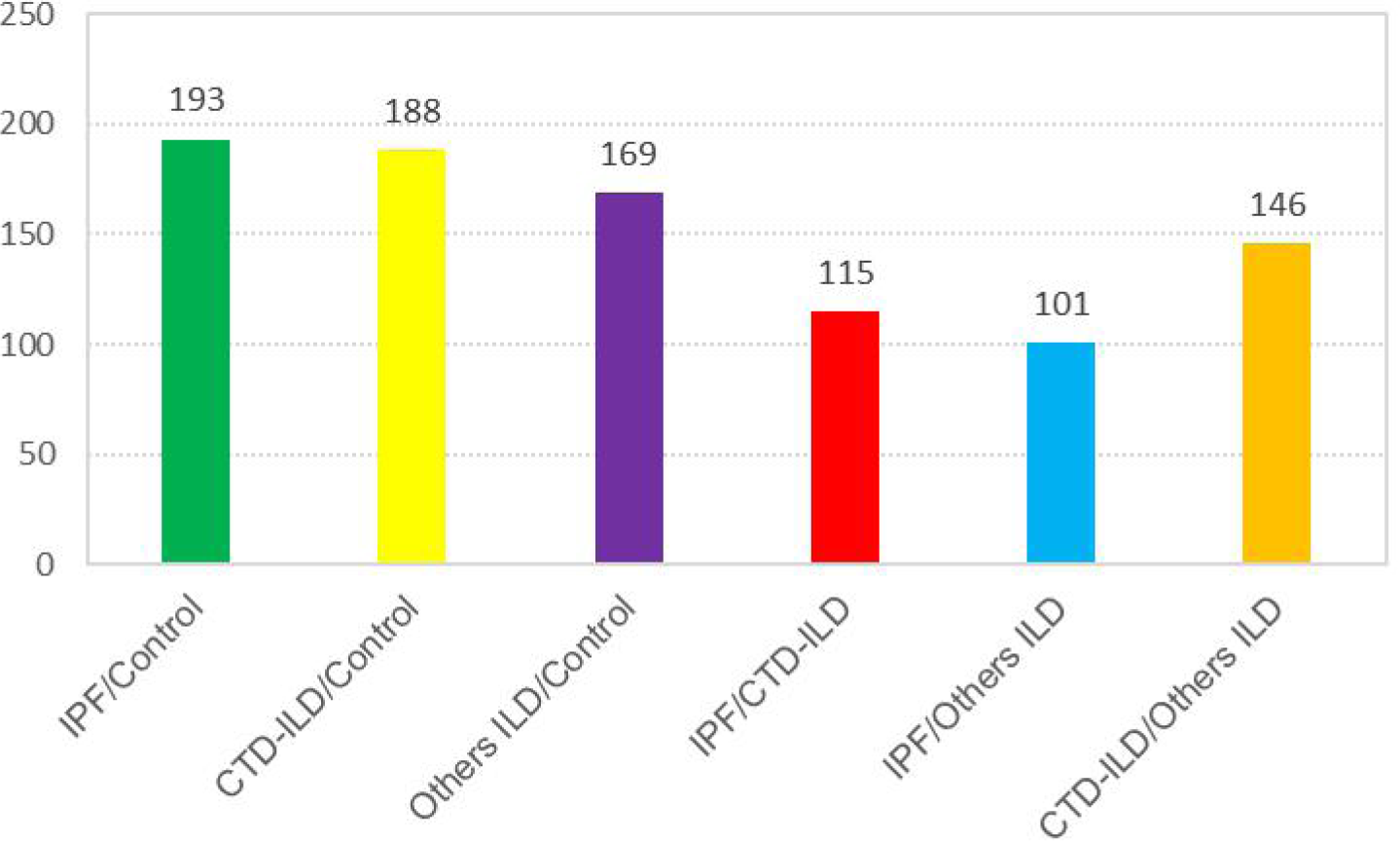
Histogram of the difference metabolic species number between two groups

**Table 5.** Difference of metabolic species between two groups

| Comparison group | Differential metabolic<br>species number | Number of species of ion differential metabolism |  |
| --- | --- | --- | --- |
|  |  | Positive ion mode | Negative ion mode |
| IPF and control | 193 | 177 | 16 |
| IPF and CTD-ILD | 115 | 85 | 30 |
| IPF and Others | 101 | 68 | 33 |
| CTD-ILD and Control | 188 | 169 | 19 |
| CTD-ILD and Others | 146 | 117 | 29 |
| Others and Control | 169 | 156 | 13 |

The color from blue to red indicates the expression abundance of metabolites from low to high, that is, the redder indicates the higher the expression abundance of differential metabolites. Figure 6.

**Fig. 6.**
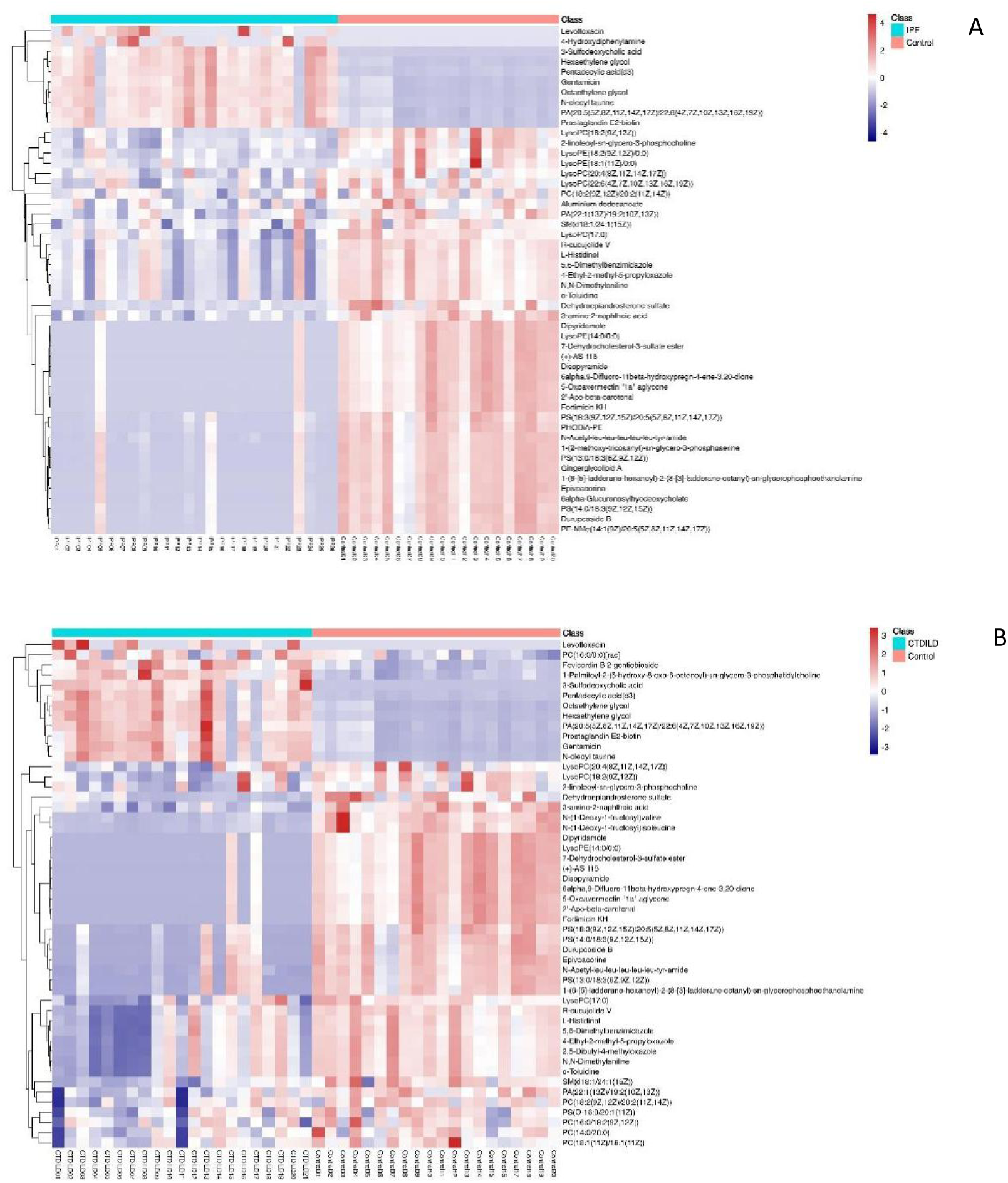
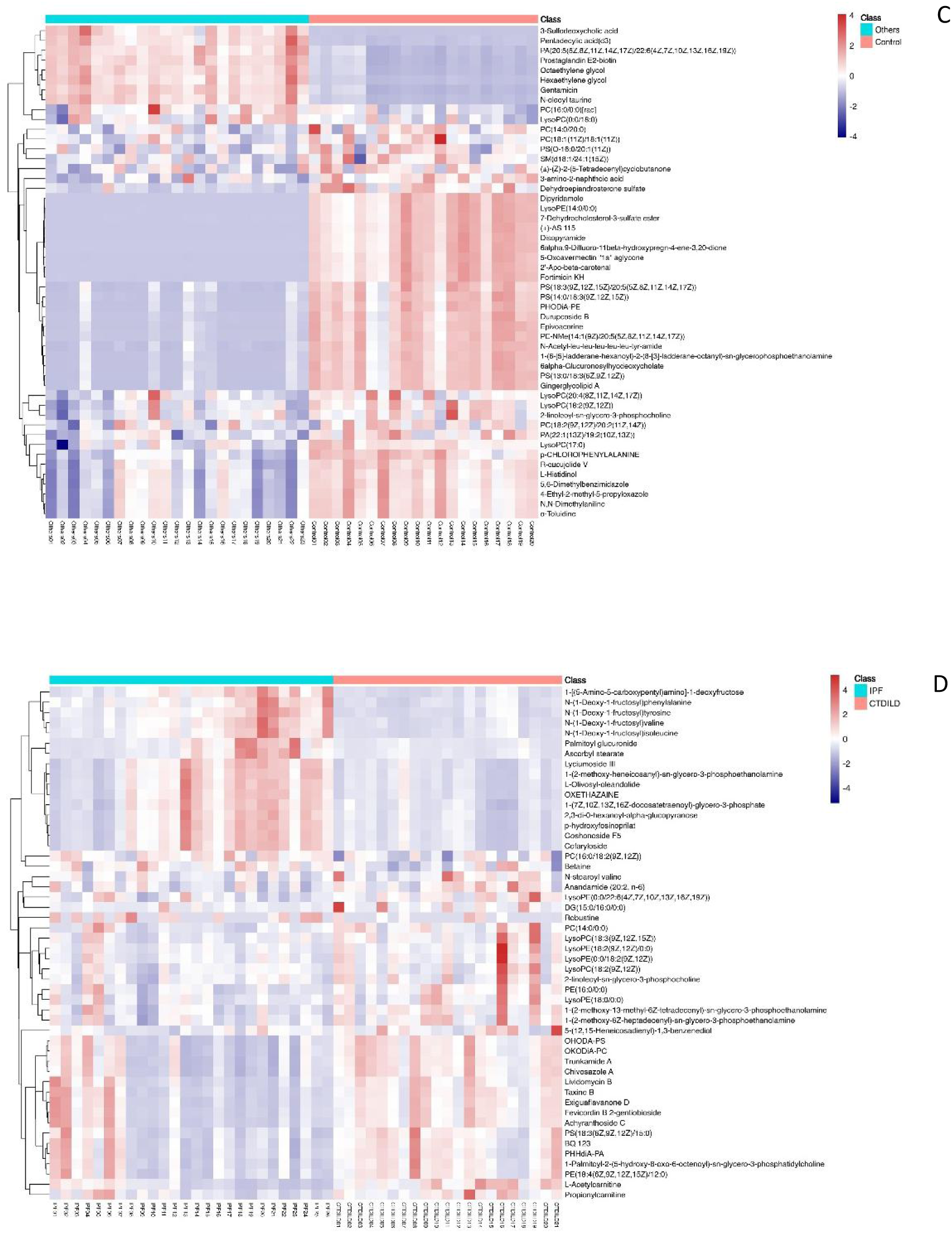
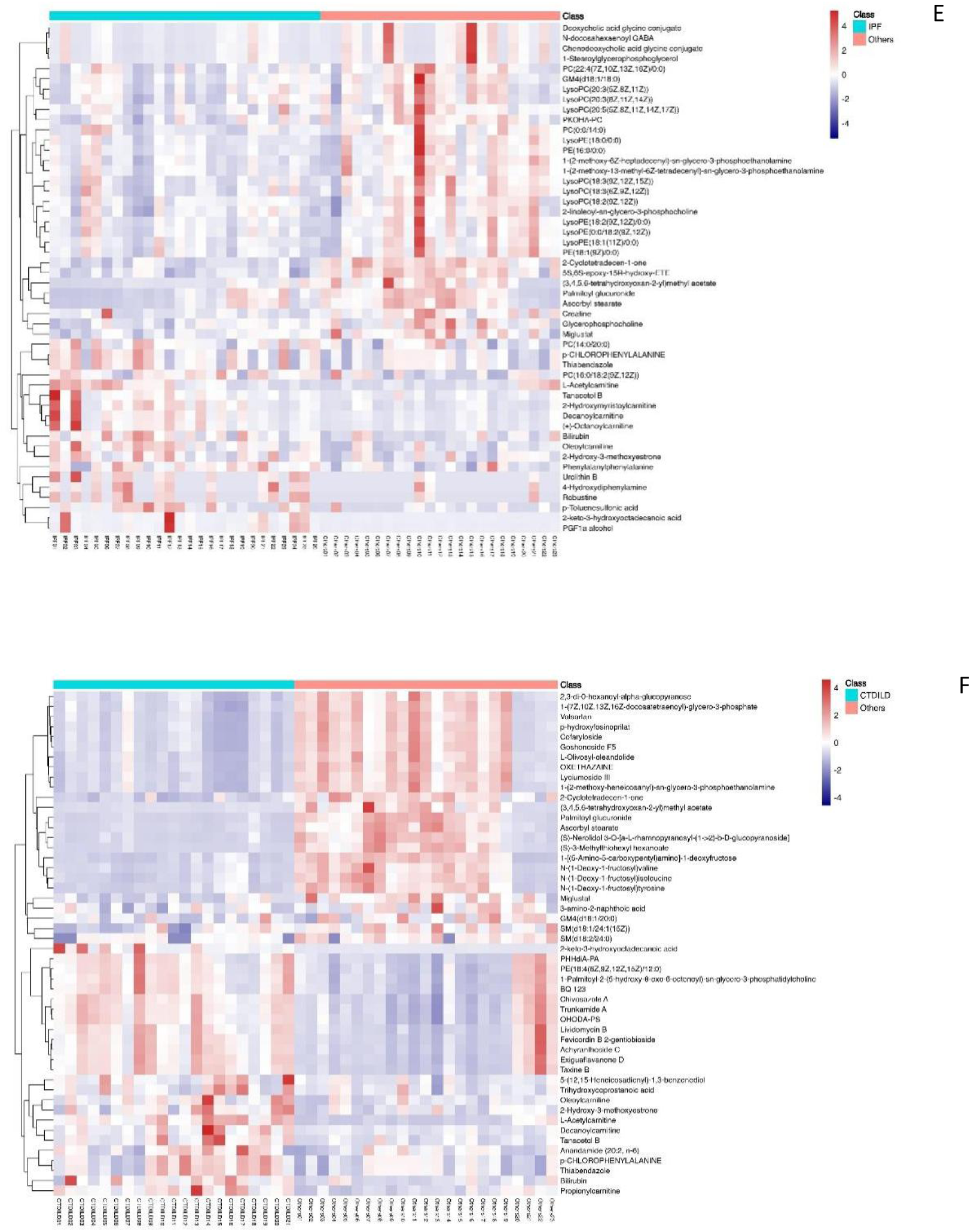
Thermal map analysis of differential metabolites Note: A: Thermographic analysis of the first 50 differential metabolites between A: IPF group and healthy control group; B: Thermographic analysis of the first 50 differential metabolites between B: CTD-ILD group and healthy control group; C: Thermogram analysis of the first 50 kinds of differential metabolites between other groups and healthy control group; D: Thermographic analysis of the first 50 differential metabolites in D: IPF group and CTD-ILD group; E: Thermogram analysis of the first 50 differential metabolites between E: IPF group and other groups; Thermographic analysis of the top 50 differential metabolites in F: CTD-ILD group and other groups

### 2.5 Correlation analysis of differential metabolites

Through correlation analysis of the first 50 metabolites with significant differences, according to the relevant statistical knowledge, when the absolute value of R is greater than 0.8, it is considered to be strongly correlated, and when the absolute value of R is equal to 1, it is completely correlated. Through correlation matrix analysis, red represents positive correlation and blue represents negative correlation. The correlation analysis of each group is shown in Figure 7.

**Fig. 7.**
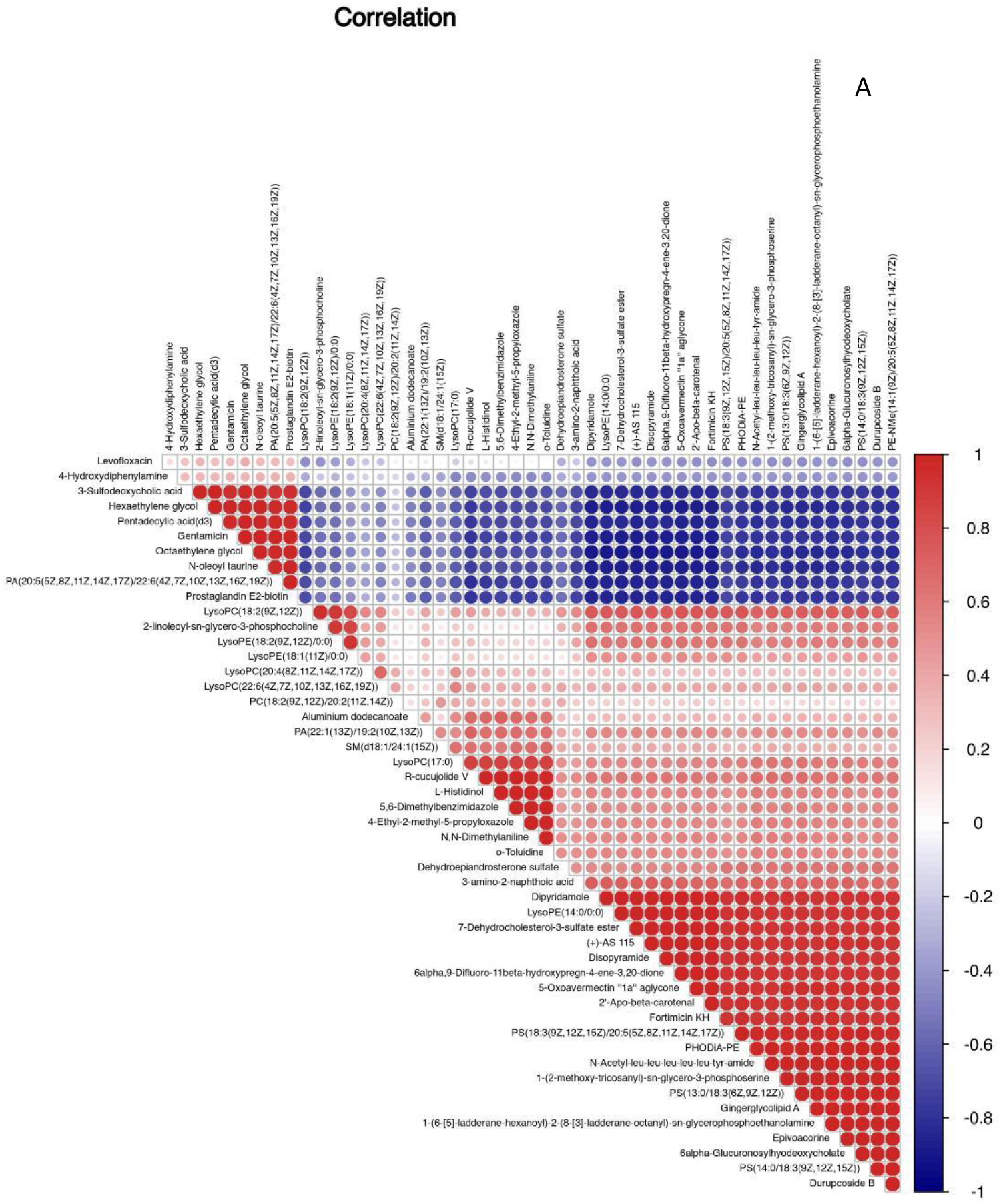
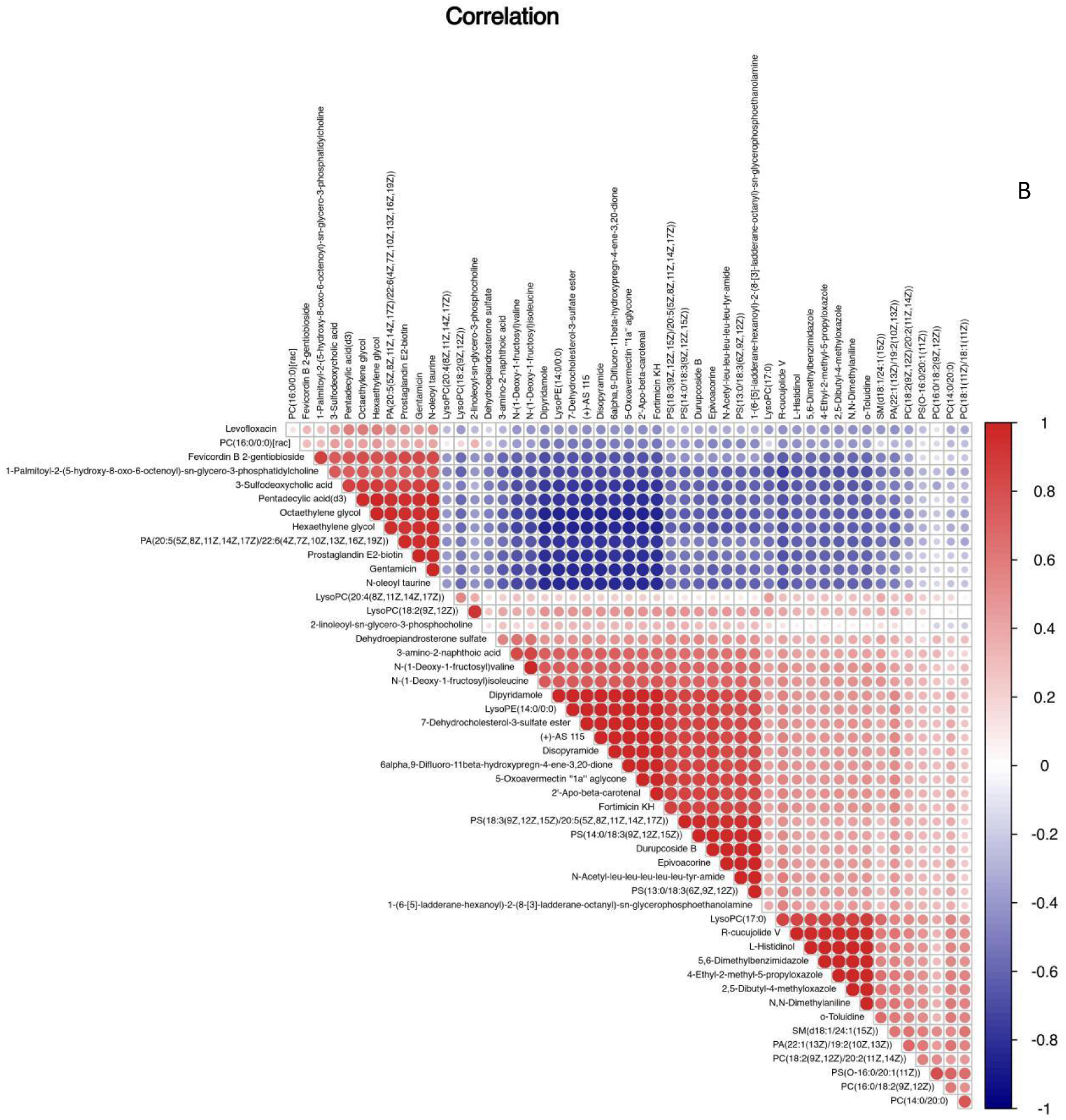
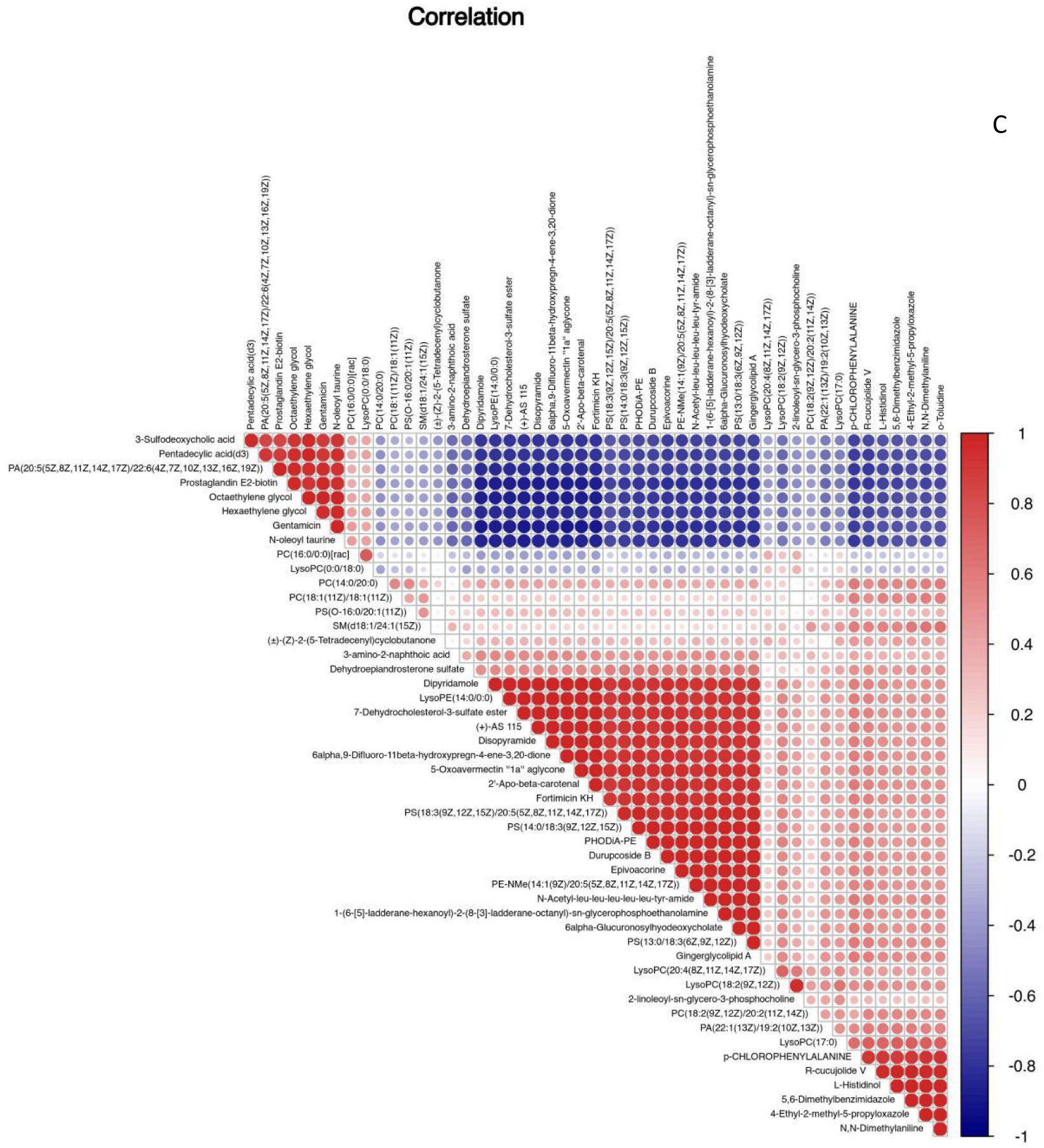
Correlation analysis of differential metabolites Note: A: Correlation analysis of differential metabolites between A: IPF group and healthy control group; B: Correlation analysis of differential metabolites between B: CTD-ILD group and healthy control group; C: Correlation analysis of differential metabolites between other groups and healthy control group

### 2.6 Metabolic pathway analysis of differential metabolites

On the basis of KEGG database, we analyze the metabolic pathway enrichment of the differential metabolites, and use hypergeometric test to find out the pathway items that are significantly enriched in the significantly differentially expressed metabolites compared with the whole background. p-value in the metabolic pathway is the significance of the metabolic pathway enrichment, and select the significantly enriched pathway to draw the bubble chart. We draw the bubble chart with P < 0.05 as the standard, and we take the pathways with Rich factor>0.1 and dot size > 2 as IPF, CTD-ILD and other interstitial lung diseases as the most relevant metabolic pathways. We can find that the most relevant metabolic pathways in IPF mainly include choline metabolic pathway, metabolic pathway and linoleic acid metabolic pathway in cancer. The most relevant metabolic pathways of CTD-ILD include choline metabolic pathway and retrograde neural signal metabolic pathway in cancer. The most relevant metabolic pathways in other interstitial lung diseases also include choline metabolic pathway and retrograde neural signal metabolic pathway in cancer. Therefore, we can find that there is no difference in the most relevant metabolic pathways between CTD-ILD group and other groups, so the metabolic pathways and linoleic acid metabolic pathways can be regarded as the most relevant metabolic pathways of IPF, which can be distinguished from other interstitial lung diseases. The metabolites involved in the metabolic pathways are L-Glutamate, PE(18:3(6Z,9Z,12Z)/P-18:0) and PC (18) 1z)/20: 2 (11z, 14z)), in linoleic acid metabolism pathway, there are three differential metabolites, namely 12,13-EpOME, 13S-HODE and PC (18: 2 (9z, 12Z)/20:2(11Z,14Z)). Figures 8.1, 8.2 and 8.3, and tables 6.1, 6.2 and 6.3.

**Fig. 8.1.**
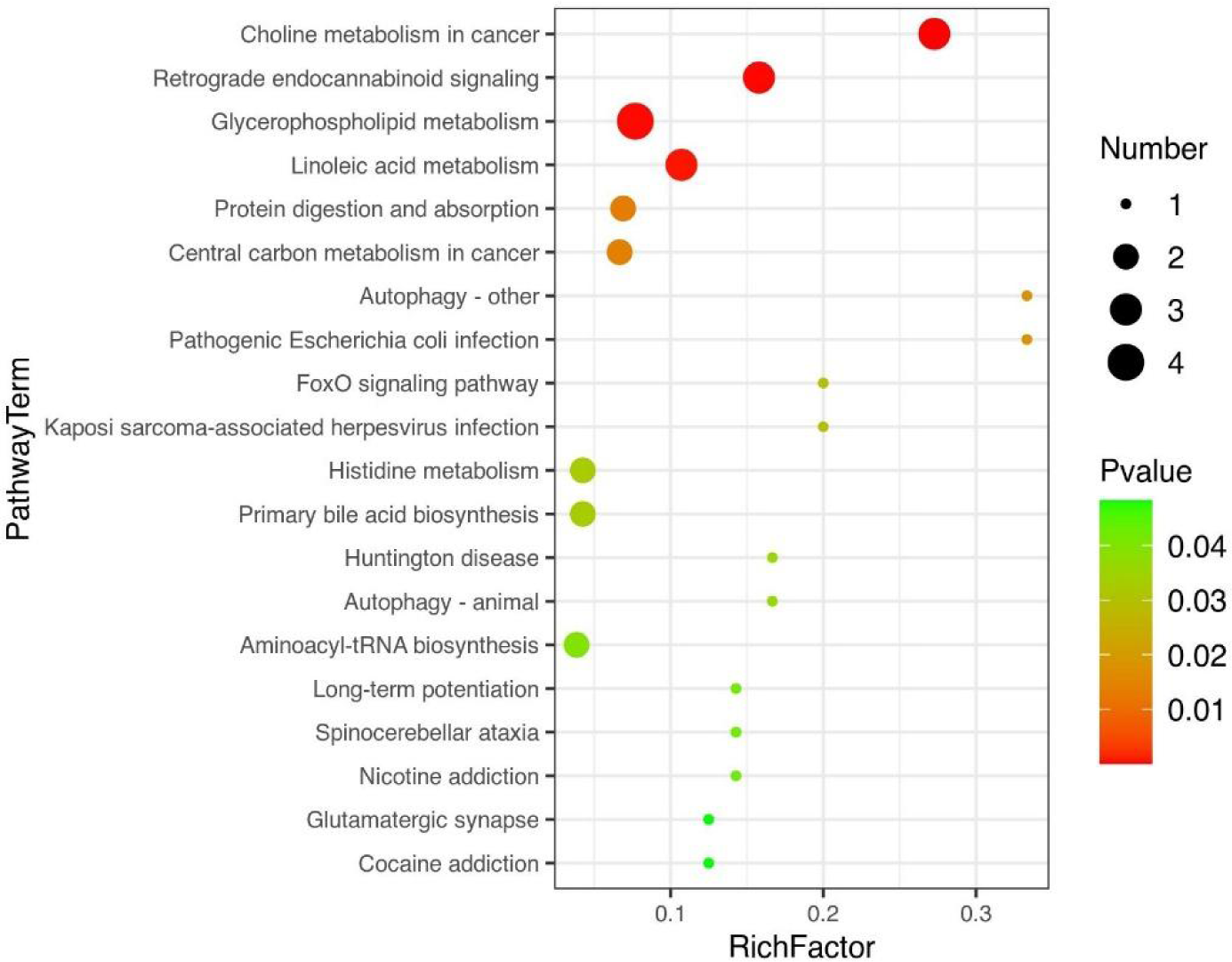
Bubble charts of IPF group and healthy control group with P < 0.05

**Fig. 8.2.**
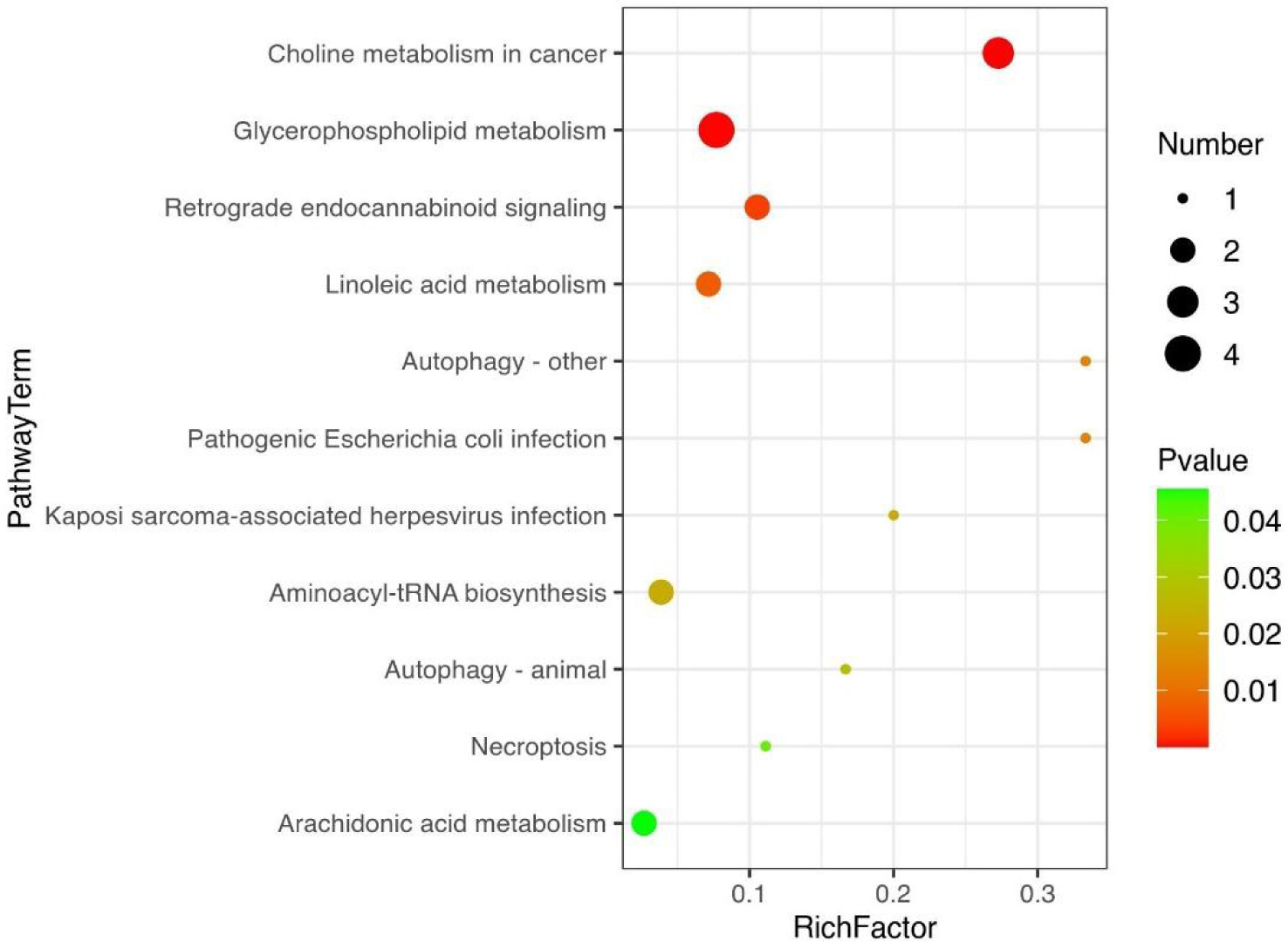
Bubble charts of CTD-ILD group and healthy control group with P < 0.05

**Fig. 8.3.**
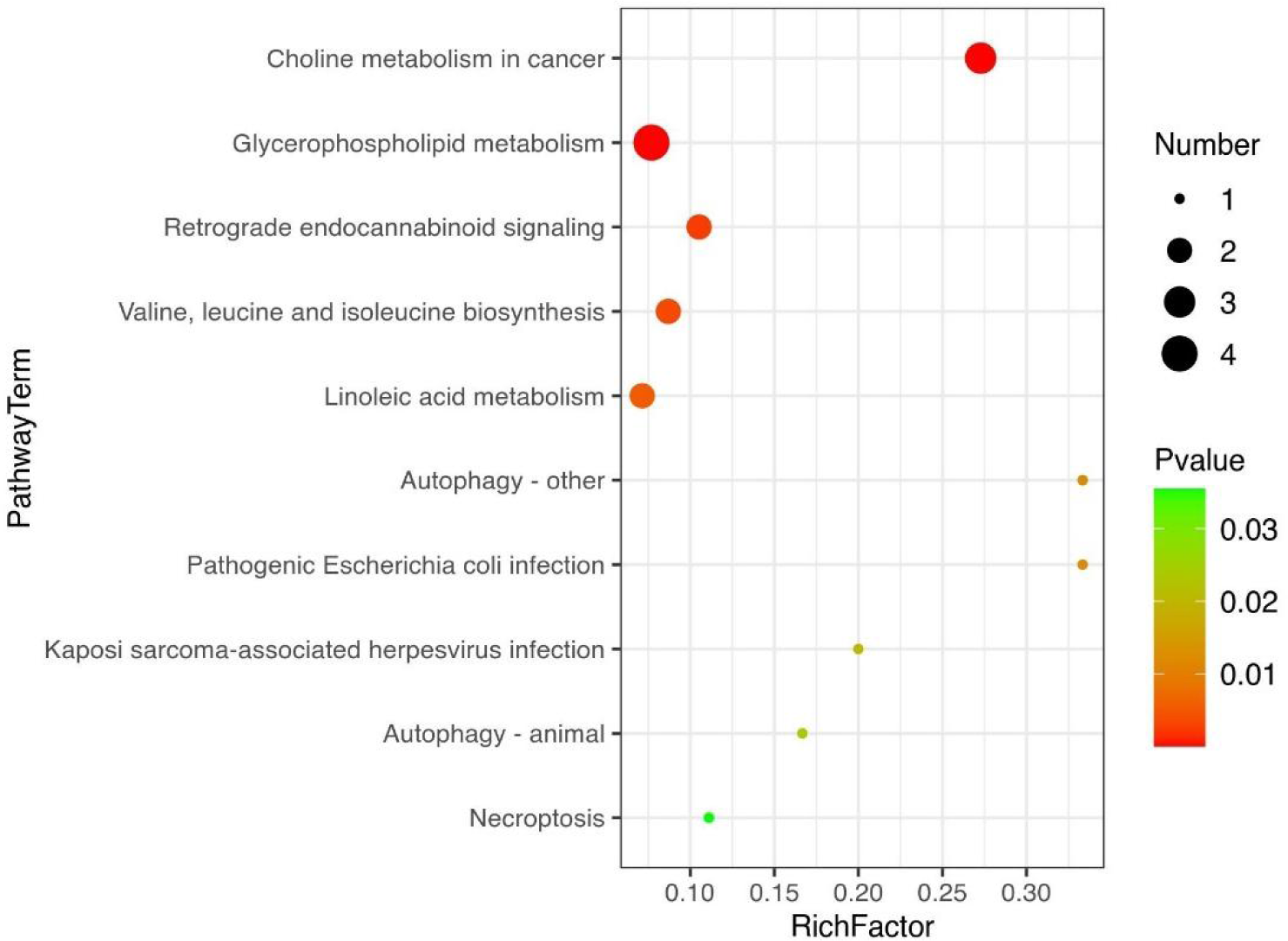
Bubble charts of other groups and healthy control group P < 0.05

**Fig. 9.**
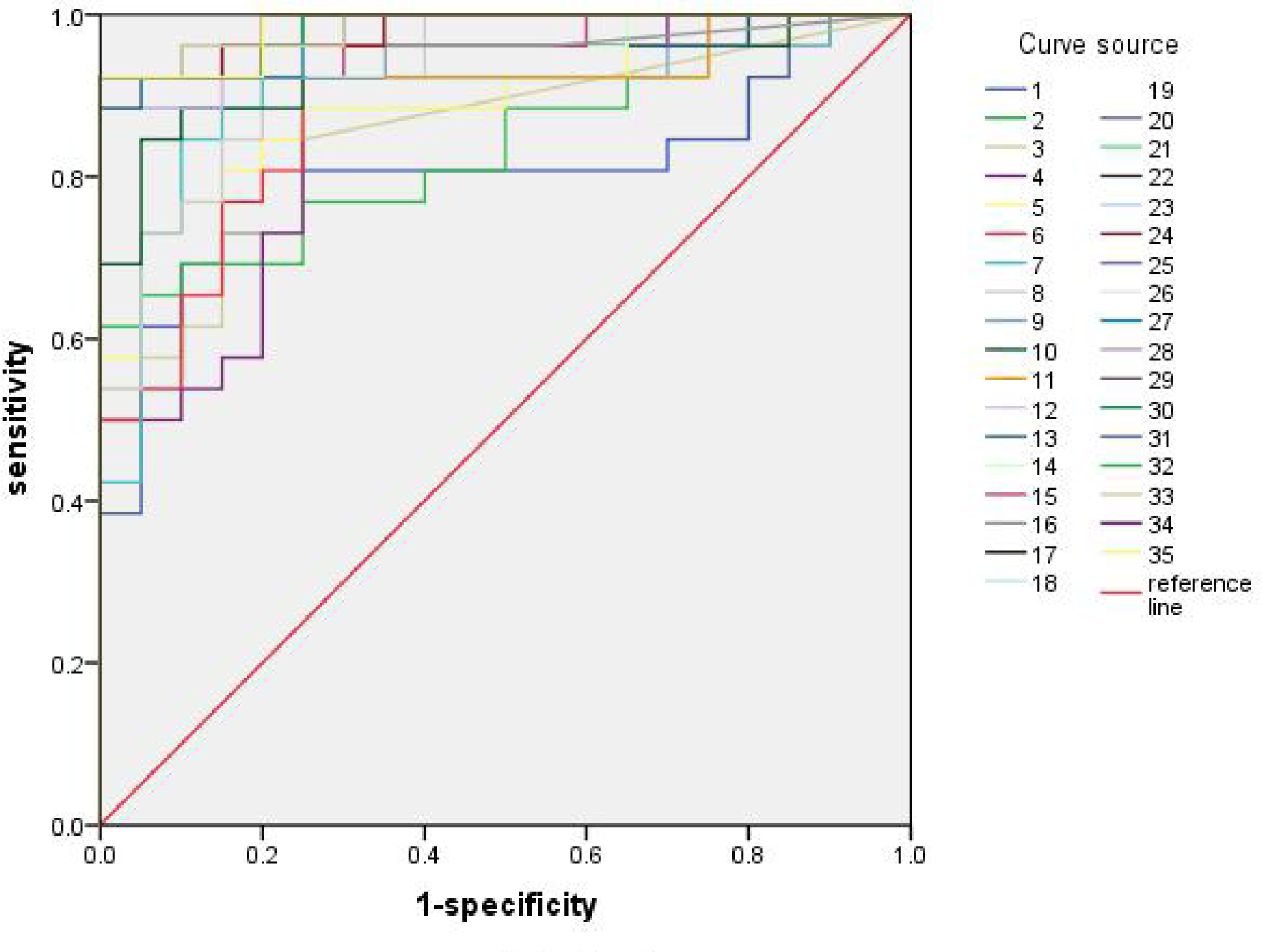
ROC curve of differential metabolites between IPF and healthy control group

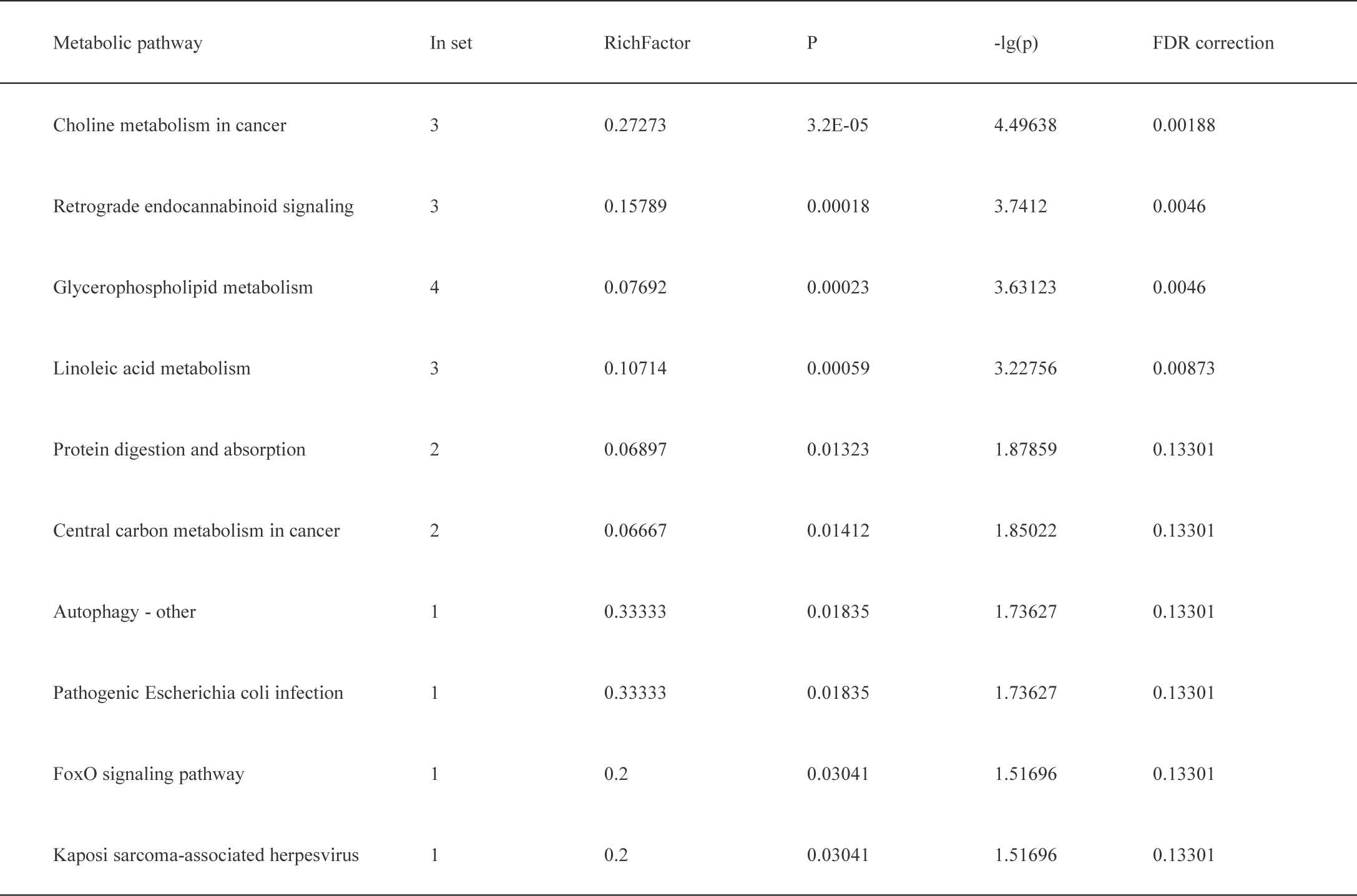
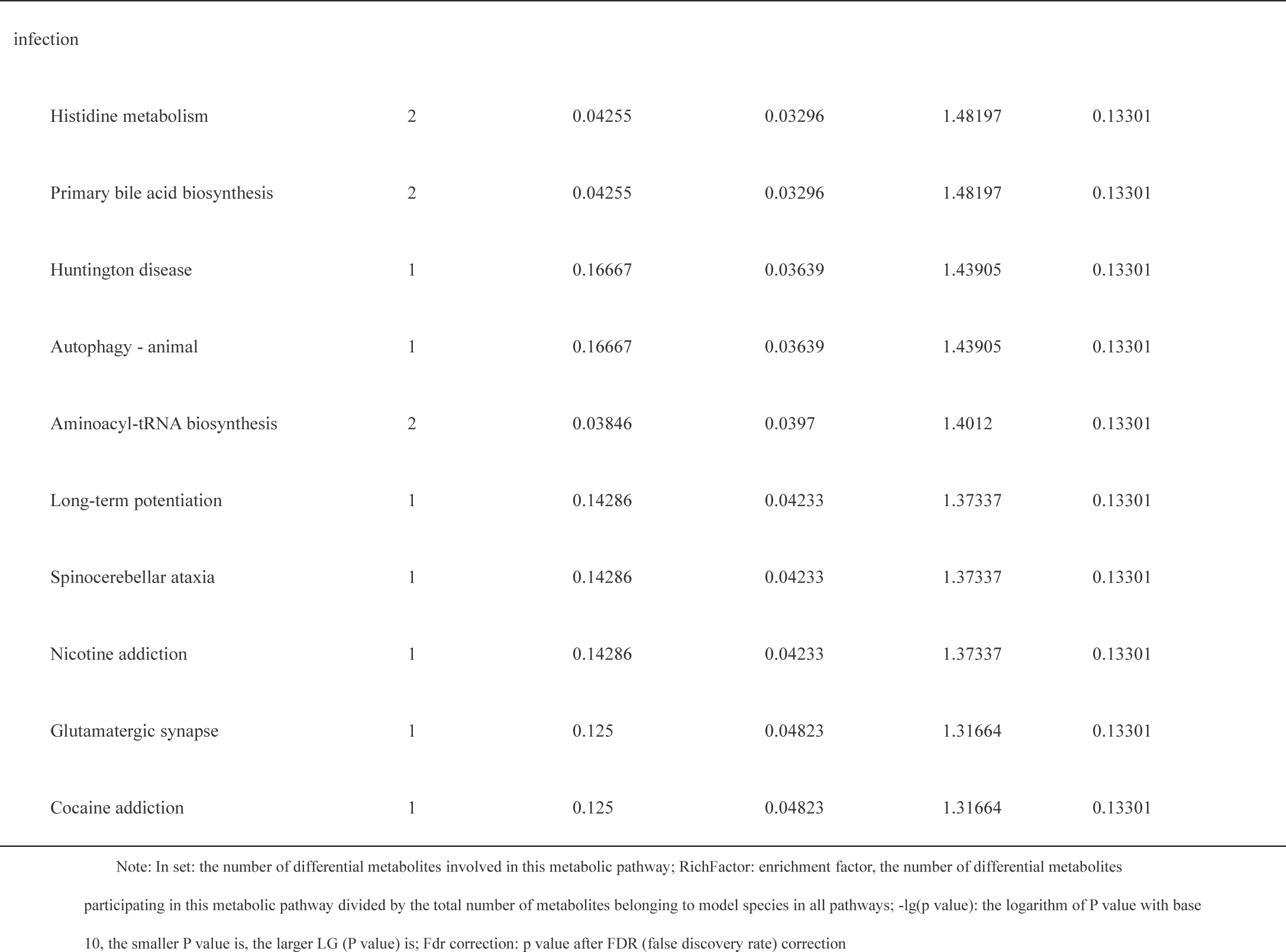

**Table 6.2.** Metabolic pathway analysis results of serum differential metabolites in CTD-ILD group and healthy control group

| Metabolic pathway | In set | RichFactor | P-value | -lg(p-value) | FDR correction |
| --- | --- | --- | --- | --- | --- |
| Choline metabolism in cancer | 3 | 0.27273 | 3.2E-05 | 4.49638 | 0.00188 |
| Retrograde endocannabinoid signaling | 3 | 0.15789 | 0.00018 | 3.7412 | 0.0046 |
| Glycerophospholipid metabolism | 4 | 0.07692 | 0.00023 | 3.63123 | 0.0046 |
| Linoleic acid metabolism | 3 | 0.10714 | 0.00059 | 3.22756 | 0.00873 |
| Protein digestion and absorption | 2 | 0.06897 | 0.01323 | 1.87859 | 0.13301 |
| Central carbon metabolism in cancer | 2 | 0.06667 | 0.01412 | 1.85022 | 0.13301 |
| Autophagy - other | 1 | 0.33333 | 0.01835 | 1.73627 | 0.13301 |
| Pathogenic Escherichia coli infection | 1 | 0.33333 | 0.01835 | 1.73627 | 0.13301 |
| FoxO signaling pathway | 1 | 0.2 | 0.03041 | 1.51696 | 0.13301 |
| Kaposi sarcoma-associated herpesvirus infection | 1 | 0.2 | 0.03041 | 1.51696 | 0.13301 |
| Histidine metabolism | 2 | 0.04255 | 0.03296 | 1.48197 | 0.13301 |

Note: In set: the number of differential metabolites involved in this metabolic pathway; RichFactor: enrichment factor, the number of differential metabolites participating in this metabolic pathway divided by the total number of metabolites belonging to model species in all pathways; -lg(p value): the logarithm of P value with base 10, the smaller P value is, the larger LG (P value) is; Fdr correction: p value after FDR correction

**Table 6.3.**
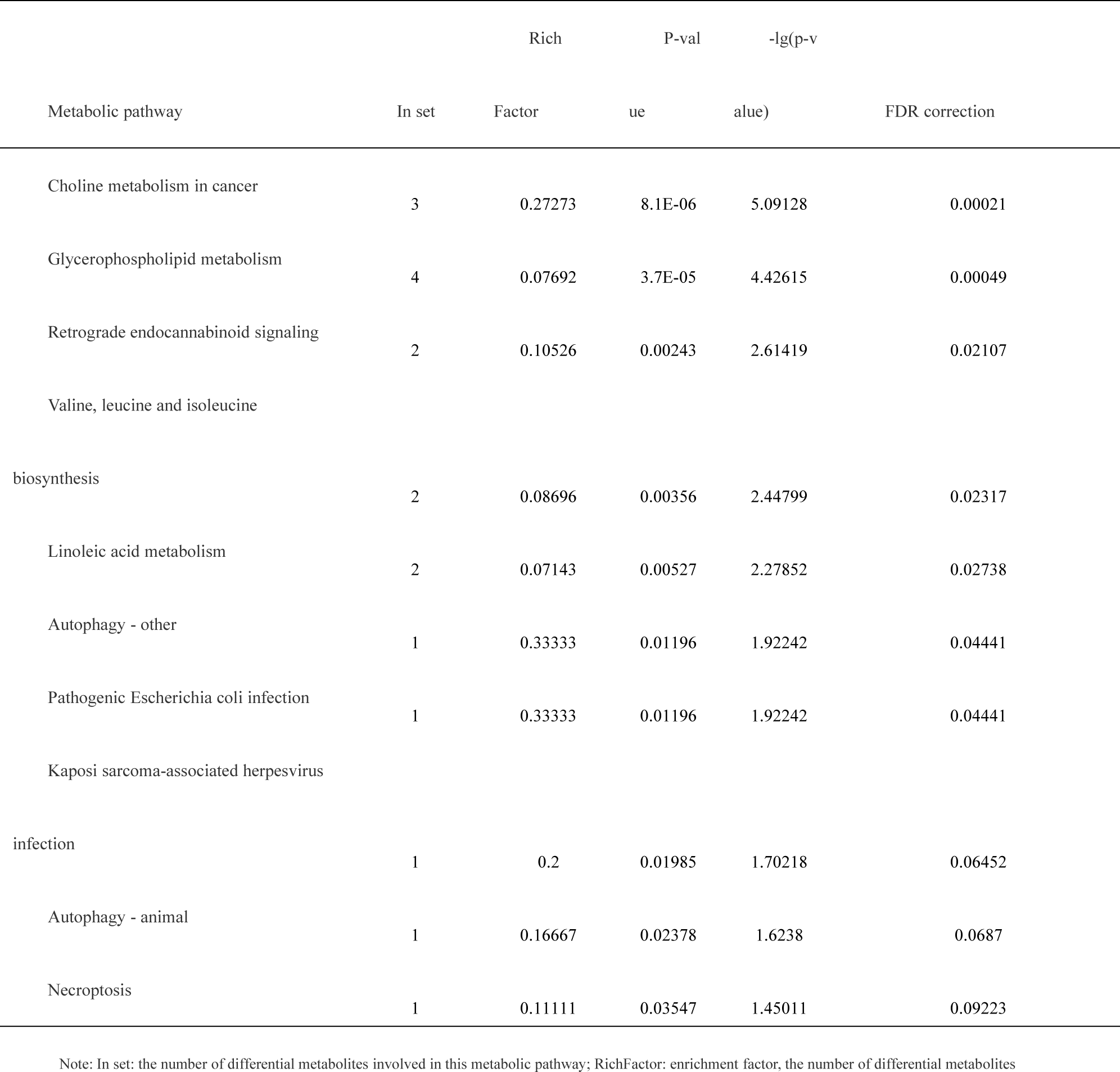
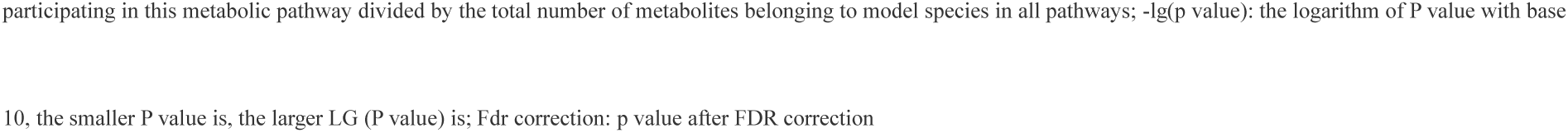
Analysis results of metabolic pathways of serum differential metabolites in other groups and healthy control groups

| Metabolic pathway | In set | Rich<br>Factor | P-val<br>ue | -lg(p-v<br>alue) | FDR correction |
| --- | --- | --- | --- | --- | --- |
| Choline metabolism in cancer | 3 | 0.27273 | 8.1E-06 | 5.09128 | 0.00021 |
| Glycerophospholipid metabolism | 4 | 0.07692 | 3.7E-05 | 4.42615 | 0.00049 |
| Retrograde endocannabinoid signaling | 2 | 0.10526 | 0.00243 | 2.61419 | 0.02107 |
| Valine, leucine and isoleucine<br>biosynthesis | 2 | 0.08696 | 0.00356 | 2.44799 | 0.02317 |
| Linoleic acid metabolism | 2 | 0.07143 | 0.00527 | 2.27852 | 0.02738 |
| Autophagy - other | 1 | 0.33333 | 0.01196 | 1.92242 | 0.04441 |
| Pathogenic Escherichia coli infection | 1 | 0.33333 | 0.01196 | 1.92242 | 0.04441 |
| Kaposi sarcoma-associated herpesvirus<br>infection | 1 | 0.2 | 0.01985 | 1.70218 | 0.06452 |
| Autophagy - animal | 1 | 0.16667 | 0.02378 | 1.6238 | 0.0687 |
| Necroptosis | 1 | 0.11111 | 0.03547 | 1.45011 | 0.09223 |

Note: In set: the number of differential metabolites involved in this metabolic pathway; RichFactor: enrichment factor, the number of differential metabolites
participating in this metabolic pathway divided by the total number of metabolites belonging to model species in all pathways; $-\lg(p \text{ value})$ : the logarithm of P value with base
10, the smaller P value is, the larger LG (P value) is; Fdr correction: p value after FDR correction

### 2.7 ROC curve

According to the analysis of differential metabolites, in IPF group and healthy control group, we screened out 35 differential metabolites with AUC>0.80 as the standard, among which 3 metabolites with the highest AUC were 3- sulfodeoxycholic acid, 20,22-dihydrodigitalis glycoside, () 14 (15)-EET-Si, and the AUC values were all 0.988. L- acetyl carnitine, 5-hydroxydodecanoate and L- glutamic acid are involved in significant enrichment pathways, among which the ROC curve area of L- glutamic acid is the largest, with AUC value of 0.921, cutoff value of 40,145.490, sensitivity of 80%, specificity of 96%, and participation in metabolic pathways.

## 3 Discussion

Clinically, the differential diagnosis of ILD diseases has always been a difficult problem. Including nearly 300 kinds of diseases, ILD can be divided into idiopathic interstitial pneumonia (IIP), autoimmune ILD(CTD-ILD, vasculitis, IPAF), granulomatous ILD, occupational environment-related ILD and other ILD types according to the etiology. Pathological manifestations of the disease can be divided into different types, such as inflammation, fibrosis, and coexistence of inflammation and fibrosis. In recent years, the concepts of fibrotic ILD (fILD) ILD(progressive fibrosing ILD (PF-ILD) have also been paid attention to. Based on the previous retrospective investigation by the research team, the main diseases of ILD diagnosed in our department are autoimmune ILD, IPF, sarcoidosis, other ILD types and occupational environment-related ILD. Due to the relatively small number of cases of various diseases included in ILD in this study, the main subgroups of ILD were combined and analyzed, including IPF, CTD-ILD (including vasculitis and IPAF) and other ILDs. IPF is a rare disease, and it is also the most common disease in IIP. The prevalence rate is about 8.2 per 100,000, the onset age is between 60 and 70 years old, and the incidence rate of males is high, and tobacco is the risk factor of IPF [19]. The 5-year survival rate of IPF is 20%-40%, which is second only to many malignant tumors. In acute exacerbation, the hospitalization rate is more than 50%, and the median survival time is 3-4 months [20, 21]. Patients without systemic anti-fibrosis treatment or lung transplantation have a survival time of about 3.5-4.5 years [22], and the prognosis is extremely poor. The molecular pathological mechanism of IPF includes the activation of fibroblasts, alveolar epithelial dysfunction, oxidative stress, vascular modification, gene remodeling and the wrong repair mechanism of aging, etc, but it is still unclear at present [23]. At present, the diagnosis of IPF still has a strong exclusive diagnostic nature. Even if the imaging is consistent with typical interstitial pneumonia (UIP), other types of ILD that can cause UIP still need to be excluded for differentiation. In addition, the diagnosis of IPF is often delayed clinically, and it takes 1.5 years from symptoms to diagnosis of IPF, which is not conducive to early intervention and prognosis of patients [20]. Therefore, exploring the diagnosis and differential diagnosis markers of ILD and IPF is of great significance for patients to get timely diagnosis, make effective and scientific treatment decisions, and improve the prognosis of patients.

With the research of genomics in recent years, metabolomics is a research hotspot after genomics, transcriptomics and proteomics, and it is an important part of biological system. By detecting the changes of metabolic spectrum, we can study the metabolism of the whole organism, further reveal the changes of endogenous substances and gene levels, and it has been widely used in many fields. Related literatures have expounded the related metabolism of idiopathic pulmonary fibrosis, including pathogenesis, biomarkers and therapeutic targets [24].

In this study, serum samples of IPF patients, CTD-ILD patients, other ILD patients and healthy controls were detected by LC-MS technology, hoping to explore potential biomarkers of different interstitial lung diseases by metabonomics. Through LC-MS analysis of 90 collected serum samples, the differential metabolites among different groups were found. Through OPLS-DA, T test and correlation analysis, the differential metabolites between the two groups were finally determined, including 193 metabolites in IPF group and healthy control group, 188 metabolites in CTD-ILD group and healthy control group, and 169 metabolites in other kinds of interstitial lung diseases and healthy control group. Through the correlation analysis of the first 50 most obvious metabolites, we found the correlation between the different metabolites of each group and the healthy control group. Then, the enrichment analysis of metabolic pathways of differential metabolites showed that IPF, CTD-ILD and other interstitial lung diseases were the most relevant metabolic pathways, and IPF had three main metabolic pathways, namely choline metabolic pathway, metabolic pathway and linoleic acid metabolic pathway in cancer. There are two metabolic pathways most related to CTD-ILD, including choline metabolic pathway and retrograde neural signal metabolic pathway in cancer. The most relevant metabolic pathways in other interstitial lung diseases also include choline metabolic pathway and retrograde neural signal metabolic pathway in cancer. Comprehensive analysis shows that choline metabolic pathway in cancer is a common metabolic pathway in interstitial lung diseases, while IPF has unique metabolic pathway and linoleic acid metabolic pathway. In IPF group and healthy controls, through the analysis of differential metabolites, we screened out 35 differential metabolites with AUC>0.80 as the standard, among which 3 metabolites with the highest AUC were 3- sulfodeoxycholic acid, 20,22-dihydrodigitalis glycoside, () 14 (15)-EET-Si, and the AUC values were all 0.988. 22- dihydrodigitalis glycoside belongs to both lipid and lipid molecules, and the diagnostic significance of lipid molecules in idiopathic pulmonary fibrosis further explained in this study is consistent with previous research results. L- acetyl carnitine, 5-hydroxydodecanoate and L- glutamic acid are involved in significant enrichment pathways, among which the ROC curve area of L- glutamic acid is the largest, with AUC value of 0.921, cutoff value of 40,145.490, sensitivity of 80%, specificity of 96%, and participation in metabolic pathways.

This study is mainly to explore new biomarkers of IPF in the metabolic pathway with significant enrichment pathway, in which the AUC value of L- glutamic acid is the largest. L- glutamic acid is an acidic amino acid, one of the basic amino acids for nitrogen metabolism in organisms, and one of the main components of protein. As shown in fig. 10, in the metabolic pathway, L- glutamic acid participates in the synaptic vesicles cycle and promotes the energy metabolism of the body. L- glutamic acid can generate α-ketoglutaric acid through oxidative deamination, which participates in citric acid cycle and further participates in energy metabolism. L- glutamic acid is also a sugar-producing amino acid, which is isomerized into sugar through alanine-glucose cycle and participates in ornithine cycle to synthesize urea for metabolism. And ammonia and L- glutamic acid are synthesized into glutamine under the action of glutamine synthetase, transported to liver and kidney through blood, and hydrolyzed by glutamine enzyme to generate glutamic acid and ammonia, which further affects ornithine circulation and metabolism of arginine and ornithine. Glutamine is closely related to the formation of fibrosis. In fibrotic cells, the activation of TGF-β1 increases the decomposition of glutamine, which is mainly related to the synthesis of glutamine subtype 1(GLS1) induced by TGF-β1, and this process depends on protein kinases activated by SMAD3 and p38 mitogen [25] The latest literature points out that L- arginine is related to pulmonary fibrosis [26]. Arginine is an essential amino acid and a basic component of protein synthesis. Metabolites are involved in many ways, mainly involved in energy generation and ornithine cycle. Ornithine is one of its components. Ornithine is converted into spermidine and spermine through decarboxylation. These two ornithine derivatives are important substances for regulating cell growth and promoting cell proliferation. Ornithine can also be converted into proline and semi-proline under certain conditions, regulating cell redox, and is an important component of fibrotic collagen, which further shows that arginine and ornithine are of great significance in the pathogenesis of IPF [27]. In a study, the metabolic changes of lung tissues involved in the pathogenesis of IPF were analyzed by gas chromatography-mass spectrometry, and it was found that the concentration of proline in lung tissues of IPF patients was higher than that of healthy controls [28]. In this study, the content of glutamic acid in IPF group is higher than that in healthy control group, and it can be further speculated that it will affect the production of proline, suggesting that glutamic acid may promote the formation of pulmonary fibrosis through metabolic pathway in IPF.

**Fig. 10.**
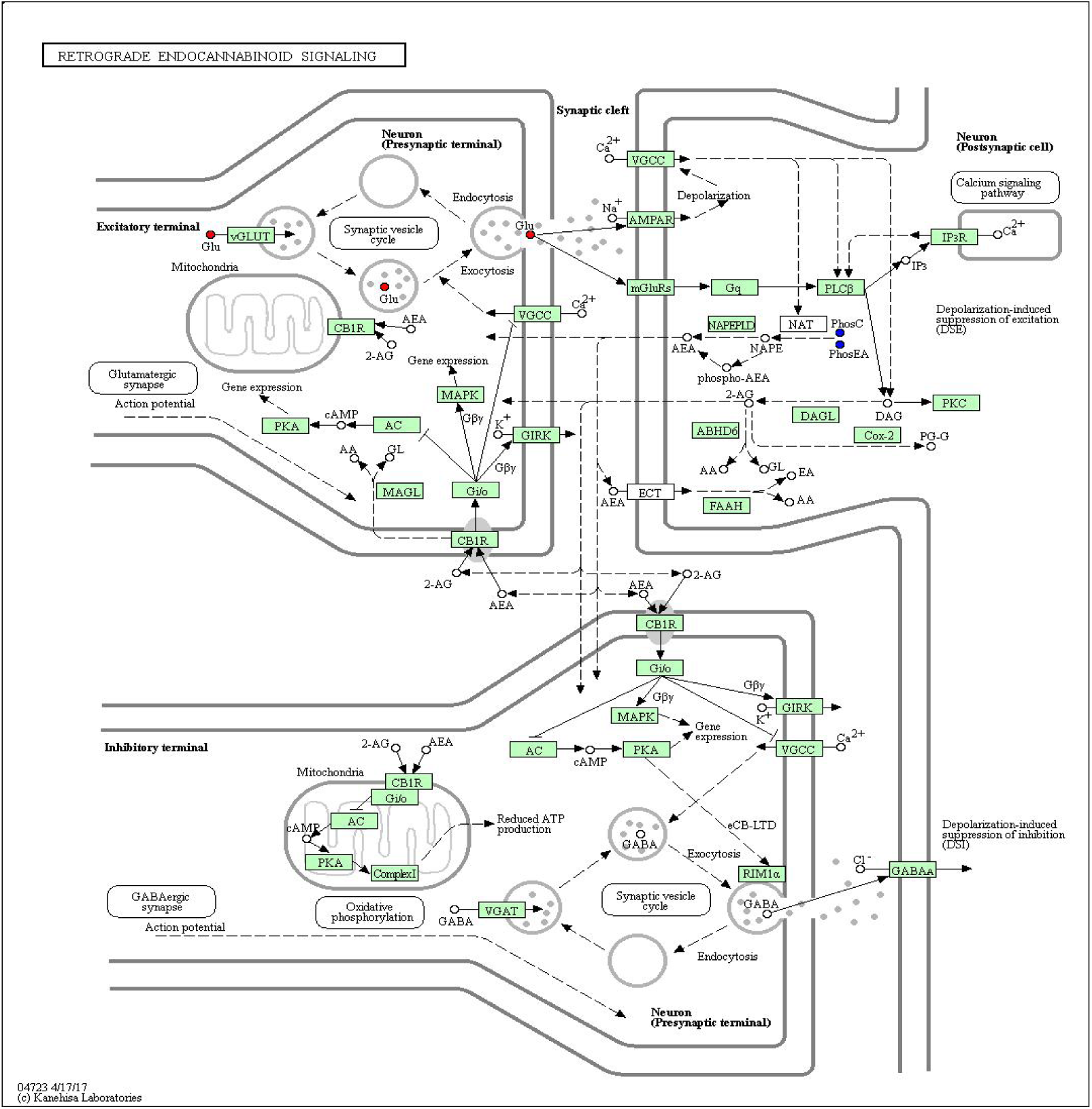
metabolic pathway diagram

There are three kinds of differential metabolites in linoleic acid metabolic pathway, namely 12,13 epoxy octadecenoic acid, 13 thiohydroxy octadecadienoic acid and phosphatidylcholine, which belong to monounsaturated fatty acids, and phosphatidylcholine belongs to phospholipids. Phosphatidylcholine is synthesized through diacylglycerol pathway, which is activated by choline to generate CTD- choline, and then condensed with diacylglycerol to form phosphatidylcholine. Is an important component of cell membrane, which plays an important role in the process of cell proliferation and differentiation, and type II alveolar epithelial cells can synthesize special phosphatidylcholine, which is one of the main components of alveolar surfactant, and has the function of reducing alveolar surface tension, which is beneficial to the expansion of alveoli. Alveolar surfactant is closely related to pulmonary fibrosis [29]. The level of phosphatidylcholine is regulated by various phospholipases, such as phospholipase A and phospholipase D, which catalyze the hydrolysis of phosphatidylcholine [30]. As shown in Figure 11, choline phosphatide is metabolized to synthesize linoleic acid salt, which further forms unsaturated fatty acids and participates in linoleic acid metabolism pathway. Some endogenous substances, such as 9 and 13 hydroxyl 10e, 12z octadecadienoic acid and its metabolites, are generated by oxidation of linoleic acid, and under the action of heavy metal ions, γ-linoleic acid is oxidized to further generate arachidonic acid, which has anti-inflammatory effect. In this study, the expression of 12,13-EpOME and 13S-HODE in IPF group is lower than that in healthy control group, so it can be inferred that the anti-inflammatory ability of IPF patients is decreased. In a recent lipidomics study on plasma of IPF patients, 62 lipids were identified, but it was not clear which was a potential biomarker [31]. Lipid-related research on bleomycin-induced pulmonary fibrosis rats shows that the synthesis of phospholipids and sphingolipid lipids is enhanced, and the matrix remodeling is increased on the basis of decomposition [32]. In this study, phosphatidylcholine is down-regulated in linoleic acid metabolic pathway, which indicates that the abnormality of linoleic acid metabolic pathway may affect the anti-inflammatory mechanism of the body in the pathogenesis of IPF.

**Fig. 11.**
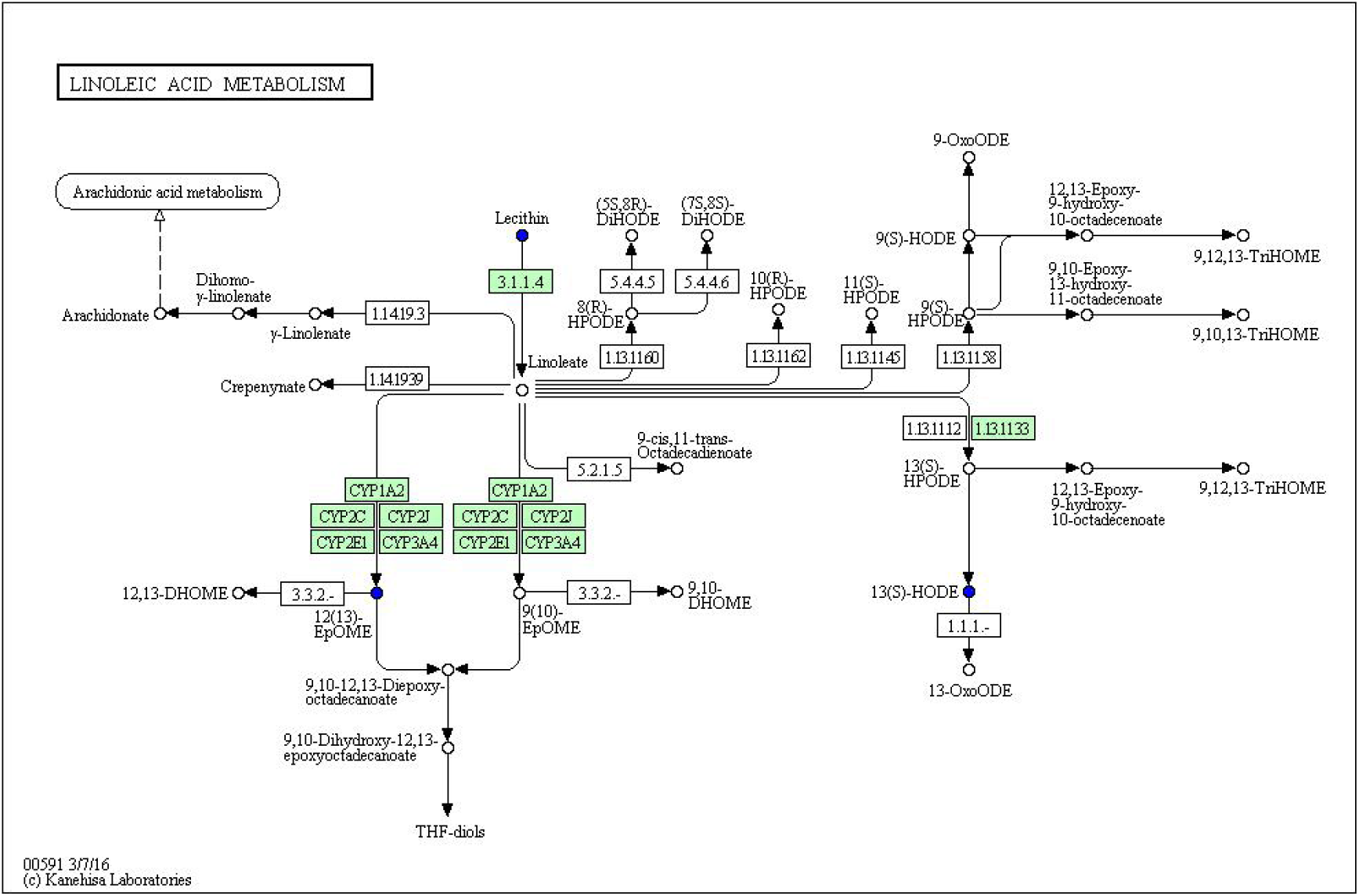
Linoleic acid metabolic pathway

## 4 Conclusion

n this study, the serum samples of IPF patients, CTD-ILD patients, other interstitial lung diseases and healthy controls were metabonomics analyzed by LM-MS technology. After analysis and identification, the following conclusions were drawn.

There are differences in metabolites in different treatment groups, including 193 metabolites in IPF group and healthy control group, 188 metabolites in CTD-ILD group and healthy control group, and 169 metabolites in other interstitial lung diseases and healthy control group.

Through the enrichment analysis of metabolic pathways of differential metabolites, it is found that IPF has three main metabolic pathways, namely choline metabolic pathway, metabolic pathway and linoleic acid metabolic pathway in cancer; There are two metabolic pathways most related to CTD-ILD, including choline metabolic pathway and retrograde neural signal metabolic pathway in cancer. The most relevant metabolic pathways in other interstitial lung diseases also include choline metabolic pathway and retrograde nerve signal metabolic pathway in cancer. Through comprehensive analysis, it is found that there are two metabolic pathways different from other groups for IPF, namely metabolic pathway and linoleic acid metabolic pathway, which are of great significance in the pathogenesis of IPF.

3. For IPF, ROC curve analysis shows that there are 35 kinds of differential metabolites with AUC greater than 0.80, and the metabolites with the highest diagnostic value are 3- sulfodeoxycholic acid, 20,22-dihydrodigitalis glycoside and () 14 (15)-EET-Si, with AUC values of 0.985 and sensitivity of 100%.

## The second chapter is a preliminary study of serum GDF-15 in ILD patients based on metabonomics

### 1 research materials and methods

research object and grouping The subjects were the same as above, and the general clinical data, lung function test data and serum samples were collected. Serum levels of type I collagen, KL-6, IL-1β, TNF-α, IGF-1 and GDF-15 were detected by Enzyme linked immunosorbent assay (ELISA) in patients with ILD and healthy controls. The patient with ILD was diagnosed as ILD through multidisciplinary discussion of clinical, imaging and pathology, and the disease type was clear, and the enrolled patients had signed the informed consent form. Inclusion criteria and exclusion criteria are as shown in Chapter I, 1.4.1.

### 2 data collection

Collecting pulmonary function indicators: using Jaeger brand pulmonary function meter to detect pulmonary function, repeating the measurement for 3 times to get the best value, and interpreting the results. Pulmonary function indexes: including forced vital capacity FVC, total lung capacity TLC, vital capacity VC, carbon monoxide dispersion DLco and unit dispersion DLCO/VA. Record the measured value and the percentage of measured value/predicted value respectively.

### 3 Serum detection indicators

Collect serum samples as shown in Chapter 1.4.2, and conduct ELISA detection uniformly. Freeze-thaw at 4℃ 24 hours before detection, and use ELISA detection kit uniformly to detect type I collagen, KL-6, IL-1β, TNF-α, IGF-1 and GDF-15.

### 4 experimental equipment and reagents

Main equipment: DH36001B electrothermal constant temperature incubator (Tianjin Tester Company), ELX-800 microplate reader (American BIOTEK Company), Proline micropipette (Suzhou BIOHIT Company).

### 5 Main reagents

Human Escherichia coli (Collagen Type I) ELISA kit (Wuhan Fein Company, China, item number: EH7533), Human MUC-1(Mucin-1) ELISA Kit (Wuhan Fein Company, China, item number: EH0406), leukocyte −1β(IL-1β)ELISA detection kit (Shanghai Xin Tumor necrosis factor α(TNF-α)ELISA kit (Shanghai Xinaosheng Biotechnology Company, article number: WLE05), Human IGF-1 ELISA Kit (China Lianke Company, article number: EK1131), Human GDF-15 ELISA Kit (China Lianke Company, article number: EK1100).

### 6 experimental methods

6.1 Reagent preparation: Reagent preparation is carried out according to the instructions of each kit. 1.4.2 operation steps

6.2(1)experimental steps of type I collagen and MUC-1: washing plates, adding samples, incubating, washing, labeling antibodies with biotin, and then carrying out the next step after washing, labeling streptavidin conjugate (SABC) with HRP, adding TMB substrate, adding 90 μl TMB substrate to each well, covering the plate, incubating at 37℃ in the dark for 10-20 minutes, and finally, each well. The adding sequence of the stopping solution should be the same as that of TMB substrate solution, and OD absorbance should be immediately read by microplate reader at 450 nm.

(2)Experimental steps of IL-1β and TNF-α: coat the antibody according to the instructions, take 100 μl of diluted standard samples each, add them into 7 wells in a row of 96-well plates in turn, add 1×PBST to the blank wells, then carry out sample detection, and finally add 50 μ L of TMB stop solution to stop the reaction, and read the absorbance at 450 nm by microplate reader.

(3)Experimental steps of IGF-1 and GDF-15 detection: according to the instructions, wash the plate first, then add samples, then discard the liquid, wash each well with 300 μL washing liquid for 6 times, add 100 μL diluted horseradish peroxidase-labeled streptavidin to each well, seal the plate with sealing film, shake at 300 rpm, incubate at room temperature for 45 minutes, and then wash it for 5 times. Incubate at room temperature for 5-30 minutes in the dark, add 100 μL of stopping solution to each well, stop the reaction, and measure the OD value at the maximum absorption wavelength of 450 nm and the reference wavelength of 570 nm. The calibrated OD value is the measured value of 450 nm MINUS the measured value of 570 nm.

### 7 statistical analysis

Continuous variables that conform to normal distribution are expressed as mean standard deviation (x± s). Parameters that do not conform to normal distribution are expressed by median and quartile spacing M(P25, P75). According to normal distribution, T test is used between two groups, single-factor analysis of variance is used for comparison among multiple groups, Mann-Whitney U test is used for comparison between two groups, Kruskal-Wallis rank sum test is used for comparison among multiple groups, and GraphPad Prism8.0 is used for comparison between groups. Two-sided inspection, take α=0.05. Chi-square test was used to compare the classification and rate. P0.05, the difference was statistically significant.

Pearson correlation analysis was used to detect the correlation between serum biomarkers and other factors, and R (version 3.6.2) software was used for correlation heat map. ROC curve was used to analyze the diagnostic efficiency of each index, and the best critical value, sensitivity and specificity were found out.

## 2 Results

### 1.1 Comparison of pulmonary function in each group

For the measured value of DLco, the carbon monoxide dispersion of IPF group is lower than that of other groups, P < 0.05, and the difference is statistically significant. For the dispersion coefficient, IPF group was lower than CTD-ILD group and other groups, P < 0.05, and the difference was statistically significant. The percentage of FVC measured value/predicted value in IPF group is lower than that in other groups, with significant difference (P < 0.05), but there is no difference between IPF group and CTD-ILD group. As for the percentage of measured value/predicted value of DLco, IPF group is lower than other groups, with significant difference, and the difference is statistically significant (P < 0.05). As shown in Table 8。

**Table 7.**
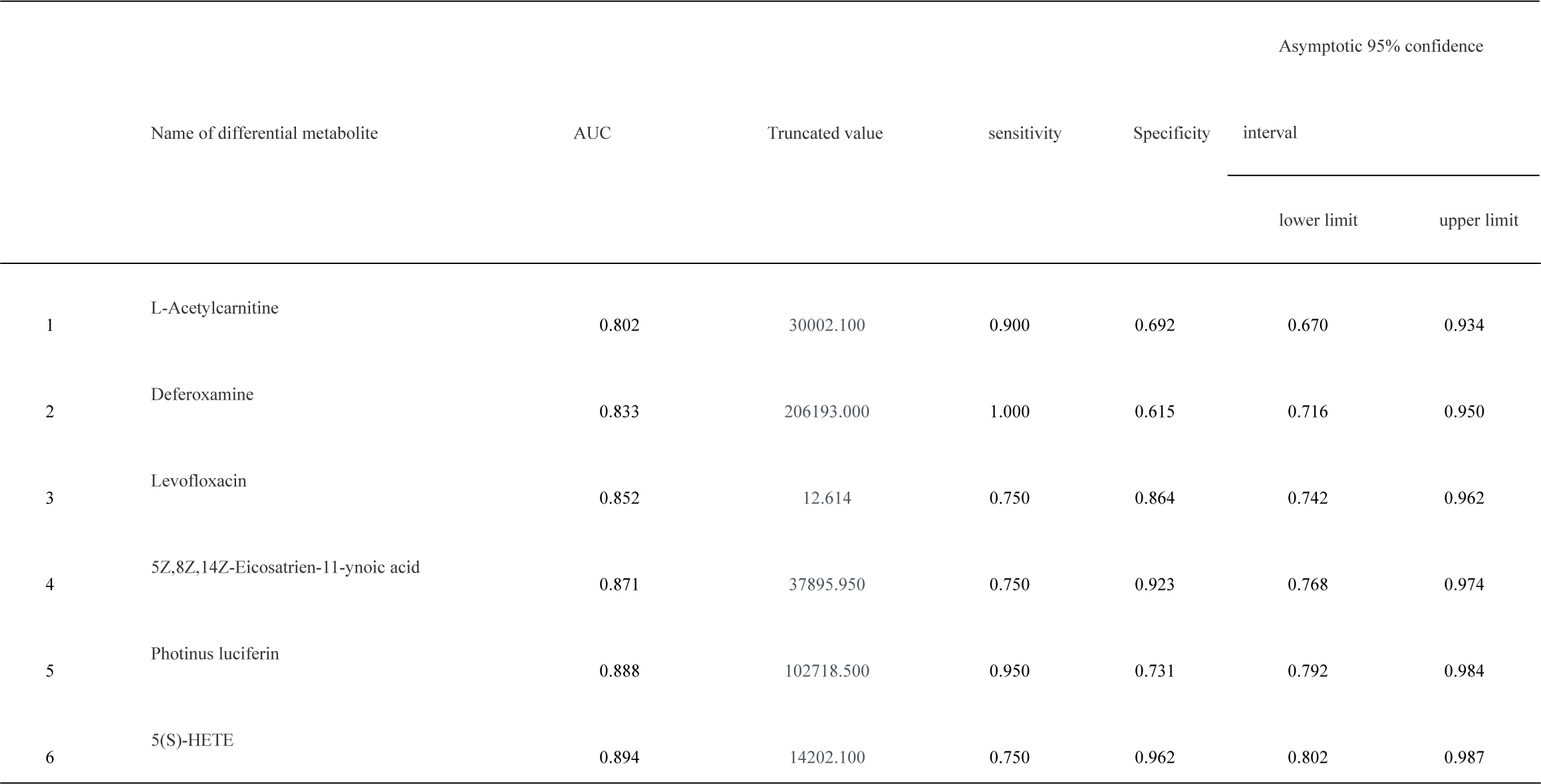
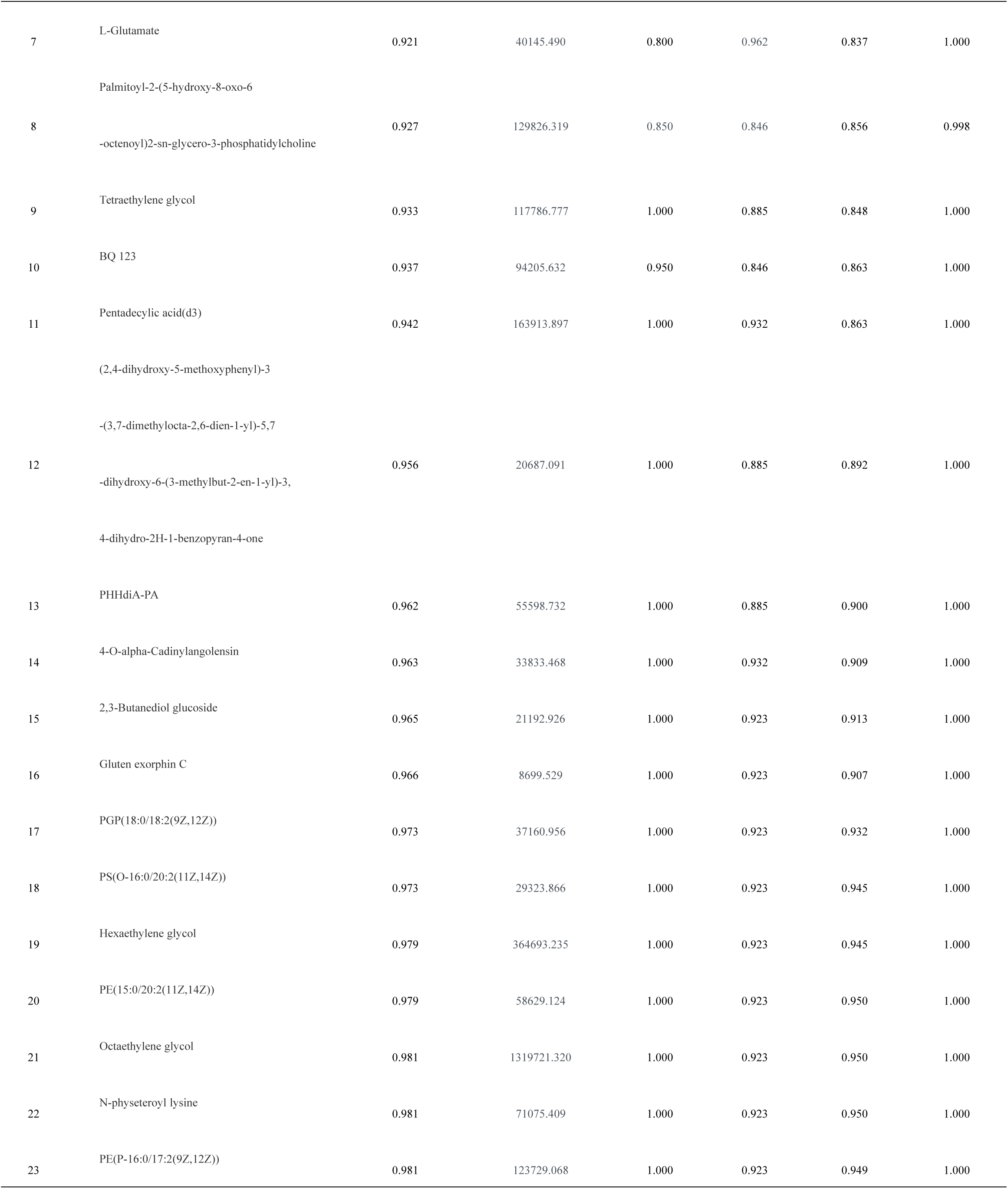
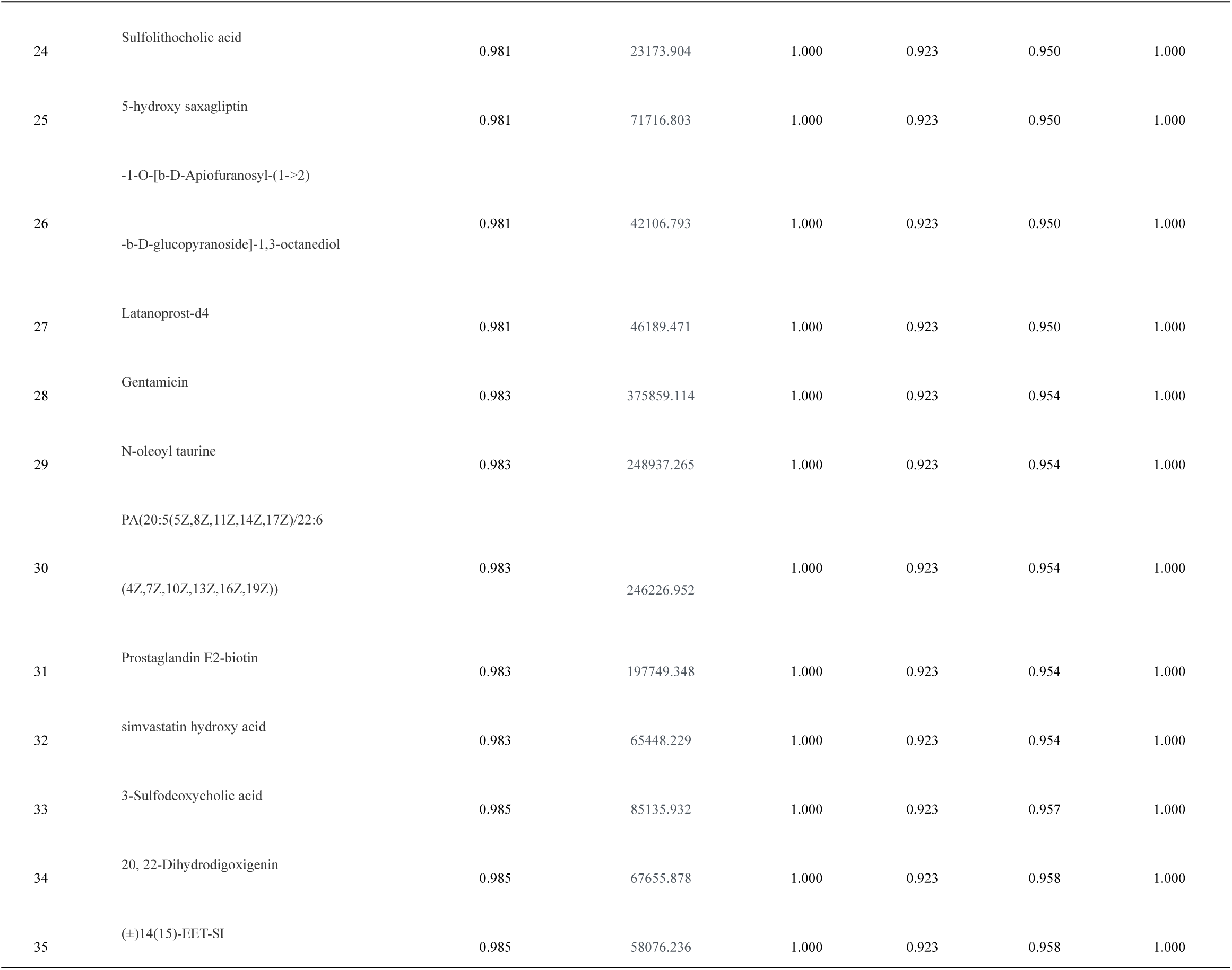
ROC curve area of differential metabolites between IPF group and healthy control group

**Table 8.** Pulmonary function indexes of each group

|  | IPF (n=16) | CTD-ILD (n=11) | Others (n=13) | F value | P value |
| --- | --- | --- | --- | --- | --- |
| FVC measured value | 2.32±0.84 | 2.21±0.76 | 3.10±1.77 | 2.097 | 0.137 |
| TLC measured value | 3.99±0.69 | 3.81±0.81 | 4.21±1.31 | 0.519 | 0.600 |
| VC measured value | 2.36±0.85 | 2.24±0.74 | 2.81±0.90 | 1.578 | 0.220 |
| DLco measured value | 3.48±1.57b | 4.40±1.39 | 5.47±1.55 | 6.193 | 0.005 |
| DLco/VA measured value | 0.95±0.24ab | 1.25±0.37 | 1.31±0.27 | 6.552 | 0.004 |
| FVC Measured value/predicted value percentage | 68.46±19.02b | 77.20±21.84 | 87.18±15.21 | 3.587 | 0.038 |
| TLC Measured value/predicted value percentage | 65.53±11.81b | 76.25±15.56 | 77.77±13.94 | 3.488 | 0.041 |
| VC Measured value/predicted value percentage | 70.97±19.79 | 76.42±21.89 | 85.57±14.13 | 2.179 | 0.127 |
| DLco Measured value/predicted value percentage | 47.42±18.52b | 58.84±20.13 | 67.45±12.04 | 4.947 | 0.012 |
| DLco/VA Measured value/predicted value percentage | 75.58±20.20 | 81.40±17.08 | 84.41±14.88 | 1.441 | 0.250 |

Note: a:CTD-ILD b:Others; FVC: vital capacity; TLC: total lung volume; VC: vital capacity, DLco: total diffusion capacity; DLco/VA: unit dispersion

### 1.2 Comparison of detection indexes in serum

For these six indexes detected in serum, IPF group and CTD-ILD group are higher than the healthy control group, and the difference is statistically significant. For IGF-1, other groups are higher than the healthy control group, and the difference is statistically significant. And these six indexes, although IPF group is slightly higher than CTD-ILD group, have no significant difference. The difference between IPF group and other groups is significant (P < 0.05), as shown in Table 9 and Figure 12. Comparing all ILD with healthy control group, it is found that these six indexes of serum detection are higher than those of healthy control group, and the difference is statistically significant (P < 0.05), as shown in Table 10 and Figure 13. Through the comparative analysis of IPF group, non-IPF group and healthy control group, it is found that the expression of GDF-15 in IPF group and non-IPF group is higher than that in healthy control group, with significant difference, but the difference is not statistically significant between IPF group and non-IPF group, as shown in Table 11 and Figure 14.

**Fig. 12.**
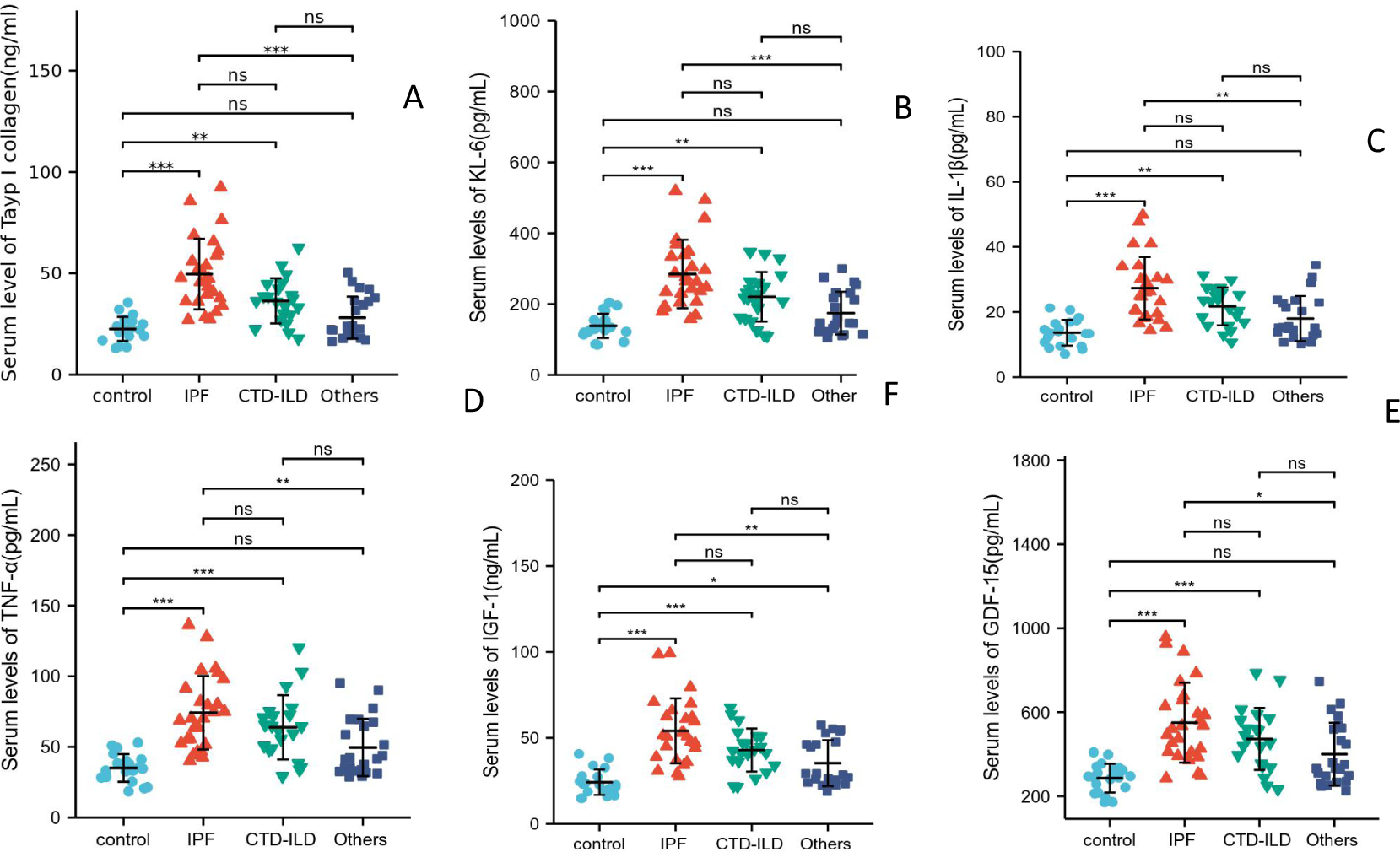
Comparison of serum expression levels of four groups of serum samples Note: ns, P≥0.05; *, P< 0.05; **, P<0.01; ***, P<0.001; Control: healthy control group; IPF: idiopathic pulmonary fibrosis group; CTD-ILD: connective tissue-associated interstitial pneumonia; Others: other interstitial pneumonia; A: the serum level of type I collagen; B: serum level of KL-6; C: the serum level of IL-1β; D: Serum level of TNF-α; E: serum level of IGF-1; F: Serum level of GDF-15

**Fig. 13.**
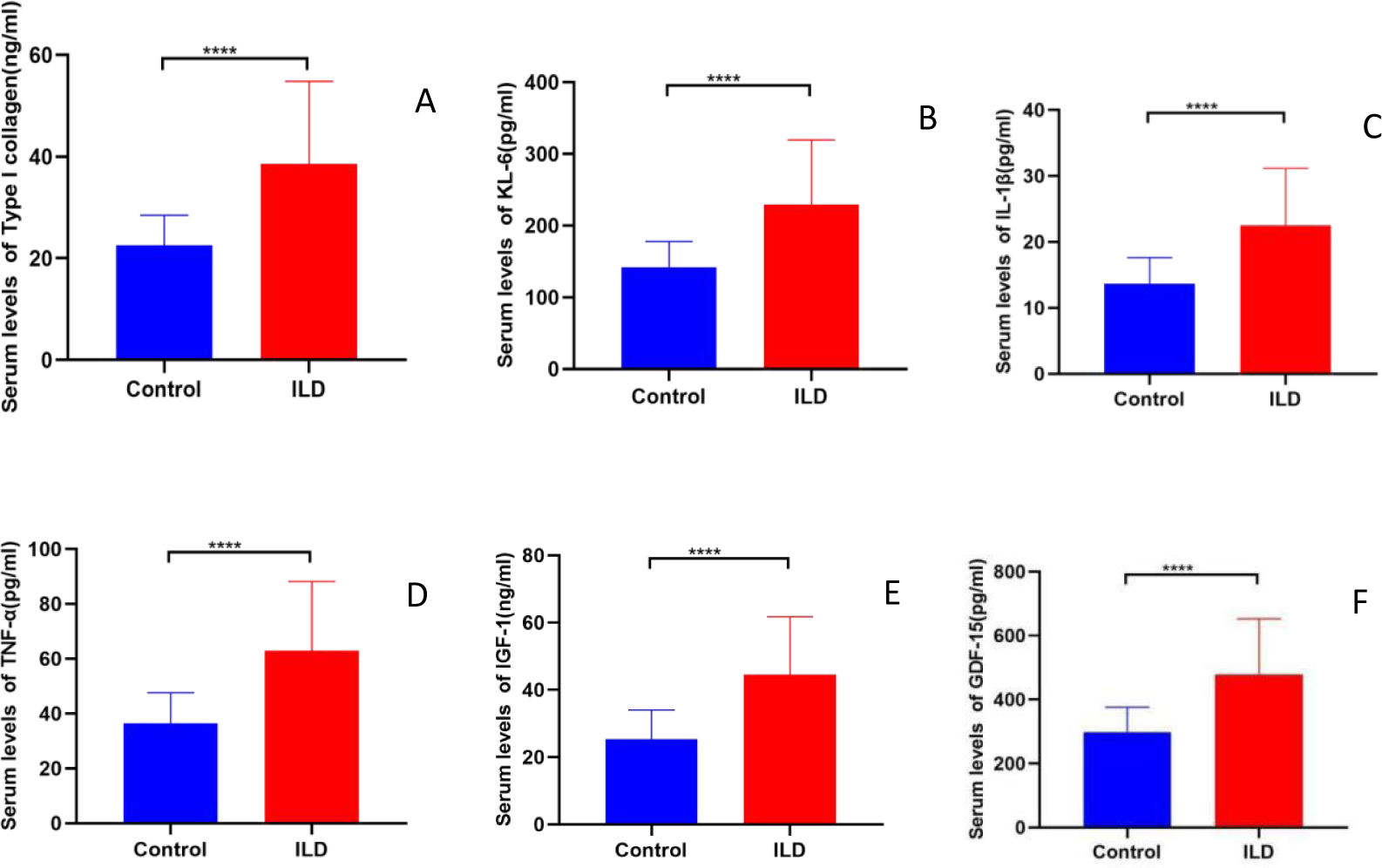
Comparison of serum detection indexes between ILD group and healthy control group Note: * * *: P < 0.0001, control: healthy control group; ILD: interstitial lung disease group; A: the serum level of type I collagen; B: serum level of KL-6; C: the serum level of IL-1β; D: Serum level of TNF-α; E: serum level of IGF-1; F: Serum level of GDF-15

**Fig. 14.**
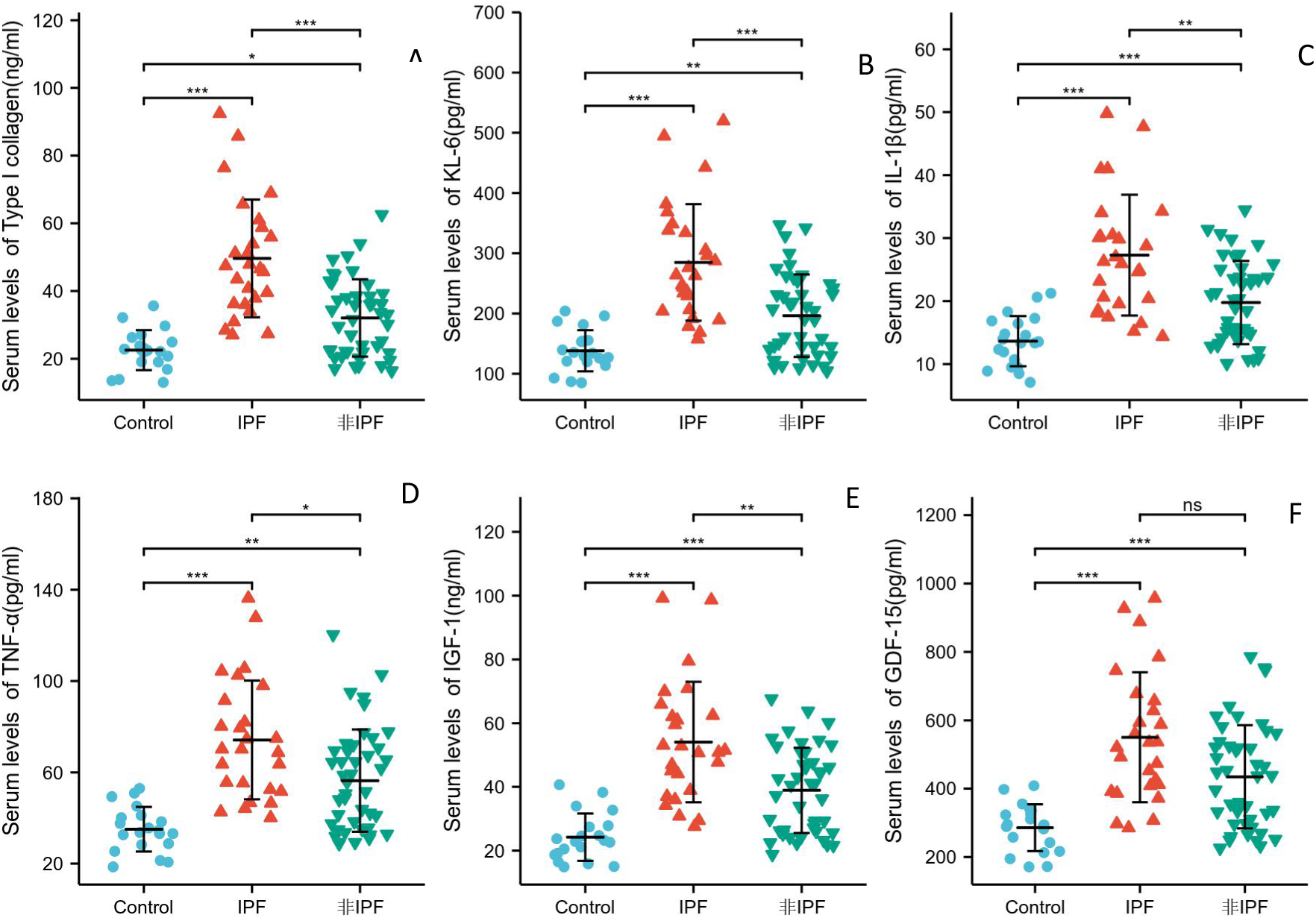
Comparison of serum detection indexes among IPF group, non-IPF group and healthy control group Note: NS: P ≥ 0.05; *: P< 0.05; **: P<0.01; * * *: P < 0.001, control: healthy control group; A: the serum level of type I collagen; B: serum level of KL-6; C: the serum level of IL-1β; D: Serum level of TNF-α; E: serum level of IGF-1; F: Serum level of GDF-15

**Table 9.**
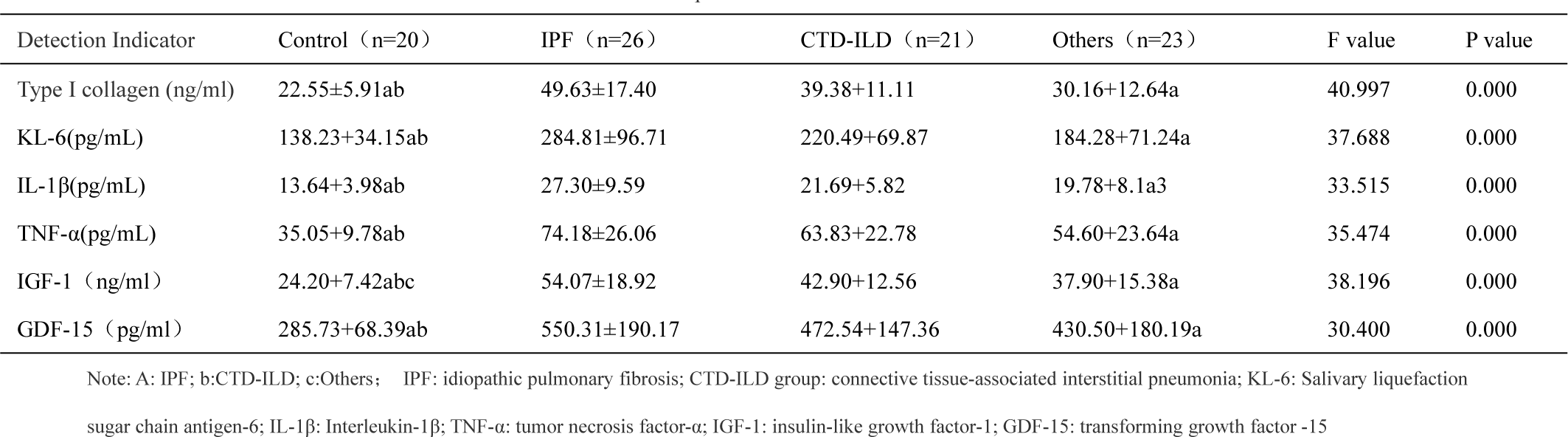
Comparison of detection indexes in serum

**Table 10.**
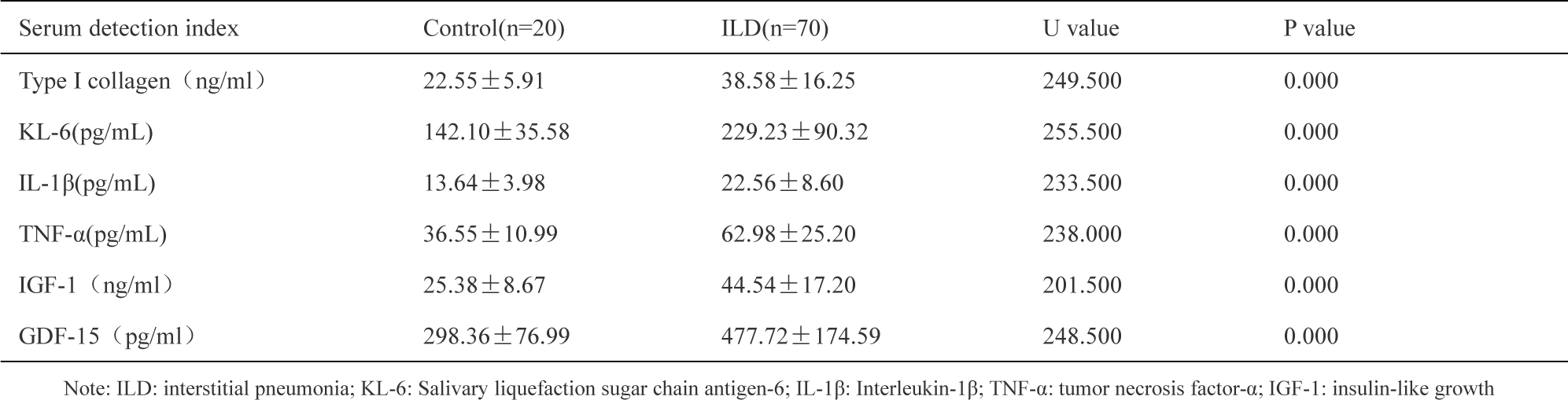
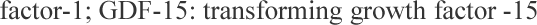
Comparison of serum detection indexes between ILD and healthy control group

| Serum detection index | Control(n=20) | ILD(n=70) | U value | P value |
| --- | --- | --- | --- | --- |
| Type I collagen (ng/ml) | 22.55 ± 5.91 | 38.58 ± 16.25 | 249.500 | 0.000 |
| KL-6(pg/mL) | 142.10 ± 35.58 | 229.23 ± 90.32 | 255.500 | 0.000 |
| IL-1β(pg/mL) | 13.64 ± 3.98 | 22.56 ± 8.60 | 233.500 | 0.000 |
| TNF-α(pg/mL) | 36.55 ± 10.99 | 62.98 ± 25.20 | 238.000 | 0.000 |
| IGF-1 (ng/ml) | 25.38 ± 8.67 | 44.54 ± 17.20 | 201.500 | 0.000 |
| GDF-15 (pg/ml) | 298.36 ± 76.99 | 477.72 ± 174.59 | 248.500 | 0.000 |

Note: ILD: interstitial pneumonia; KL-6: Salivary liquefaction sugar chain antigen-6; IL-1β: Interleukin-1β; TNF-α: tumor necrosis factor-α; IGF-1: insulin-like growth
factor-1; GDF-15: transforming growth factor -15

**Table 11.**
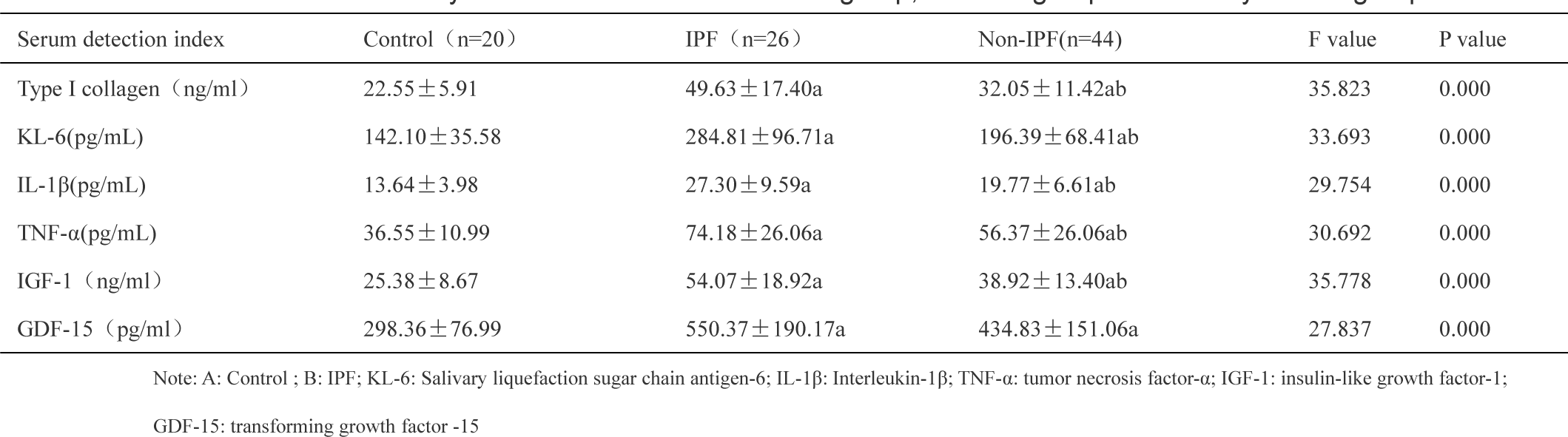
Analysis of detection indexes of IPF group, non-IPF group and healthy control group

### 2.3 The correlation between various detection indexes in serum and lung function

It can be seen from the analysis that the percentage of measured value/predicted value of GDF-15 and DLco is negatively correlated, and the difference is statistically significant. Type I collagen was negatively correlated with carbon monoxide dispersion, dispersion coefficient, FVC measured value/predicted value percentage, TLC measured value/predicted value percentage, VC measured value/predicted value percentage and DLco measured value/predicted value percentage, and the difference was statistically significant. KL-6 is negatively correlated with VC, DLco, DLco/VA, FVC/predicted percentage, TLC/predicted percentage, VC/predicted percentage and DLco/VA/predicted percentage, and the difference is statistically significant. IL-1β is negatively correlated with measured value of DLco, percentage of VC measured value/predicted value and percentage of measured value/predicted value of DLco, and the difference is statistically significant. IGF-1 is negatively correlated with FVC, VC, DLco, DLco/VA, FVC/predicted percentage, VC/predicted percentage and DLco/predicted percentage, and the difference is statistically significant. As shown in Table 15.

### 1.3 Correlation between GDF-15 and other serum detection indexes

GDF-15 was positively correlated with type I collagen, KL-6, interleukin −1β, tumor necrosis factor-α and insulin-like growth factor −1, and the difference was statistically significant (P < 0.05), and the positive correlation with TNF-α was the strongest. It can be seen from the table that KL-6 has the greatest correlation with type I collagen, which is statistically significant (P < 0.05). As shown in Table 13 and Figure 15.

**Fig. 15.**
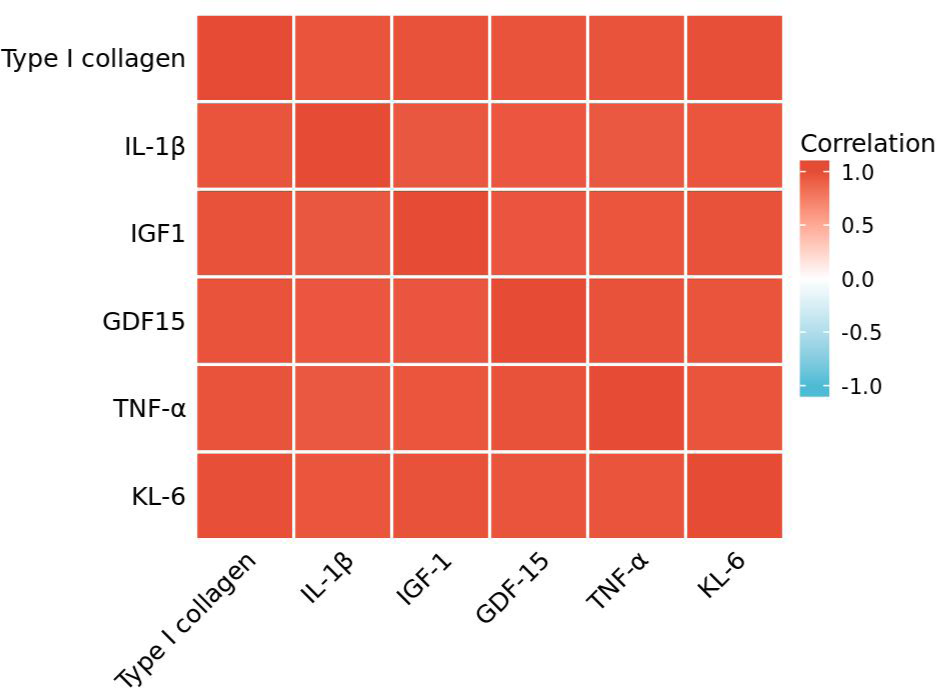
Correlation heat map of serum detection indexes Note: KL-6: Salivary liquefaction sugar chain antigen-6; IL-1β: Interleukin-1β; TNF-α: tumor necrosis factor-α; IGF-1: insulin-like growth factor-1; GDF-15: transforming growth factor −15

**Table 12.** Correlation between detection indexes in serum and lung function

| Pulmonary<br>function index<br>(n=40) | GDF-15 | | Type I collagen | | KL-6 | | IL-1 $\beta$ | | TNF- $\alpha$ | | IGF-1 | |
| --- | --- | --- | --- | --- | --- | --- | --- | --- | --- | --- | --- | --- |
|  | R | P | R | P | R | P | R | P | R | P | R | P |
| FVC measured value | -0.253 | 0.115 | -0.301 | 0.059 | -0.305 | 0.056 | -0.245 | 0.128 | -0.275 | 0.086 | -0.330 | 0.038 |
| TLC measured value | -0.015 | 0.755 | -0.113 | 0.486 | -0.106 | 0.516 | -0.057 | 0.728 | -0.097 | 0.551 | -0.170 | 0.295 |
| VC measured value | -0.239 | 0.138 | -0.293 | 0.066 | -0.320 | 0.044 | -0.254 | 0.113 | -0.262 | 0.103 | -0.335 | 0.035 |
| DLco measured value | -0.274 | 0.087 | -0.403 | 0.010 | -0.402 | 0.010 | -0.357 | 0.024 | -0.262 | 0.103 | -0.335 | 0.035 |
| DLco/VA measured value | -0.193 | 0.232 | -0.314 | 0.049 | -0.327 | 0.040 | -0.290 | 0.069 | -0.289 | 0.070 | -0.369 | 0.019 |
| FVC Measured value/predicted value percentage | -0.258 | 0.180 | -0.355 | 0.025 | -0.357 | 0.024 | -0.305 | 0.055 | -0.243 | 0.132 | -0.368 | 0.020 |
| TLC Measured value/predicted value percentage | -0.209 | 0.196 | -0.317 | 0.046 | -0.324 | 0.041 | -0.246 | 0.126 | -0.186 | 0.251 | -0.312 | 0.050 |
| VC Measured value/predicted value percentage | -0.277 | 0.083 | -0.346 | 0.029 | -0.374 | 0.017 | -0.333 | 0.036 | -0.252 | 0.116 | -0.362 | 0.022 |
| DLco Measured value/predicted value percentage | -0.320 | 0.044 | -0.428 | 0.006 | -0.422 | 0.007 | -0.401 | 0.010 | -0.306 | 0.055 | -0.388 | 0.013 |
| DLco/VA Measured value/predicted value percentage | -0.121 | 0.456 | -0.198 | 0.221 | -0.205 | 0.204 | -0.234 | 0.147 | -0.111 | 0.497 | -0.128 | 0.429 |

Note: KL-6: Salivary liquefaction sugar chain antigen-6; IL-1β : Interleukin-1β ; TNF-α : tumor necrosis factor-α ; IGF-1: insulin-like growth factor-1; GDF-15: transforming growth factor-15; FVC: vital capacity; TLC: total lung volume; VC: vital capacity, DLco: total diffusion capacity; DLco/VA: unit dispersion; R: correlation coefficient

**Table 13.** Correlation analysis of serum detection indexes

| Serum index | Type I collagen | | IL-1 $\beta$ | | IGF-1 | | GDF-15 | | TNF- $\alpha$ | | KL-6 | |
| --- | --- | --- | --- | --- | --- | --- | --- | --- | --- | --- | --- | --- |
|  | r | P | r | P | r | P | r | P | r | P | r | P |
| Type I collagen | 1 | 0.000 | 0.959 | 0.000 | 0.969 | 0.000 | 0.963 | 0.000 | 0.965 | 0.000 | 0.984 | 0.000 |
| IL-1β | 0.959 | 0.000 | 1 | 0.000 | 0.944 | 0.000 | 0.948 | 0.000 | 0.938 | 0.000 | 0.957 | 0.000 |
| IGF-1 | 0.969 | 0.000 | 0.944 | 0.000 | 1 | 0.000 | 0.953 | 0.000 | 0.956 | 0.000 | 0.969 | 0.000 |
| GDF-15 | 0.963 | 0.000 | 0.948 | 0.000 | 0.953 | 0.000 | 1 | 0.000 | 0.971 | 0.000 | 0.958 | 0.000 |
| TNF-α | 0.965 | 0.000 | 0.938 | 0.000 | 0.956 | 0.000 | 0.971 | 0.000 | 1 | 0.000 | 0.959 | 0.000 |
| KL-6 | 0.984 | 0.000 | 0.957 | 0.000 | 0.969 | 0.000 | 0.958 | 0.000 | 0.959 | 0.000 | 1 | 0.000 |

### 1.4 ROC curve of detection indexes in serum

Through ROC curve analysis, in IPF group, the areas under ROC curve of the six indexes detected in serum were all greater than 0.9, which had high diagnostic value. In CTD-ILD group, the AUC of IGF-1 was greater than 0.9, the cut-off value was 35.081ng/ml, the sensitivity was 90%, the specificity was 76%, and the area under ROC curve was greater than 0.8, and the difference was statistically significant. In other groups, the ROC curve area of these six indicators ranged from 0.6 to 0.8, among which IGF-1, GDF-15 and TNF-α had diagnostic value, while the other indicators had no diagnostic value. Figure 16.1 16.2 16.3, Table 14.1 14.2 14.3

**Fig. 16.1.**
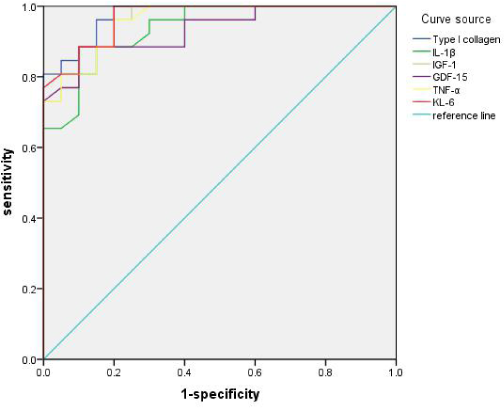
ROC curve of IPF and healthy control group

**Fig. 16.2.**
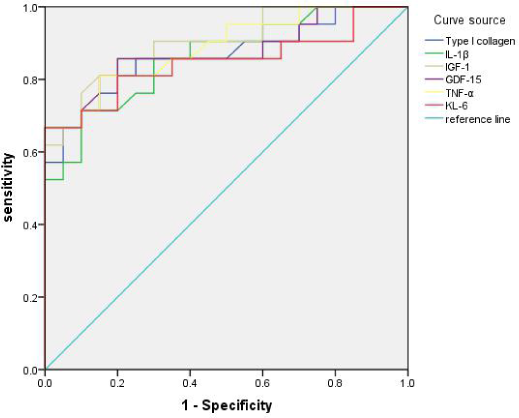
ROC curve of CTD-ILD group and healthy control group

**Fig. 16.3.**
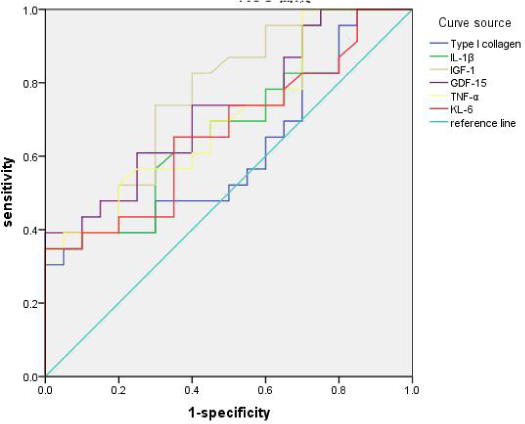
ROC curve of other groups and healthy control group

**Table 14.1.** ROC curve area of IPF group and healthy control group

| Serum detection index | AUC | Truncated value | sensitivity | Specificity | P value | Asymptotic 95% confidence interval |  |
| --- | --- | --- | --- | --- | --- | --- | --- |
|  |  |  |  |  |  | lower limit | upper limit |
| Type I collagen (ng/ml) | 0.975 | 27.247 | 0.85 | 0.96 | 0.000 | 0.940 | 1.000 |
| IL-1 $\beta$ (pg/mL) | 0.937 | 17.367 | 0.86 | 0.89 | 0.000 | 0.872 | 1.000 |
| IGF-1 (ng/ml) | 0.962 | 34.069 | 0.90 | 0.86 | 0.000 | 0.915 | 1.000 |
| GDF-15 (pg/ml) | 0.934 | 363.280 | 0.90 | 0.89 | 0.000 | 0.865 | 1.000 |
| TNF- $\alpha$ (pg/mL) | 0.959 | 41.860 | 0.80 | 0.96 | 0.000 | 0.910 | 1.000 |
| KL-6(pg/mL) | 0.968 | 156.529 | 0.80 | 1.00 | 0.000 | 0.928 | 1.000 |

**Table 14.2.** ROC curve of CTD-ILD group and healthy control group

| Serum detection index | AUC | Truncated value | sensitivity | Specificity | P value | Asymptotic 95% confidence interval |  |
| --- | --- | --- | --- | --- | --- | --- | --- |
|  |  |  |  |  |  | lower limit | upper limit |
| Type I collagen<br>(ng/ml) | 0.865 | 32.651 | 0.95 | 0.67 | 0.000 | 0.751 | 0.980 |
| IL-1 β (pg/mL) | 0.862 | 18.409 | 0.90 | 0.71 | 0.000 | 0.751 | 0.973 |
| IGF-1 (ng/ml) | 0.901 | 35.081 | 0.90 | 0.76 | 0.000 | 0.808 | 0.994 |
| GDF-15 (pg/ml) | 0.875 | 423.125 | 1.00 | 0.67 | 0.000 | 0.763 | 0.987 |
| TNF- α (pg/mL) | 0.888 | 54.267 | 1.00 | 0.67 | 0.000 | 0.789 | 0.988 |
| KL-6(pg/mL) | 0.848 | 204.629 | 1.00 | 0.67 | 0.000 | 0.721 | 0.974 |

**Table 14.3.** ROC curves of other ILD groups and healthy control groups

| Serum detection index | AUC | Truncated<br>value | sensitivity | Specificity | P value | Asymptotic 95%<br>confidence interval |  |
| --- | --- | --- | --- | --- | --- | --- | --- |
|  |  |  |  |  |  | lower<br>limit | upper limit |
| Type I collagen<br>(ng/ml) | 0.609 | 35.850 | 1.00 | 0.30 | 0.223 | 0.438 | 0.780 |
| IL-1 β (pg/mL) | 0.690 | 21.307 | 1.00 | 0.35 | 0.033 | 0.533 | 0.848 |
| IGF-1 (ng/ml) | 0.768 | 24.939 | 0.70 | 0.74 | 0.003 | 0.628 | 0.908 |
| GDF-15 (pg/ml) | 0.726 | 428.652 | 1.00 | 0.39 | 0.011 | 0.576 | 0.876 |
| TNF- α (pg/mL) | 0.702 | 55.837 | 1.00 | 0.35 | 0.024 | 0.547 | 0.857 |
| KL-6(pg/mL) | 0.665 | 208.058 | 1.00 | 0.35 | 0.064 | 0.503 | 0.827 |

Note: IPF: Idiopathic pulmonary fibrosis; CTD-ILD: connective tissue-associated interstitial pneumonia; ILD: interstitial pneumonia; KL-6:

### 1.5 Correlation between L- glutamic acid and serum detection indexes in idiopathic pulmonary fibrosis

In IPF, L- glutamic acid was positively correlated with type I collagen, KL-6, IGF-1, IL-1β, TNF-α and GDF-15, and the difference was statistically significant (P < 0.05), with the highest correlation with IGF-1. As shown in Table 15 and Figure 17

**Fig. 17.**
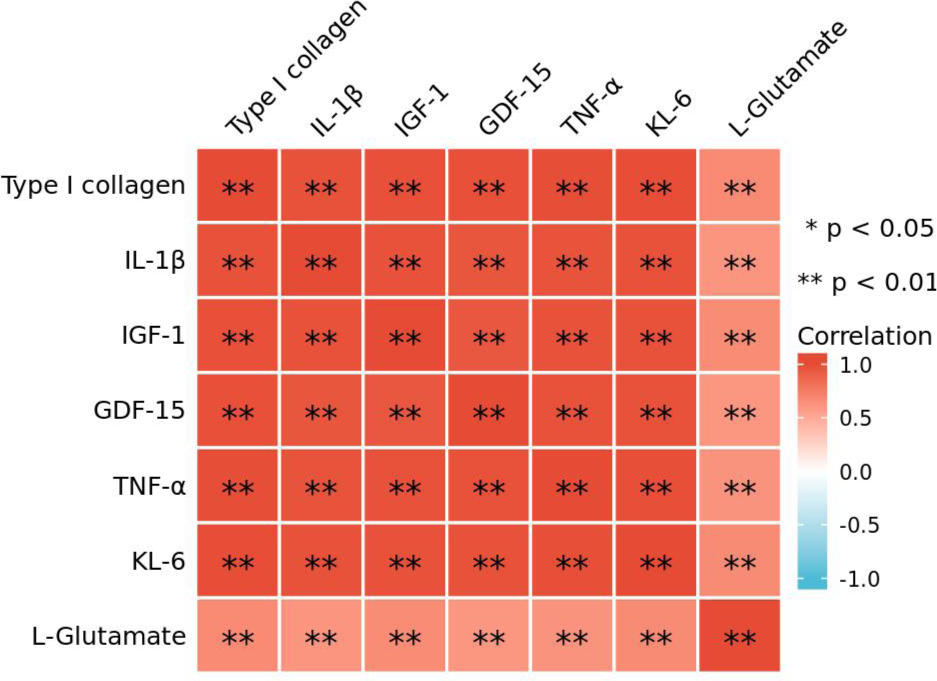
Correlation heat map of L-Glutamate and serum detection indexes in IPF group Note: IPF: Idiopathic pulmonary fibrosis; L-glutamic acid: L-glutamic acid; KL-6: Salivary liquefaction sugar chain antigen-6; IL-1β: Interleukin-1β; TNF-α: tumor necrosis factor-α; IGF-1: insulin-like growth factor-1; GDF-15: transforming growth factor-15; R: correlation coefficient

**Table 15.** Correlation analysis between L-glutamic acid and detection indexes in serum

| Detection Indicator | r | P |
| --- | --- | --- |
| Type I collagen (ng/ml) | 0.50 | 0.01 |
| KL-6(pg/mL) | 0.54 | 0.00 |
| IL-1 β (pg/mL) | 0.49 | 0.01 |
| TNF- α (pg/mL) | 0.44 | 0.02 |
| IGF-1 (ng/ml) | 0.55 | 0.00 |
| GDF-15 (pg/ml) | 0.44 | 0.02 |

## 3 Discussion

ILD mainly shows diffuse pulmonary lesions, which is a chronic disease caused by different degrees of inflammatory cell infiltration and fibrosis. The lesions involve alveolar walls and perialveolar tissues, mainly occurring in fibrous tissues, lymphatic vessels and blood vessels of pulmonary interstitium. There are many kinds of ILD, about 300 kinds, among which IPF and CTD-ILD are common, IPF has poor prognosis and low survival rate [33]. The clinical manifestations, treatment and prognosis of different diseases and subtypes of ILD are very different, so it is of great significance for the correct diagnosis of ILD.

The pathogenesis of IPF mainly includes the injury of alveolar epithelium, the destruction of lung tissue and the imbalance of regulation of aging genes. In the early stage of IPF, repeated damage and repair of alveolar epithelium are related to abnormal repair, and in this process, cytokine production, activation of various signaling pathways, epigenetics and family inheritance play an important role [34, 35]. These factors will lead to the release of pro-inflammatory and pro-fibrosis factors, induce the proliferation and activation of fibroblasts and myofibroblasts, and promote the synthesis and secretion of extracellular matrix, especially collagen fibers. The regulation mechanism of apoptosis is out of balance, and the growth factors and signal pathways are continuously activated abnormally, which makes the disease develop continuously [36]. Among them, transforming growth factor (TGF-β) is the most important fibrogenic cytokine, which plays a central role in various organ fibrosis. It can activate downstream signal pathways, activate fibroblasts and myofibroblasts, and inhibit metalloproteinases to reduce collagen decomposition, thus promoting the formation of fibrosis [37].

At present, the etiology of interstitial lung diseases is not clear, and the main causes are smoking, virus infection, environmental factors (dust, asbestos), genetic factors and gastroesophageal reflux [38]. German guidelines suggest that IPF patients are more likely to be complicated with cardiovascular diseases, lung cancer, depression, sleep-related respiratory diseases, and may be complicated with emphysema or chronic obstructive pulmonary diseases, which are closely related to smoking, and the prognosis of IPF is mainly pulmonary fibrosis. Then, the treatment of IPF mainly delays or reverses pulmonary fibrosis. On the premise of treatment, the diagnosis of IPF requires accuracy and timeliness. At present, the diagnosis of IPF is still exclusive, excluding known interstitial lung diseases, and differential diagnosis mainly includes CTD-ILD, pneumoconiosis, sarcoidosis, allergic pneumonia, etc. [20]. At present, we can explore more new biomarkers, provide more evidence for the diagnosis and differentiation of interstitial lung diseases, improve the diagnosis rate of diseases, take timely treatment measures, delay the process of pulmonary fibrosis and improve the survival rate.

At present, the diagnostic basis of ILD mainly includes clinical, imaging and pathological data. Multidisciplinary discussion is of great significance to the diagnosis of ILD, and it is the gold standard for the diagnosis of interstitial lung diseases [39]. The course of IPF may be slow progress, rapid progress, acute exacerbation and other types. At present, there is no recognized and effective forward-looking prediction method for the course of IPF and acute exacerbation. As for IPF treatment, anti-fibrosis therapy (pirfenidone and Nidanib) and lung transplantation are the mainstream at present, but there is also a lack of effective guidelines for individualized treatment of patients. Serum biomarkers are the focus of people’s attention, which can detect and diagnose diseases early, and evaluate the severity, treatment and prognosis of diseases. At present, the diagnosis of interstitial lung diseases mainly depends on HRCT and pathological biopsy, which are expensive and invasive. Compared with this, the serum biomarker measurement method is simple, rapid, non-invasive, less invasive and relatively cheap, and it can also be continuously detected to further evaluate the condition and prognosis. Therefore, it is of great significance to explore more new biomarkers and new biological targets.

Type I collagen is an important indicator of fibrosis. Studies show that type I collagen increases in silicosis fibrosis group [40]. KL-6 is an antigen on the surface of type II alveolar epithelial cells, a kind of high molecular mucin, also known as epithelial mucin 1. A study shows that KL-6 is of great significance for the diagnosis of idiopathic pulmonary fibers, and combined with SP-A, SP-D and MMP-7, it is sensitive for the early diagnosis of idiopathic pulmonary fibrosis. In a study of 142 patients with ILD, it was suggested that the concentration of KL-6 could distinguish IPF from IPF-LC, and the high concentration was correlated with the prognosis [41]. Interleukin −1β is a key pro-inflammatory cytokine, which plays an important role in cell proliferation, differentiation and apoptosis, and is also a fibrogenic factor. Recent studies have found that IL-1β is higher in acute exacerbation of idiopathic pulmonary fibrosis than in stable period, and it is an independent risk factor for death within 3 months [42]. Tumor necrosis factor-α is a multi-directional pro-inflammatory cytokine secreted by many inflammatory cells. In lung tissue, TNF-α can resist the repeated damage and repair of alveolar epithelium by promoting the apoptosis of type II alveolar epithelial cells, so it plays an important role in the formation of pulmonary fibrosis, and it can further promote the inflammatory response by recruiting neutrophils [43]. It is pointed out that TNF-α participates in the formation of pulmonary fibrosis by activating NF-κB signaling pathway, and it is found to be elevated in alveolar lavage fluid of patients with connective tissue-associated interstitial pneumonia, which is of great significance for the diagnosis of interstitial lung diseases [44]. IGF-1 is an active protein polypeptide molecular substance, a necessary substance in the process of growth hormone production, and an important factor to promote cell growth. IGF-1 signaling pathway is activated in the lung tissue of IPF patients, which can age alveolar epithelial cells and further cause pulmonary fibrosis. IGF-1/PI3K/AKT signaling pathway is activated in the lung tissue of IPF patients, and IGF-1 enhances core fucose glycosylation further in vitro and in vivo. GDF-15 is a branch member of transforming growth factor −β(TGF-β) superfamily. Related studies suggest that GDF-15 is closely related to metabolism [12]. Related studies show that GDF-15 plays an important role in energy homeostasis and weight regulation, and can be used as a therapeutic target for metabolic diseases [46]. A recent study shows that the increase of GDF-15 level in blood may precede the development of pulmonary fibrosis, and may mediate the relationship between aging and interstitial lung abnormalities, but further research is needed [13]. It has been suggested that GDF-15 may promote pulmonary fibrosis by activating macrophages and fibroblasts [14], but there is no clear research on the relationship between GDF-15 and interstitial lung diseases.

A total of 90 cases were included in this study, including 26 IPF patients, 21 CTD-ILD21 patients, 23 other ILDs and 20 healthy controls. By analyzing the basic clinical data of different groups, it was found that the carbon monoxide tolerance of IPF group was significantly lower than that of other interstitial lung diseases, and the diffusion coefficient of idiopathic pulmonary fibrosis group was lower than that of CTD-ILD group and other groups, as shown in Table 8. For these six indexes detected in serum, IPF group and CTD-ILD group are higher than the healthy control group, and the difference is statistically significant. For IGF-1, other groups are higher than the healthy control group, and the difference is statistically significant. And these six indexes, although IPF group is slightly higher than CTD-ILD group, have no significant difference. The difference between IPF group and other groups is statistically significant. Through analysis, it is found that these six indexes can not distinguish IPF from CTD-ILD, but can be used as markers to distinguish IPF from other interstitial lung diseases. Through the comparison of all ILD and healthy control group, it is found that these six indexes of serum detection are higher than those of healthy control group, and the difference is statistically significant (P < 0.05), as shown in Table 10 and Figure 13. Comparative analysis of IPF group, non-IPF group and healthy control group showed that the expression of GDF-15 in IPF group and non-IPF group was higher than that in healthy control group, and the difference was statistically significant (P < 0.05), but the difference was not statistically significant between IPF group and non-IPF group, and other serum indexes were significantly different between two groups. Through the correlation analysis between serum indexes and lung function, it was found that GDF-15 was negatively correlated with the percentage of measured value/predicted value of DLco, which was consistent with the results of an IPF plasma proteomics [47], while type I collagen was negatively correlated with carbon monoxide dispersion, dispersion coefficient, FVC%pred, TLC%pred, VC%pred and DLco%pred. KL-6 was negatively correlated with the measured values of VC, DLco, DLco/VA, FVC%pred, TLC%pred, VC%pred and DLco/VA%pred. IL-1β was negatively correlated with measured values of DLco, VC%pred and DLco%pred. IGF-1 is negatively correlated with FVC measured values, VC measured values, DLco measured values, DLco/VA measured values, FVC%pred, VC%pred and DLco%pred. As shown in Table 12, the indicators of lung function with diffusion dysfunction and restrictive ventilation dysfunction in previous studies are negatively correlated with KL-6 [48], which is consistent with the results of this study.

By analyzing the correlation between GDF-15 and other detection indexes, we can find that GDF-15 is positively correlated with type I collagen, KL-6, IL-1β, TNF-α and IGF-1, and the positive correlation with TNF-α is the strongest. Recently, it has been found that GDF-15 can inhibit the synthesis of myocardial fibroblasts and collagen under the condition of high glucose induction [49], and related literatures suggest that when TGF-β, IL-1β and TNF-α are activated by induction, the expression of GDF-15 is up-regulated [50], showing a positive correlation, which is consistent with the results of this study. In this study, the cut-off value, sensitivity and specificity of various detection indexes were obtained through the analysis of ROC curve. In IPF group, the areas under ROC curve of six detection indexes in serum were all greater than 0.9, which had high diagnostic value, among which the AUG of GDF-15 was 0.934, the cut-off value was 363.280pg/ml, the sensitivity was 90%, and the specificity was 89%. In the CTD-ILD group, the AUG of IGF-1 is greater than 0.9, the cutoff value is 35.081ng/ml, the sensitivity is 90%, the specificity is 76%, and the area under the ROC curve of the residual index is greater than 0.8, and the difference is statistically significant, among which the AUC of GDF-15 is 0.875, the sensitivity is 100%, the specificity is 69%. In other groups, the ROC curve area of these six indicators ranged from 0.6 to 0.8, among which IGF-1, GDF-15 and TNF-α had diagnostic value, while the other indicators had no diagnostic value. Therefore, it can be seen that in IPF, type I collagen, IL-1β, GDF-15, TNF-α and KL-6 are of high significance for the diagnosis of IPF, IGF-1 is of high diagnostic value in IPF and CTD-ILD, while type I collagen, IL-1β and KL-6 have no diagnostic value in other groups, so they may be used to differentiate IPF.

For IPF, on the basis of serum metabonomics, it was found that L- glutamic acid has high diagnostic value, with AUG of 0.921, sensitivity of 96% and specificity of 80%. Through correlation analysis with other six indexes, it was found that there was a positive correlation, and the correlation coefficient with GDF-15 was 0.44. GDF-15 may prevent the excessive activation of fibrotic cells in the process of lung tissue remodeling through TGF-β pathway [52], GDF-15 is a cytokine that regulates energy metabolism, AMPK-p53 pathway is involved in PPARb/d-mediated increase of GDF-15, and the up-regulation of GDF-15 is mediated by MAPK/ERK activation [53, 54] A study on cervical cancer shows that GDF-15 down-regulates the expression of p21 through PI3K/AKT and MAPK/ERK signaling pathways, and promotes cell proliferation [55]. In the metabonomics of this study, L- glutamic acid is involved in the metabolic pathway, and it is an up-regulator. In this pathway, L- glutamic acid is involved in MAPK pathway, L- glutamic acid is positively correlated with GDF-15, and GDF-15 has a great relationship with metabolism. MAPK pathway can regulate Smad pathway, which is a signal transduction pathway that inhibits cell proliferation. Studies show that TGF-β1 promotes the formation of pulmonary fibrosis through Smad pathway [56], GDF-15 induces HSC, and increases the phosphorylation of SMAD2 and SMADA3, which plays an important role in the formation of pulmonary fibrosis [57]. Therefore, in IPF, GDF-15 may be involved in MAPK signaling pathway to activate Smad pathway, and further activate related fibrogenic factors, thus promoting pulmonary fibrosis. However, which link GDF-15 specifically participates in MAPK signaling pathway activation still needs to be verified by experiments.

Due to the limited time and funds, the sample size of this study is still insufficient, and a larger sample size is still needed. At the same time, a confirmatory experiment is also needed to further clarify the metabolic pathway of GDF-15. Due to the small number of samples for patients to improve their lung function, the results may be biased and related to the degree of cooperation of patients, which may affect the accuracy of the data. Due to the low follow-up rate of patients, continuous monitoring is not possible, and the progress and prognosis of the disease cannot be evaluated. At present, whether type I collagen, KL-6, IL-1β, TNF-α, IGF-1 and GDF-15 can be used as diagnostic and differential diagnostic markers of ILD still needs a large number of prospective studies.

## 4 Conclusion

1. The expressions of type I collagen, KL-6, IL-1β, TNF-α, IGF-1 and GDF-15 are different in different interstitial lung diseases and correlated with lung function; GDF-15 is positively correlated with other indicators;
2. Type I collagen, KL-6, IL-1β, TNF-α, IGF-1 and GDF-15 have good diagnostic efficacy in IPF and CTD-ILD, and can be distinguished from other interstitial lung diseases;
3. Comparative analysis between ILD and healthy controls showed that the expression levels of type I collagen, KL-6, IL-1β, TNF-α, IGF-1 and GDF-15 in serum were higher than those in healthy controls, which indicated that these biomarkers were involved in the pathogenesis of ILD. Compared with IPF, non-IPF and healthy control group, the expression of GDF-15 in serum has no difference between IPF and non-IPF. 4. GDF-15 is positively correlated with L- glutamic acid in IPF, which may be involved in metabolic pathway and further promote pulmonary fibrosis.

